# OSCA: a tool for omic-data-based complex trait analysis

**DOI:** 10.1101/445163

**Authors:** Futao Zhang, Wenhan Chen, Zhihong Zhu, Qian Zhang, Marta F. Nabais, Ting Qi, Ian J. Deary, Naomi R. Wray, Peter M. Visscher, Allan F. McRae, Jian Yang

## Abstract

The rapid increase of omic data in the past decades has greatly facilitated the investigation of associations between omic profiles such as DNA methylation (DNAm) and complex traits in large cohorts. Here, we proposed a mixed-linear-model-based method (called MOMENT) that tests for association between a DNAm probe and trait with all other distal probes fitted in multiple random-effect components to account for the effects of unobserved confounders as well as the correlations between distal probes induced by the confounders. We demonstrated by simulations that MOMENT showed a lower false positive rate and more robustness than existing methods. MOMENT has been implemented in a versatile software package (called OSCA) together with a number of other implementations for omic-data-based analysis including the estimation of variance in a trait captured by all measures of multiple omic profiles, omic-data-based quantitative trait locus (xQTL) analysis, and meta-analysis of xQTL data.

## Introduction

The rapid proliferation of genetic and omic data in large cohort-based samples in the past decade have greatly advanced our understanding of the genetic architecture of omic profiles and the molecular mechanisms underpinning the genetic variation of human complex traits [1–3]. These advances include the identification of a large number of genetic variants associated with gene expression [4, 5], DNA methylation [6, 7], histone modification [8, 9], and protein abundance [10, 11]; the discovery of omic measures associated with complex traits [12, 13]; the improved accuracy in predicting a trait using omic data [14, 15]; and the prioritization of gene targets for complex traits by integrating genetic and omic data in large samples [3, 13, 16–18]. These advances have also led to the development of software tools, focusing on a range of different aspects of omic data analysis. Therefore, a software tool that implements reliable and robust statistical methods for comprehensive analysis of omic data with high-performance computing efficiency is required.

A well-recognised challenge in omic-data-based analysis is to control for false positive rate (FPR) in the presence of confounding factors, as failing to model the confounders may lead to spurious associations [19–21] and/or a loss of statistical power [22]. While some confounders (e.g., age and sex) are known and available in most data so that their effects can be accounted for by fitting them as covariates in linear models, others are either uncharacterised or difficult to measure. For example, in DNA methylation (DNAm) data from whole blood, cell type compositions (CTCs) are evident confounders in a methylome-wide association study (MWAS; also known as an epigenome-wide association study or EWAS) [21, 23, 24] although CTCs may be useful for the prediction of some phenotypes. CTCs tend to be correlated with the DNAm at CpG sites that are differentially methylated in different cell types (namely differentially methylated sites) and have been shown to be associated with age and multiple traits and diseases [19, 21, 25, 26]. MWAS analysis without accounting for CTCs could give rise to biased test-statistics unless neither CTCs nor DNAm sites are associated with the trait in question. Although it is possible to measure CTCs directly or predict them by reference-based prediction methods [27, 28], reference-free methods that are able to correct for confounding effects without the need of characterizing all the confounders have broader applications [22, 29–32]. Moreover, the predicted CTCs often only explain a certain proportion of variation in CTCs resulting in biased test-statistics due to the uncaptured variation in CTCs. Existing reference-free methods are mainly based on the strategy of fitting a number of covariates (estimated from factor analysis or similar approaches with or without reference [22, 29, 31, 32]) in a fixed-effect model or a set of selected DNAm probes in a mixed linear model (MLM) [30]. However, uncharacterized confounders with small to moderate effects and numerous correlations between distal DNAm probes (e.g. those on different chromosomes) induced by the confounders may not be well captured by either a fixed number of principal features or a subset of selected probes.

In this study, we proposed a reference-free method (called MOA: <u>M</u>LM-based <u>o</u>mic <u>a</u>ssociation) that fits all probes as random effects in an MLM-based association analysis to account for the confounding effects, including the correlations among distal probes induced by the confounding. We then extended the method to stratify the probes into multiple random-effect components (called MOMENT: <u>m</u>ulti-c<u>o</u>mponent <u>M</u>LM-based omic association <u>e</u>xcludi<u>n</u>g the target) to model a scenario where some probes are much more strongly associated with the phenotype than others. We evaluated the performance of MOA and MOMENT by extensive simulations and demonstrated their reliability and robustness in comparison with existing methods. We have implemented MOA and MOMENT together with a comprehensive set of other methods for omic data analysis in an easy-to-use and computationally efficient software package, OSCA (<u>o</u>mic-data-ba<u>s</u>ed <u>c</u>omplex trait <u>a</u>nalysis).

## Results

### Overview of the OSCA software

OSCA comprises four main modules: 1) data management for which we designed a binary format to efficiently store and manage omic data; 2) linear-regression-and MLM-based methods (including the methods proposed in this study) to test for associations between omic measures and complex traits; 3) methods to estimate the proportion of variance in a complex trait captured by all the measures of one or multiple omic profiles (e.g., all SNPs and DNAm probes), and to predict the trait phenotype in a new sample based on the joint effects of all omic measures estimated in a discovery sample; and 4) an efficient implementation of the methods to identify genetic variants associated with an omic profile, e.g., DNA methylation quantitative trait loci (mQTL) analysis. We will describe the methods based on DNAm data but the methods and software tool are in principle applicable to other types of omic data including gene expression, histone modification, and protein abundance. The computer code of OSCA is written in C++ programming language and supports multi-threading based on OpenMP for high-performance computing. The compiled binary files are freely available at http://cnsgenomics.com/software/osca/.

### MLM-based omic association analysis methods

One of the primary applications of OSCA is to test for associations between omic measures (e.g., DNAm probes) and a complex trait (e.g., body mass index (BMI)) correcting for confounding effects. In an MWAS, the test-statistics of null probes can be inflated because of the associations of probes with confounders that are correlated with the phenotype. Note that, even if the confounders are not directly associated with the phenotype, the presence of confounders (e.g., CTCs or experimental batches) can cause correlations between the trait-associated probes and the null probes in distal genomic regions or even on different chromosomes, giving rise to inflated test-statistics of the null probes (see the simulation results below). Existing methods that fit a number of covariates computed from dimension reduction approaches in a fixed-effect model [22, 31, 32] or a set of selected DNAm probes in an MLM [30] may not be sufficient to correct for confounding effects widely spread among a large number of probes or correlations between distal probes induced by the confounding. We propose two MLM-based approaches (MOA and MOMENT) that include all the (distal) probes as random effects in the model to account for the effects of the confounders on the trait and probes as well as the correlations among distal probes. We show by simulations (see below) that both MOA and MOMENT are more robust than existing methods in controlling for false positive rate (FPR) and family-wise error rate (FWER) in MWAS (see below).

Here we start with a general MLM that fits all probes as random effects, i.e.,

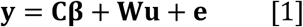

where **y** is an *n* × 1 vector of phenotype values with *n* being the sample size, **C** is an *n* × *p* matrix for covariates (e.g., age and sex) with *p* being the number of covariates, **β** is a *p* × 1 vector of the effects of covariates on the phenotype, **W** is an *n* × *m* matrix of standardised DNAm measures of all *m* probes, **u** is an *m* × 1 vector of the joint effects of all probes on the phenotype, and **e** is an *n* × 1 vector of residuals. In this model, **β** are fixed effects whereas **u** and **e** are random effects with 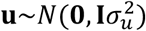 and 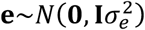 The variance-covariance matrix for **y** is 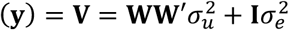. This equation can be re-written as

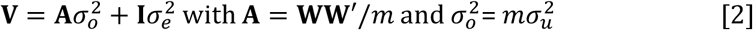

where **A** is defined as the omic-data-based relationship matrix (ORM) (**Methods**) and 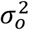 is the amount of phenotypic variance captured by all probes. The variance components (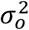 and 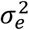) in such an MLM can be estimated by REML algorithms [33]. Analogous to the method for estimating SNP-based heritability [34, 35], the proportion of variance in the phenotype captured by all the probes can be defined as 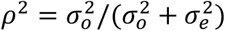. We name this variance-estimation method OREML following the nomenclature of GREML [34]. The estimated joint probe effects (**û**) from this model by a random-effect estimation approach (e.g., BLUP [36]) can be used to predict the phenotypes of individuals in a new sample based on omic data, i.e., **ŷ**_new_ = **W**_new_**û**. We call this OBLUP.

Model [1] can be extended to test for association between a probe *i* and the trait, i.e.,

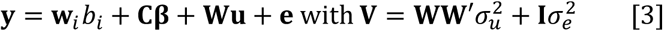

In comparison to model [1], this model has two additional terms, **w**_*i*_ (an *n* × 1 vector of standardised DNAm measures of a probe *i*, i.e., the target probe) and *b*_*i*_ (the effect of probe *i* on the phenotype; fixed effect). The probe effect *b*_*i*_ (together with the covariates’ effects) can be estimated by the generalized least squares (GLS) approach, i.e., 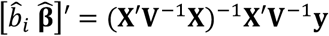 and var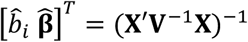 with **X** = [**w**_*i*_ **C**]. The sampling variance (standard error (SE) squared) of 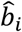 is the first diagonal element of var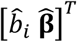 The null hypothesis (*H*_*0*_: *b*_*i*_= 0) can be tested by a two-sided *t*-test (or approximately chi-squared test if sample size is large) given 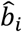 and its SE. We call this method MOA. Applying this method to test each of the probes across the genome is extremely computationally expensive because the variance components 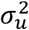 and 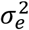 need to be estimated repeatedly for each probe by REML that requires the computation of **V**^−1^ (computational complexity of *O*(*n*^*3*^)) multiple times in an iterative process. To speed up the computation, we use a two-step approach as in [37] to compute **V**^−1^, with the first step to perform an eigendecomposition of **WW***’* and the second step to compute **V**^−1^ based on the eigenvalues and eigenvectors. Since the eigendecomposition only needs to be done once for the whole genome scan, this two-step approach reduces the complexity of computing **V**^−1^ by orders of magnitude when testing each specific probe. Moreover, as the proportion of phenotypic variance attributable to a single probe is often very small, we can further speed up the computation by an approximate approach (similar to the approximate MLM-based GWAS methods [38, 39]) that only requires to compute **V**^−1^ once, assuming that the estimates of 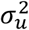 and 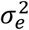 under the null (i.e., *b*_*i*_ = 0) are approximately equal to those under the alternative (i.e., *b*_*i*_ ≠ 0). Both the approximate and exact MOA approaches have been implemented in OSCA.

There are two properties of the MOA method worthy of consideration. First, the target probe is fitted twice in the MOA model, once as a fixed effect (*b*_*I*_) and again as a random effect (the *i*-th element of **u**), resulting in a loss of power to detect *b*_*I*_ (a recognised issue in MLM-based association analysis with SNP data [39, 40]). This problem can be solved by leaving out probes in close physical proximity of the target probe (including the target) from the random-effect term because DNAm status of CpG sites in close physical proximity are likely to be regulated by the same mechanism and therefore tend to be highly correlated. This strategy has been used previously in both GWAS (genome-wide association study) [39, 40] and MWAS [30]. In practice, we exclude the probes <50Kb from the target probe. Note that the distance parameter may differ for other types of omic data (e.g., a window size of 100Kbp is recommended for gene expression data; see below for details). Second, MOA assumes a single distribution to all the probe effects in the random-effect term, which may not be well fitted to data if some probes have much stronger associations with the trait than other probes. For example, if CTCs are associated with the phenotype, then all the probes that are highly differentially methylated in different cell types [41–43] may present a very different distribution of effects from the other probes. One solution to this issue is to stratify the probes into multiple groups by the association test statistics (from linear regression) and fit them as separate random-effect terms in the model. We extended the MOA method with the two modifications mentioned above and named it as MOMENT (<u>m</u>ulti-c<u>o</u>mponent <u>M</u>LM-based omic association <u>e</u>xcludi<u>n</u>g the target). The MOMENT model can be written as

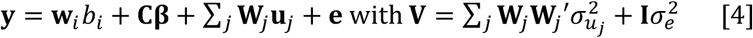

where **W**_*j*_ is an *n* × *m*_*j*_ matrix of standardised DNAm measures of the probes in the *j*-th group with *m*_*j*_ being the number of probes in the group (excluding probes within 50Kb of the target probe). In practice, the probes are split into two groups by association p-values from a linear regression model (i.e., **y** = **w**_*i*_ *b*_*i*_ + **Cβ** + **e**) at a methylome-wide significant threshold (all the methylome-wide significant probes in the first group and the other probes in the second group). The GLS method described in model [3] can be used to estimate *b*_*I*_ and its SE for hypothesis testing. Like the exact MOA method, MOMENT is also computationally intensive when applied in a methylome-wide analysis. We can use a similar approximation approach as described above (i.e., using the variance components estimated under the null to compute 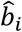 and SE) to reduce the computing cost. The variance components are re-estimated when one or more probes are excluded from the first group in case that the proportion of phenotypic variance captured by some of the probes in the first group are large.

### Simulation analysis

To quantify the false positive rate (or family-wise error rate) and statistical power of MOMENT (implemented in OSCA), we performed simulations based on DNAm and CTCs [44] measures on samples from the Lothian Birth Cohorts (LBC) in three scenarios (**Supplementary Note 1**). We simulated a phenotype 1) with effects from a set of “causal probes” (randomly selected from all probes on the odd chromosomes) but no direct effects from the CTCs; 2) with small to large effects from CTCs but no effects from the probes; and 3) with effects from both the causal probes and CTCs (**Supplementary Note 1**). Note that we only sampled the causal probes from the odd chromosomes in scenarios 1 and 3, leaving the probes on the even chromosomes to quantify false positive rate under the null, and that the DNAm measures were adjusted for age, sex, experimental batches, and smoking status. Results from our models were compared to 6 different methods including: 1) Unadj: linear regression without adjustment; 2) CTCadj: linear regression with CTCs fitted as covariates. 3) SVA: linear regression with the SVA surrogate variables fitted as covariates [22]; 4) LFMM2-ridge: a latent factor mixed model (LFMM) using ridge algorithm for confounder estimation [32]; 5) LFMM2-lasso: an LFMM using lasso algorithm for confounder estimation [32]; 6) ReFACTor: linear regression with the first 5 sparse principal components (PCs) from ReFACTor fitted as covariates [31]; 7) 5PCs: linear regression with the first 5 PCs, computed from a principal component analysis (PCA), fitted as covariates; and 8) FaST-LMM-EWASher: a set of selected probes fitted as random effect in an MLM [30]. For completeness of the analysis, we also included MOA (implemented in OSCA) in the comparison. We validated using a subset of data generated from simulation scenario 1 that the test-statistics from the approximate MOA/MOMENT approach were extremely highly correlated with those from the corresponding exact approach (Pearson correlation >0.999 for causal probes and >0.998 for null probes; **Supplementary Figure 1**). Hence, for the ease of computation, we used the approximate MOA/MOMENT approach in all the subsequent analyses.

In simulation scenario 1, although there were no direct effects of the CTCs on the phenotype, the test-statistics from Unadj at the null probes were inflated (**Figure 1a** and **Supplementary Table 1**) because the null and causal probes – albeit on different sets of chromosomes – are correlated through their correlations with systematic biases such as CTCs. The mean genomic inflation factor (*λ*) [45] of the null probes (on the even chromosomes) from 100 simulation replicates was 7.67 for Unadj (**Supplementary Table 1**), where *λ* is defined as the median of *χ*^2^ test-statistics of the null probes divided by its expected value. CTCadj reduced but not completely removed the inflation in test-statistics of the null probes (**Figure 1a** and **Supplementary Table 1**), suggesting that the inflation was caused by correlations between the causal and null probes because of the confounding effects of both CTCs and other unobserved confounders. While all the other methods were much less inflated compared to Unadj, MOMENT and MOA showed the least inflation with a mean *λ* value close to 1. It is slightly surprising to observe that the family-wise error rates (FWERs) of all the methods except MOA and MOMENT were highly inflated (FWERs > 0.6) (**Supplementary Figure 2a** and **Supplementary Table 1**) despite the relatively small genomic inflation at the null probes for most of the methods (**Figure 1a**). Here, FWER is defined as the proportion of simulation replicates with at least one null probe at MWAS p-value < 0.05 / *m* with *m* being the number of null probes, which can be interpreted as the probability of observing one or more false positives at a methylome-wide significance level in a single experiment. There was no inflation in FWER for MOMENT, and a marginal inflation for MOA (**Supplementary Figure 2a** and **Supplementary Table 1**), showing the effectiveness of using all (distal) probes to account for the probe correlations. We also quantified the FPR, defined as the proportion of null probes with p-values < 0.05 in each simulation replicate. The differences in FPR among the methods showed a similar pattern to the differences in genomic inflation factor (**Supplementary Figure 2b** and **Supplementary Table 1**). We then compared power among the methods. Since the test-statistics of many approaches were highly inflated, it is not very meaningful to compare power without accounting for the inflation. We therefore used the Area Under the ROC Curve (AUC) to compare power of the methods given the same level of FPR. Apart from Unadj and CTC, the AUCs of all the methods were on similar levels (**Figure 1b**). The conclusions held in additional simulations varying the number of causal probes and the proportion of phenotypic variance captured by the causal probes (**Supplementary Figures 3** and **4**) despite that the inflation in FWER for the existing methods appeared to increase with the increase of the proportion of variance captured per causal probe. Additionally, we applied BACON, a summary-data based method that seeks to remove genomic inflation taking the true positives into consideration, to the test-statistics of all probes produced by the methods tested above. We showed that the inflation in test-statistics of the null probes for Unadj was substantially reduced but not completely removed by the BACON adjustment and that the test-statistics from MOA and MOMENT remained almost unchanged after the BACON adjustment (**Supplementary Figure 5**).

**Figure 1.**
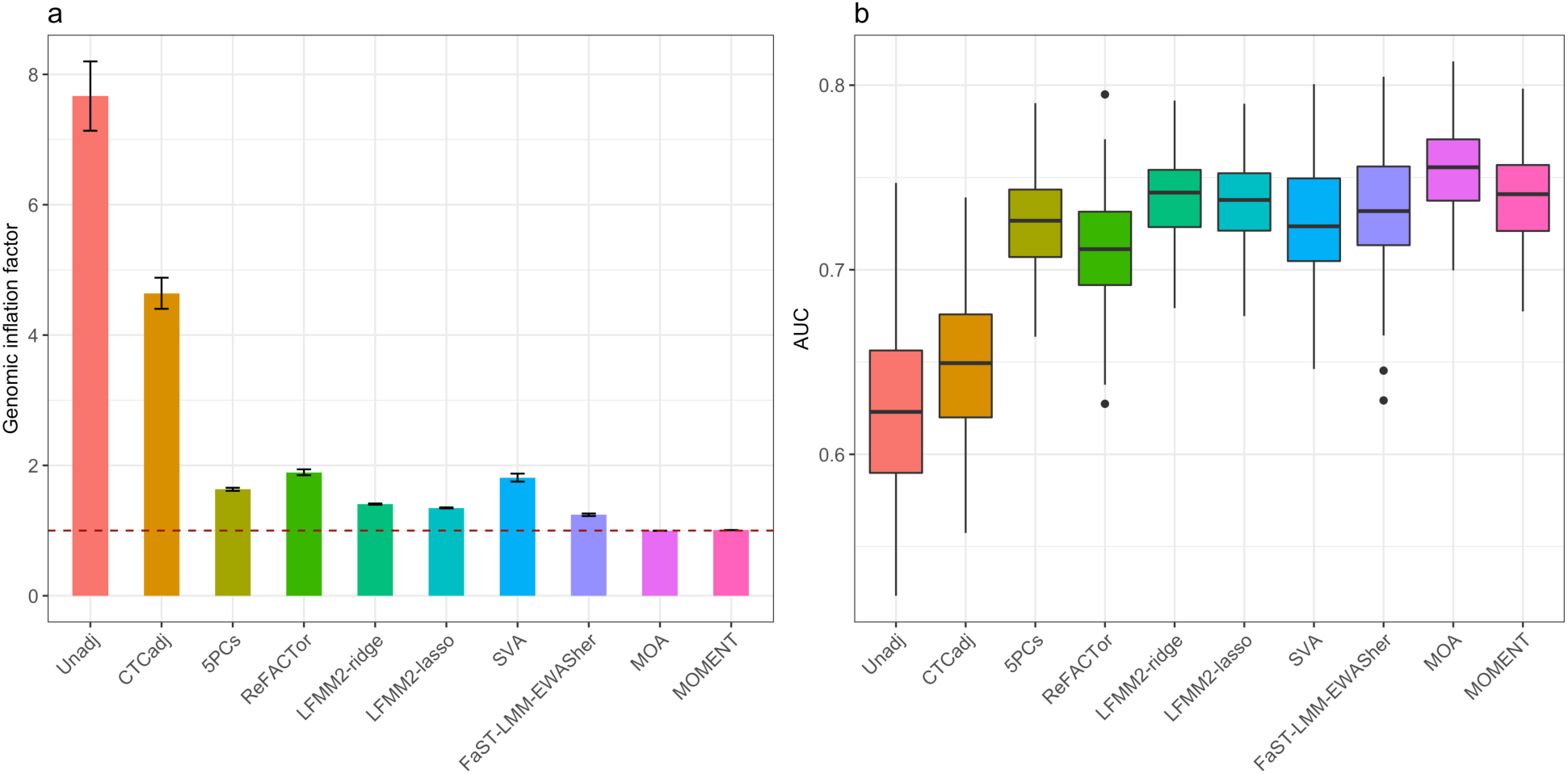
Power and false positive rate for the MWAS methods in simulation scenario 1. The phenotypes were simulated based on the effects from 100 causal probes but no direct effects from the CTCs. (a) Mean genomic inflation factor from a method across 100 simulation replicates with an error bar representing +/- SE of the mean. The dashed line at 1 shows the expected value if there is no inflation. (b) Box plot of AUCs for each method from 100 simulation replicates.

In simulation scenario 2 where there is no direct probe-trait association, all the probes are null and their *χ*^2^ test-statistics are expected to follow a *χ*^2^ distribution with 1 degree of freedom if the effects of CTCs have been well accounted for. The results showed that the *λ* value was close to 1 for all the methods except Unadj and FaST-LMM-EWASher (**Figure 2a**). It seems that, for some of the methods (e.g. 5PCs and ReFACTor), the *λ* value slightly increased with the increase of the proportion of variance explained by the CTCs 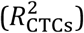 (**Figure 2a**). The FPRs of the methods were highly consistent with the genomic inflation factors (**Supplementary Figure 6**). Nevertheless, a non-inflated median test-statistic does not necessarily mean that the FWER has been well controlled for. In fact, most methods showed inflated FWER in this simulation scenario, and the FWERs of all the methods increased with increasing 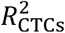 (**Figure 2b**). The FWERs of 5PCs, ReFACTor, LFMM2-ridge, and LFMM2-lasso were close to the expected value (i.e., 0.05) when 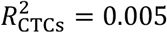 and increased to a level between 0.15 and 0.2 when 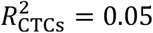 (**Figure 2b**). The relationship between FWER and 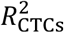 was relatively flat for SVA with its FWER varying from 0.05 to 0.1 when 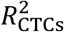 increased from 0.005 to 0.05. Although FaST-LMM-EWASher showed the most deflated test-statistics among all the methods (**Figure 2a**), its FWER was substantially higher than all the other methods except Unadj (**Figure 2b**), likely due to its feature selection strategy (**Supplementary Note 2**). MOA and MOMENT performed similarly in this simulation scenario and showed the lowest inflation in FWER among all the methods with their FWER being lower than 0.05 when 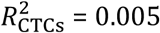 and increased to about 0.1 when 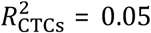 (**Figure 2b**). In addition, we performed a linear regression analysis with the known CTCs fitted as covariates; as expected, the FWER of was close to 0.05 irrespective of the level of 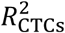 (see below for the analysis with predicted CTCs).

**Figure 2.**
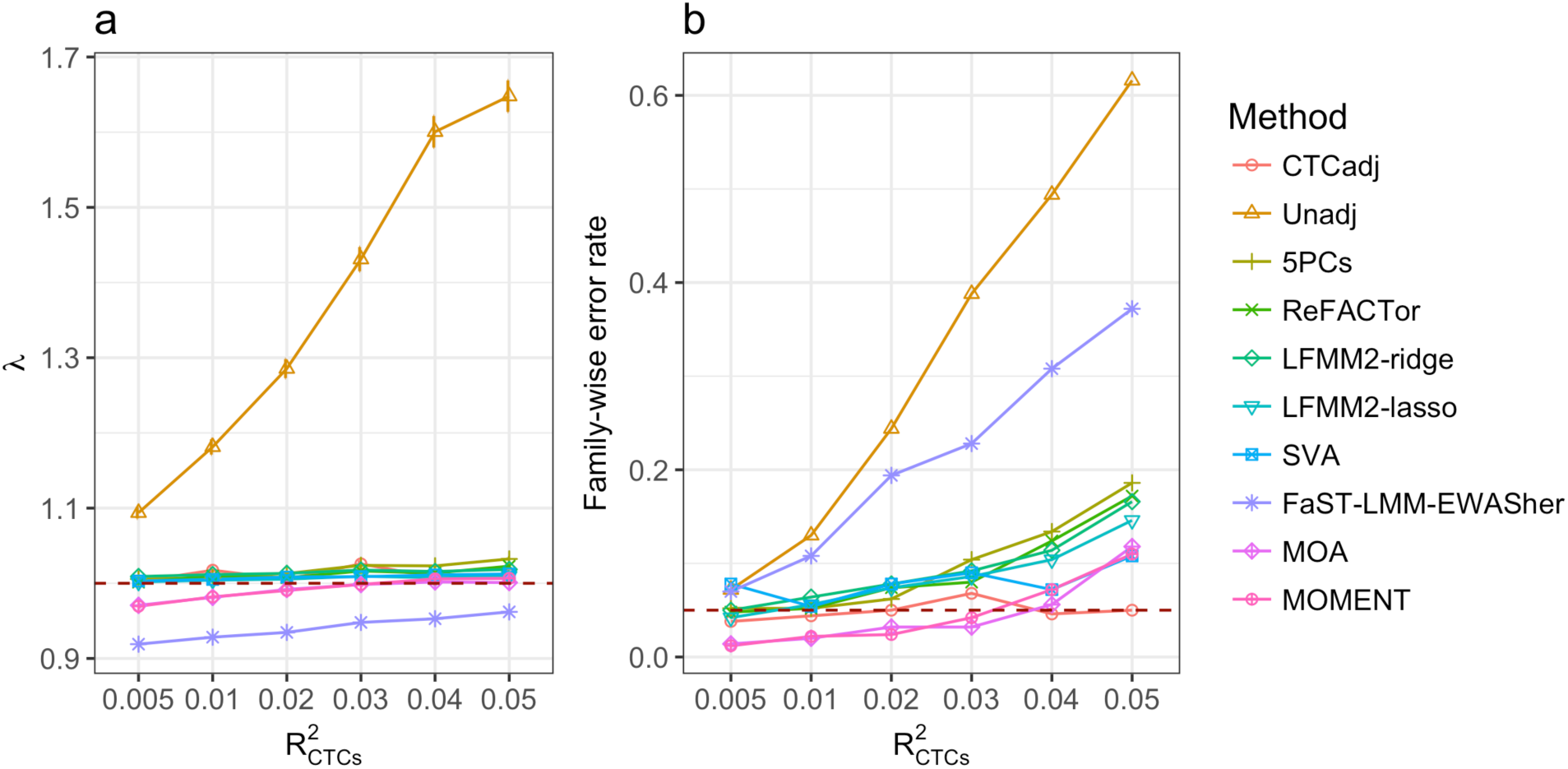
Genomic inflation factor and family-wise error rate for the MWAS methods in simulation scenario 2 (effects from CTCs but no causal probes). Shown on the horizontal axis are the 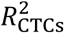 values used to simulate the phenotype. (a) Each dot represents the mean *λ* value from 1000 simulation replicates given a specified 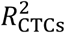 value for a method with an error bar representing +/- the SE of the mean. (b) Each dot represents the family-wise error rate, calculated as the proportion of simulation replicates with one or more null probes detected at a methylome-wide significance level.

We also compared the methods under the circumstance (simulation scenario 3) where there were associations between the phenotype and CTCs 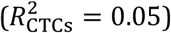 and the null probes were correlated with distal causal probes because both of them were correlated with CTCs (**Supplementary Note 1**). The results were similar to those above (**Figure 1** and **Supplementary Figure 2**). That is, the FWER of MOMENT was close to the expected value, demonstrating the reliability and robustness of the method. The FWER of MOA is slightly higher than that of MOMENT but much lower than those of the other methods which showed strong inflation in FWER and/or FPR due to the correlations between causal and null probes (**Supplementary Figures 7a, 7c** and **7d**, and **Supplementary Table 2**). All the methods showed similar levels of AUC except for Unadj and CTCadj (**Supplementary Figure 7b**). The conclusions held with different sample sizes (**Supplementary Figures 8** and **9**) or different numbers of causal probes with smaller or larger variance explained per causal probe (**Supplementary Figures 10** and **11**). The conclusions also held if we simulated confounding effects on experimental batches in lieu of CTCs (**Supplementary Figures 12** and **13**). We further demonstrated that the result from MOA/MOMENT analysis of the whole sample was consistent with that from a meta-analysis of summary statistics from MOA/MOMENT analyses in two halves of the sample (**Supplementary Figure 14**) and that the methods were applicable to case-control phenotypes (**Supplementary Figures 15** and **16**).

To explore the applicability of the proposed methods to other types of omic data, we tested the methods by simulation based on a real gene expression data set (19,648 gene expression probes on 1,219 Mexican American individuals) from the San Antonio Family Heart Study (SAFHS) [46–48] (**Methods**) under simulation scenario 1 (i.e., quantitative phenotypes simulated based on the expression levels of 100 randomly selected causal probes; **Supplementary Note 1**). The result showed that both MOMENT and MOA performed similarly (in comparison to the other methods) as in the simulations based on DNAm data (**Supplementary Figure 17**).

We further compared the computational complexity among the MWAS methods tested in this study and quantified their runtime and memory usage of the methods using simulated and real phenotypes in the LBC (**Supplementary Table 3**). We found that MOA and MOMENT showed the lowest memory usage among all the methods. The approximate MOA approach was the second fastest approach (only slightly slower than LFMM2-ridge) and the approximate MOMENT approach was slower than LFMM2-ridge, approximate MOA and ReFACTor but much faster than SVA, LFMM2-lasso and EWASher.

### An application of MOMENT to real data

We applied MOMENT and the other methods to four real quantitative traits in the LBC cohorts. These traits, including BMI, height, lung function (measured in the highest score of forced expiratory volume in one second), and walking speed (measured in the time taken to walk 6 meters), were standardised and corrected for age in each gender group within each sub-cohort (LBC1936 or LBC1921) (**Methods**). The standardised phenotypes were further processed by a rank-based inverse-normal transformation. The DNAm probes were adjusted for age, sex, and experimental batches. We did not adjust the probes for CTCs or smoking status for the purpose of testing methods (see below).

Consistent with the results from simulations, the test-statistics from MOA and MOMENT were not inflated whereas all the other methods showed modest inflation for all the traits (**Figure 3, Table 1**, and **Supplementary Figures 18-21**). Three associations were identified by multiple methods, including one for BMI (cg11202345, detected by all methods), in line with a previous study [49], and two for lung function (cg05575921 and cg05951221, detected by all methods except MOMENT) (**Supplementary Table 4, Supplementary Figures 18** and **20**). It should be noted that cg05575921 was reported to be associated with smoking in a previous study [50], indicating that the association between cg05575921 and lung function might be confounded by smoking status. Moreover, MOA, LFMM2-ridge, LFMM2-lasso, and ReFACTor consistently identified 12 additional probes associated with lung function but most of the probes have been linked to smoking in a previous study [51]. Almost all the associations were not significant when smoking status was fitted as a covariate in the models (6.5% of variance in lung function associated with smoking status; **Supplementary Table 5** and **Supplementary Figure 22**), suggesting that most (if not all) of the probe associations with lung function identified by MOA, LFMM2-ridge, LFMM2-lasso and ReFACTor were owing to the confounding of smoking. None of the smoking-associated probes were methylome-wide significant for lung function in the analysis using MOMENT (**Supplementary Figure 20**) and the result remained the same when smoking status was fitted as a covariate in MOMENT (**Supplementary Figure 22**), again demonstrating the capability of MOMENT in correcting for unobserved confounding factors. This is further supported by the finding from simulations that the effects of null probes estimated from MOMENT were much less correlated with the phenotype compared to those estimated from MOA (**Supplementary Figure 23**).

**Figure 3.**
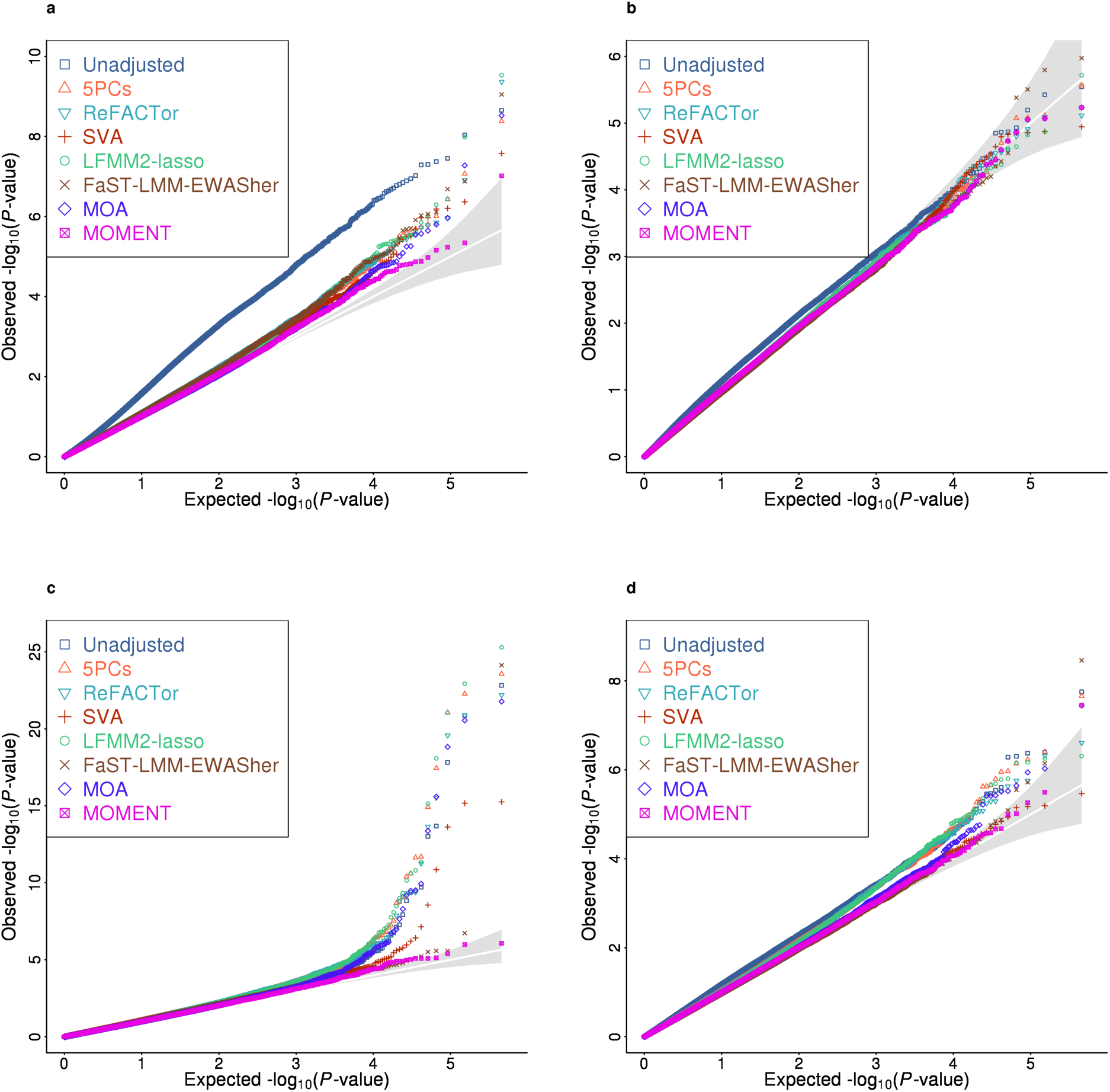
QQ plot of p-values from MWAS analysis for 4 quantitative traits in the LBC data. The DNAm measures were adjusted for age, sex, and batches. The phenotypes were stratified into groups by sex and cohort and were adjusted for age and standardised to z-scores by rank-based inverse normal transformation in each group. The phenotypes are (a) BMI, (b) height (c) lung function, and (d) walking speed.

**Table 1.**
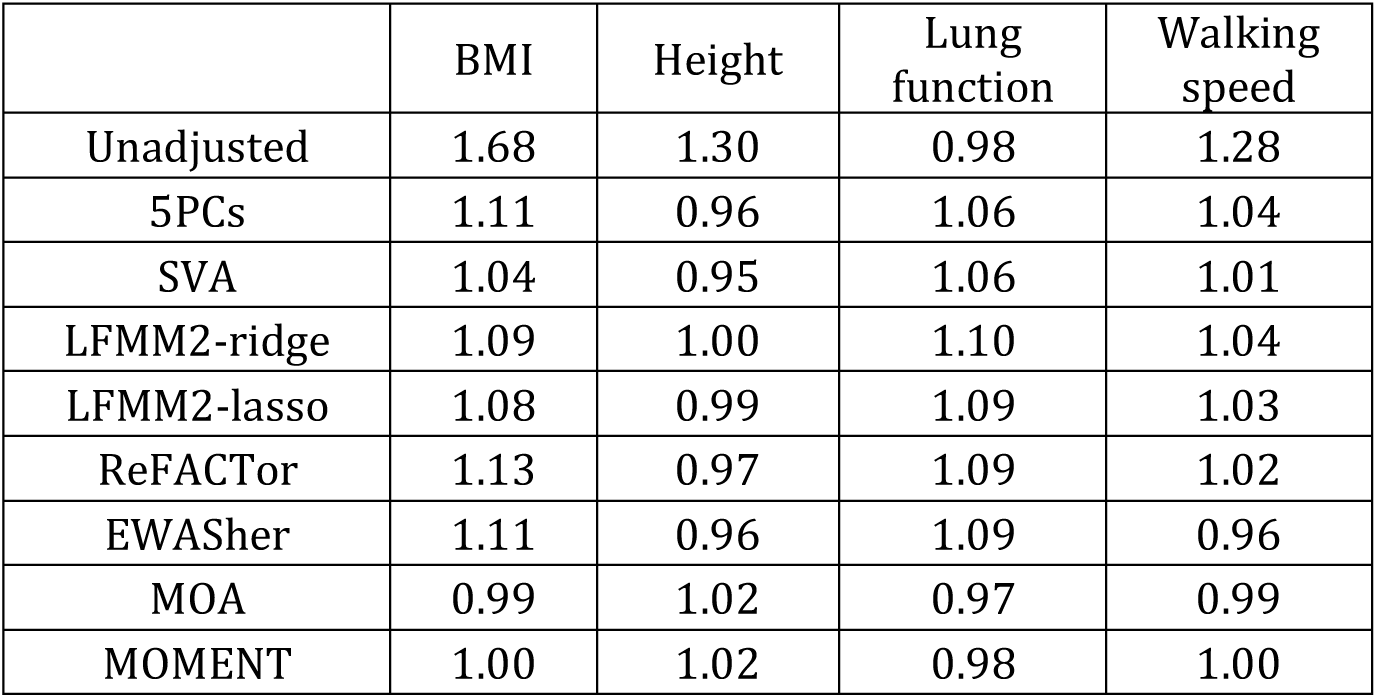
Genomic inflation factors reported by different MWAS methods for the 4 traits in the Lothian Birth Cohorts.

|  | BMI | Height | Lung<br>function | Walking<br>speed |
| --- | --- | --- | --- | --- |
| Unadjusted | 1.68 | 1.30 | 0.98 | 1.28 |
| 5PCs | 1.11 | 0.96 | 1.06 | 1.04 |
| SVA | 1.04 | 0.95 | 1.06 | 1.01 |
| LFMM2-ridge | 1.09 | 1.00 | 1.10 | 1.04 |
| LFMM2-lasso | 1.08 | 0.99 | 1.09 | 1.03 |
| ReFACTor | 1.13 | 0.97 | 1.09 | 1.02 |
| EWASher | 1.11 | 0.96 | 1.09 | 0.96 |
| MOA | 0.99 | 1.02 | 0.97 | 0.99 |
| MOMENT | 1.00 | 1.02 | 0.98 | 1.00 |

It has been shown in previous GWASs that MLM-based association analysis methods developed for quantitative traits are applicable to case-control data [37–39, 52]. We have shown by simulation that both MOMENT and MOA are applicable to case-control phenotypes regardless of whether cases are oversampled (**Supplementary Figures 15** and **16**). To demonstrate the applicability of the proposed methods to discrete phenotypes, we analysed smoking status (coded as 0, 1 or 2 for non-smoker, former smoker or current smoker) in the LBC by MOA and MOMENT in comparison with existing methods. All the methods detected a large number (at least 112) of probes at a methylome-wide significance level (*P* < 2.19e-7) except for MOMENT and EWASher which only identified 4 and 2 probes, respectively, at the methylome-wide significance level (**Supplementary Figure 24**). To validate the association signals other than those identified by MOMENT, we fitted the 4 MOMENT probes as fixed covariates in MOA. None of the additional associations remained methylome-wide significant conditioning on the 4 MOMENT probes (**Supplementary Figure 25**), suggesting that those additional associations detected by MOA (and other methods) were driven by their correlations with the 4 MOMENT signals. MOA failed in this scenario likely because the associations of the 4 MOMENT signals were too strong to be fitted in a single normal distribution with the other probes. This conclusion is further supported by the result that the accuracy of predicting/classifying smoking status in a cross-validation setting using a large number of probes detected by linear regression or MOA was even lower than that using a small number of probes detected by MOMENT (**Supplementary Table 6**). In addition, we recoded the smoking status data to a binary phenotype (0 for non-smoker and 1 for former or current smoker) and applied all the methods to the recoded binary phenotype; the conclusions were similar as above but it seemed that the analyses with the binary phenotype were less powerful than those with the categorical phenotype above (**Supplementary Figure 26**). All these results show the applicability of MOMENT to discrete traits and again demonstrate the robustness and reliability of MOMENT in controlling for false positive associations.

### Estimating variance in a phenotype captured by all probes by OREML

We have demonstrated the performance of the omic-data-based association analysis methods in OSCA by simulation and real data analysis. We then turned to evaluate the performance of OREML in estimating the proportion of variance in a complex trait captured by all probes (*ρ*^2^) by simulation in two scenarios (**Supplementary Note 1**). The results showed that under either scenario, OREML reported an unbiased estimate of *ρ*^2^ (**Supplementary Table 7**). Here the unbiasedness is defined as that the mean *ρ*^2^ estimate from 500 independent simulations is not significantly different from the *ρ*^2^ parameter used for simulation. There are two methods implemented in OSCA to compute the ORM (**Methods**). Our simulation results showed that the estimates of *ρ*^2^ based on the two methods were similar (**Supplementary Table 7**).

We also attempted to partition and estimate the proportions of phenotypic variation captured by all SNPs (i.e.,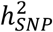) and all the DNAm probes respectively when fitted jointly in a model. We first investigated the correlation between genomic relationship matrix (GRM) and methylomic relationship matrix (MRM) in the LBC dataset. We found that the off-diagonal elements of the GRM were almost independent of those of the MRM (*r* = 0.0045; **Supplementary Figure 27**). From an OREML analysis that fits both the GRM and MRM, we estimated that all the DNAm probes captured 6.5% (SE = 0.038) of the variance for BMI but the estimate for height was nearly zero (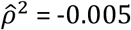 and SE = 0.0086) (**Supplementary Table 8**). These results are in line with the finding from a previous study that the accuracy of genetic risk prediction can be improved by incorporating DNAm data for BMI but not height [14].

## Discussion

In this study, we developed a versatile software tool—OSCA—to manage omic data generated from high-throughput experiments in large cohorts and to facilitate the analyses of complex traits using omic data (**Supplementary Note 4**). The primary applications of OSCA are to identify omic measures associated with a complex trait accounting for unobserved confounding factors (MOMENT) and to estimate the proportion of phenotypic variation captured by all measures of one or multiple omic profiles (OREML). A by-product of the OREML application is to estimate the joint effects of all measures of one or multiple omic profiles (i.e., OBLUP analysis) to predict the phenotype in a new sample. This has been shown to be a powerful and robust approach in age prediction using gene expression or DNAm data [53, 54]. We have also provided computationally efficient implementations in OSCA to manage large-scale omic data, and to perform omic-data-based quantitative trait locus (xQTL) analysis and meta-analysis of xQTL summary data. OSCA is an ongoing software development project so that any further methods or functions related to omic-data-based analysis can be included in the software package in the future.

We showed, by simulation, a surprisingly high error rate for all the existing MWAS/EWAS methods, mainly owing to the correlations between distal probes induced by CTCs (and/or other systematic confounders) in DNAm data (**Figure 1**). These correlations are widespread at a large number of probes across the methylome (as demonstrated by the proportion of null probes with *P*MWAS < 0.05 in simulation scenario 1; **Supplementary Figure 28**) and thus are not adequately accounted for by a fixed number of principal features computed from the data (e.g. 5PCs, ReFACTor, LFMM2, and SVA) nor a set of selected probes (e.g. FaST-LMM-EWASher). This conclusion is likely to be applicable to other types of omic data if the measures in distal genomic regions are correlated due to unmeasured confounding factors such as systematic experimental biases or unwanted biological variation, as suggested by our simulations with gene expression data (**Supplementary Figure 17**). This confounding effect can be corrected for by fitting the target probe as a fixed effect and all the other (distal) probes as random effects (i.e., the MOA or MOMENT method). In addition, we tested the robustness of MOMENT to the change of window size used to exclude probes in close physical proximity to the target probe in either direction. We varied the window size from 100Kbp to 250bp in the MOMENT analysis of data generated from simulation scenario 1 (**Supplementary Figures 29**). We found that the results remained almost unchanged when the window sizes decreased from 100Kbp to 25Kbp whereas there were a substantial number of probes showing deflated test-statistics when the window size decreased to 500bp or 250bp (**Supplementary Figures 29**). These results justify the use of 50Kbp as the default window size for MOMENT when applied to DNAm data. We also quantified the decay of correlation between a pair of gene expression probes as a function of their physical distance (**Supplementary Figure 30**), which suggests that 100Kbp is an appropriate MOMENT window size for gene expression data although the results remained almost unchanged when the window size was varied from 50Kbp to 1Mbp in the simulated data (**Supplementary Figure 31**).

Our simulation also showed that, if CTCs or batches explain a large proportion of variation in the phenotype, the FWERs of all the methods tended to be inflated (**Supplementary Figures 32** and **33**) despite that the genomic inflation factor is close to unity for most methods (**Figure 2**). We re-ran the simulation under a more extreme setting with 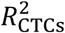 varying from 10% to 70%. In this case, the genomic inflation factors of the fixed-effect models (i.e., SVA, ReFACTor, LFMM2, and 5PCs) and the FWERs of all the methods increased as 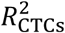 increased (to a lesser extent for FaST-LMM-EWASher), suggesting that there were a set of probes strongly associated with CTCs (**Supplementary Figure 34**). Note that even in this extreme case, MOMENT showed the lowest FWERs on average amongst all the methods. It is also of note that the FWERs of FaST-LMM-EWASher were relatively low in this scenario (**Supplementary Figure 32**), opposite to its performance when 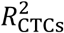 was low (**Figure 2**), possibly due to its variable selection strategy (**Supplementary Note 2**). The inflation in FWER was only slightly alleviated by fitting the predicted CTCs as covariates (**Supplementary Figures 35** and **36**). The results also suggest that it may be worth fitting measured CTCs as fixed-effect covariates in MLM-based association analyses such as MOA and MOMENT in practice although this approach is likely to be conservative as indicated by the deflated *λ* and FWER (**Supplementary Figure 37**). These conclusions are likely to be applicable to other confounding factors such as smoking status, as demonstrated in the analysis of lung function data in the LBC (**Supplementary Figure 22**). Our results also caution the interpretation of associations identified from MWAS for traits that are highly correlated with CTCs and/or other biological confounders. In addition, although our simulation shows that both MOMENT and MOA are applicable to case-control phenotypes (**Supplementary Figures 15** and **16**), direct application of linear model approaches to 0/1 traits is not ideal. If the underlying model is causal (i.e., omic measures have causal effects on the trait), a more appropriate analysis is to use a link function (e.g., a probit or logit model) that connects the 0/1 phenotype to a latent continuous trait, as in the methods recently developed for the analysis of case-control data in GWAS [55–58]. Since OSCA is an ongoing software development project, the non-linear link functions can be incorporated in the MOMENT/MOA framework in the future.

In conclusion, we showed by simulation the inflation in test-statistics of the existing MWAS methods because of the ubiquitous correlations between distal probes caused by confounding factors, and developed two new MWAS methods (MOA and MOMENT) to correct for the inflation. We demonstrated the reliability and robustness of MOMENT by simulations in a number of scenarios and real data analyses. We recommend the use of MOMENT in practice because of its robustness in the presence of unobserved confounders despite that it is slightly less powerful than MOA. We implemented both MOA and MOMENT in a computationally efficient and easy-to-use software tool OSCA together with many other functions for omic-data-based analyses (**Supplementary Figure 38**).

## Methods

### Omic-data-based relationship matrix (ORM)

We have described in equations [1] and [2] the OREML model to estimate the proportion of variance in a phenotype captured by the DNAm probes all together. In equation [1], i.e., **y** = **Cβ** + **Wu** + **e**, we define **W** as a matrix of standardised DNAm measures of all probes, and in equation [2] we define the ORM as **A** = **WW***’*/*m*. Therefore, the omic relationship between individual *j* and *k* (the *jk*-th element of **A**) can be computed as 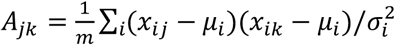, where *x*_*Ij*_ is the unstandardised DNAm level of probe *i* in individual *j, µ*_*i*_ and 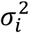 are the mean and variance of the *i*-th probe over all the individuals respectively, and *m* is the number of probes. This model implicitly assumes that the probes of smaller variance in DNAm level (unstandardised) tend to have larger effects on the phenotype (strictly speaking, stronger associations with the phenotype), and that there is no relationship between the proportion of trait variance captured by a probe and the variance of the probe. We also provide in OSCA an alternative method to compute the ORM, i.e., 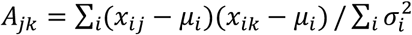. If we use this definition of ORM in the OREML analysis, we implicitly assume that there is no relationship between the probe effect on the trait and the variance of the probe but the proportion of trait variance associated with a probe increases as the variance of the probe increases. We showed by simulation and real data analysis that the difference between OREML results using the two methods was very small (**Supplementary Tables 7** and **8**).

### OREML: estimating the proportion of trait variance captured by all DNAm probes

We have shown in equations [1] and [2] an OREML model with one random-effect component to estimate the proportion of trait variance captured by all DNAm probes. The model is flexible, which can be extended to partition the trait variance into components associated with different sets of probes (e.g. a model with 22 components with all the probes on each chromosome as a component). A flexible OREML model can be written as

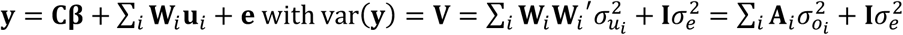

where the definitions of all the parameters and variables are similar to those in equations [1] and [2]. The variance components can be estimated by REML [33], and the proportion of the trait variance captured by the *i*-th component can be computed as 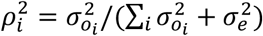

The multi-component OREML model can be applied to partition the trait variance into components associated with multiple omic profiles. For example, if SNP genotype, DNAm, and gene expression data are available for all the individuals in a cohort, a multi-component OREML model can be used to estimate the proportion of trait variance captured by all SNPs (i.e., the SNP-based heritability), the expression levels of all genes, and the DNAm levels at all the CpG sites. The model can be written as

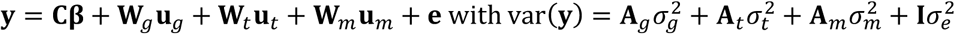

where **W**_*g*_, **W**_*t*_, and **W**_*m*_ are the matrices of standardised SNP genotypes, gene expression measures, and DNAm levels, respectively, with the corresponding effects **u**_*g*_, **u**_*t*_ and 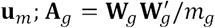 is the genomic relationship matrix (GRM) with *m*_*g*_ being the number of SNPs 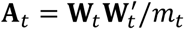 is the transcriptomic relationship matrix (TRM) with *m*_*t*_ being the number of transcripts, and 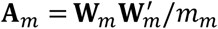 is the methylomic relationship matrix (MRM) with *m*_*m*_ being the number of DNAm probes. Note that the model can be reduced by dropping any of the variance components or expanded by including other types of omic profiles (e.g., protein abundance).

### Dataset

The LBC cohorts [59, 60] consisted of individuals born in 1921 (LBC1921) and 1936 (LBC1936), mostly living in Edinburgh city and the surrounding Lothian region of Scotland. Blood samples were collected with informed consent. The use of human participants in this study was approved by The University of Queensland Human Research Ethics Committee B (approval number: 2011001173). The LBC individuals underwent several waves of SNP genotyping and DNAm measures. DNAm levels at 485,512 CpG sites across the genome were measured on 3,191 whole blood samples from 3 waves using the Illumina HumanMethylation450 BeadChip. Duplicates or samples with an excessive proportion of low confidence calls across all probes (>5%) were removed. Probes with an excessive proportion of low confidence calls across all individuals (>5%) or probes located in sex chromosomes were excluded. In addition, probes encompassing SNPs annotated in dbSNP131 using hg19 coordinates or identified as potentially cross-hybridized methylation probes by a previous study [61] were also excluded. After these QC steps, 3,018 samples and 307,360 probes remained (**Supplementary Note 3**). We included in the analysis only the first wave (wave1) of the LBC data consisting of 436 individuals from LBC1921 (average age of 79 years) and 906 individuals from LBC1936 (average age of 70 years) (**Supplementary Table 9**). We removed probes with almost invariable beta values across individuals (standard deviation < 0.02) and retained 1,342 individuals and 228,694 probes for analysis.

There were a number of covariates available in the LBC data including age, sex, batches of the experiment (i.e. plate and position of the sample on a chip), and CTCs. The blood cell counts for different cell types, including basophils, eosinophils, monocytes, lymphocytes, and neutrophils, were quantified using an LH50 Beckman Coulter instrument on the same day of blood collection. In addition to the covariates, there are a number of traits measured on the LBC individuals including height (measured without shoes), body mass index (BMI), lung function (measured in the highest score of forced expiratory volume in one second), and walking speed (measured in the time taken to walk 6 metres) and smoking status (never smoked, ex-smoker or current smoker) [62, 63]. The numbers of missing measurements are noted in **Supplementary Table 10**. For each trait, we adjusted the phenotype for age in each gender group of each cohort (LBC1921 or LBC1936) and standardised the residuals by rank-based inverse normal transformation, which removed the age effect and potential difference in mean and variance between two gender groups or cohorts.

The LBC wave1 individuals were also genotyped by Illumina 610-Quadv1 BeadChip. The QC process of the SNP genotype data has been detailed elsewhere [14]. After excluding SNPs from sex chromosomes and SNPs with low allelic frequency (MAF < 0.01), we retained 523,062 genotyped SNPs for analysis.

We also used a set of gene expression data available at EMBL-EBI (URLs) from the San Antonio Family Heart Study (SAFHS). Sample recruitment, data generation and quality controls of the SAFHS data have been detailed elsewhere [46–48]. We used the processed and standardized gene expression data of 19,648 autosomal probes on 1,240 non-diseased Mexican American participants. Age, sex and smoking status were available in the data. We removed 21 samples with unknown smoking status and retained 1,219 individuals for analysis.

### URLs

OSCA, http://cnsgenomics.com/software/osca

ReFACTor, https://www.cs.tau.ac.il/~heran/cozygene/software/refactor.html

EWASher, https://www.microsoft.com/en-us/research/project/fast-lmm-software-papers/

SVA, https://bioconductor.org/packages/release/bioc/html/sva.html

LFMM2, https://bcm-uga.github.io/lfmm/

The LBC data: https://www.ebi.ac.uk/ega/studies/EGAS00001000910

The SADHS data: https://www.ebi.ac.uk/arrayexpress/experiments/E-TABM-305/

## Acknowledgements

This research was supported by the Australian Research Council (FT180100186), the Australian National Health and Medical Research Council (grants 1107258, 1113400, 1083656, 1078037, and 1078901) and the Sylvia & Charles Viertel Charitable Foundation. The Lothian Birth Cohorts (LBC) are supported by Age UK (Disconnected Mind programme). Methylation typing was supported by Centre for Cognitive Ageing and Cognitive Epidemiology (Pilot Fund award), Age UK, The Wellcome Trust Institutional Strategic Support Fund, The University of Edinburgh, and The University of Queensland. The LBC resource is prepared in the Centre for Cognitive Ageing and Cognitive Epidemiology, which is supported by the Medical Research Council and Biotechnology and Biological Sciences Research Council (MR/K026992/1), and which supports I.J.D..

## Authors’ contributions

JY conceived the study. JY and FZ designed the experiment. FZ developed the software tool and performed all the analyses under the guidance and/or assistance from JY, ZZ, WC, QZ, MFN and TQ. JY, AFM, PMV and NRW contributed funding and resources. IJD., PMV. and NRW contributed the DNA methylation data. FZ, JY and WC wrote the manuscript with the participation of all authors.

