## Supplementary for "OSCA: a tool for omic-data-based complex trait analysis"

Zhang *et al*.

**Supplementary Note 1: Simulating phenotypes from DNAm data**

**Simulation scenario 1: simulating a quantitative phenotype with effects from a set of causal probes (on the odd chromosomes)**

We used the DNAm data from the LBC cohort after quality control (**Methods**). In the simulation, the DNAm probes were adjusted for age, sex, experimental batches, and smoking status but not CTCs. The phenotype was simulated based on the following model

$$\boldsymbol{y} = \mathbf{X}\boldsymbol{b} + \boldsymbol{e} \quad (1)$$

where $\boldsymbol{y}$ is a vector of phenotypes, $\mathbf{X}$ is a matrix of standardised DNAm measures for 1,319 individuals and a set of causal probes (randomly sampled from all probes on the odd chromosomes), $\boldsymbol{\beta}$ is a vector of the effects of the causal probes, and $\boldsymbol{e}$ is a vector of residuals. The probe effects were generated from a standard normal distribution, and the elements in $\boldsymbol{e}$ were generated from $N(0, \sigma^2_{\mathbf{X}\boldsymbol{b}}(\frac{1}{\rho^2} - 1))$ with $\sigma^2_{\mathbf{X}\boldsymbol{b}}$ being the empirical variance of the elements in vector $\mathbf{X}\boldsymbol{b}$ and $\rho^2$ being the proportion of phenotypic variance explained by the causal probes. We simulated the phenotype with 100 causal probes with $\rho^2 = 0.5$ (per-probe $\rho^2 = 0.005$), and repeated the simulation 100 times with the causal probes resampled in each simulation replicate. We also tested the robustness of the models under two additional settings, i.e., 10 causal probes with $\rho^2 = 0.2$ (per-probe $\rho^2 = 0.02$) and 1000 causal probes with $\rho^2 = 0.4$ (per-probe $\rho^2 = 0.004$).

In each simulation replicate, we tested the associations between the simulated phenotype and all the probes using different methods. The genome-wide significance level was set to 2.19e-07 based on a Bonferroni correction for 228,694 tests.

**Simulation scenario 2: simulating a quantitative phenotype with effects from cell types but no direct effects from the probes**

We used the DNAm data and cell counts measured for five primary blood cell types in the LBC cohort to simulate a phenotype based on the following model

$$\boldsymbol{y} = \mathbf{C}\boldsymbol{\beta} + \boldsymbol{e} \quad (2)$$

where $\boldsymbol{y}$ is a vector of phenotypes, $\mathbf{C}$ is a matrix of standardised cell counts for 1,319 individuals and 5 cell types, $\boldsymbol{\beta}$ is a vector of the effects of the cell types, and $\boldsymbol{e}$ is a vector of residuals. The cell type effects were generated from a standard normal distribution, and the elements of $\boldsymbol{e}$ were generated from $N(0, \sigma^2_{\mathbf{C}\boldsymbol{\beta}}(\frac{1}{R^2_{\mathrm{CTCs}}} - 1))$ with $\sigma^2_{\mathbf{C}\boldsymbol{\beta}}$ being the empirical variance of all the elements in vector $\mathbf{C}\boldsymbol{\beta}$ and $R^2_{\mathrm{CTCs}}$being the proportion of phenotypic variance explained by CTCs. Given the observed variance explained by 5 cell types (i.e., neutrophils, lymphocytes, monocytes, eosinophils and basophils) that varied from 0 to ~0.05 for 4 real traits (i.e., BMI, height, lung function and walking speed) in LBC (**Supplementary Table 11**), we used a range of parameter values for $R^2_{\text{CTCs}}$ in the simulation, i.e., 0.005, 0.01, 0.02, 0.03, 0.04 and 0.05. We repeated each simulation scenario 1000 times (we ran more simulation replicates in this case to get a more precise estimate of power). In addition, we also simulated extreme cases with $R^2_{\text{CTCs}}$ = 0.1, 0.3, 0.5 and 0.7. In each simulation replicate, we performed the association tests using different methods and used a genome-wide significance level of 2.19e-07 based on a Bonferroni correction for 228,694 tests.

We also applied the method to simulate a phenotype with 100 replicates based on 22 experimental batches (i.e., 22 plates) in lieu of the 5 cell types with $R^2_{\text{batch}}$ ranging from 0.005 to 0.7. Note that in this case the DNAm probes were adjusted for age, sex, sample position on the chip, CTCs and smoking status.

**Simulation scenario 3: simulating a quantitative phenotype with effects from both causal probes and cell types**

In this scenario, we used the LBC data to simulate a phenotype based on the model below

$$\boldsymbol{y} = \mathbf{C}\boldsymbol{\beta} + \mathbf{X}\boldsymbol{b} + \boldsymbol{e} \qquad (3)$$

where all the parameters and variables have been defined above. The five cell types explained 5% of variance of the simulated phenotype and the 100 causal probes (randomly sampled from all probes on the odd chromosomes) explained 50% of the variance. The simulation was repeated 100 times with causal probes resampled in each simulation replicate.

We also applied the method to simulate a phenotype with 100 replicates based on 100 causal probes (explaining 50% of the variance) and 22 experimental batches (explaining 5% of the variance). In this case, the DNAm probes were adjusted for age, sex, sample position on the chip, CTCs and smoking status.

**Simulations to evaluate OSCA-OREML**

We simulated a phenotype based on model (1), i.e. $\boldsymbol{y} = \mathbf{X}\boldsymbol{b} + \boldsymbol{e}$. In this simulation, we did not limit the causal probes on the odd chromosomes but allowed them to be sampled at random across the whole genome. We simulated the phenotype in 2 scenarios, 1) the null model (i.e., $\mathbf{Xb}$ = $\mathbf{0}$), 2) the alternative model ($m$ causal probes ramdomly sampled from all the probes with $m$ = 1, 50 or 500 and the correponding $\rho^2$ = 0.1, 0.2 or 0.5) using the method described above. Each DNA methyaltion probe in LBC was adjusted for age, sex, experimental batches, smoking status and CTCs. We repeated each simulation 500 times to obtain the mean and standard deviation of the OREML estimate over repeated sampling.

**Supplementary Note 2: A brief summary of the feature selection strategy used in FaST-LMM-EWASher**

FaST-LMM-EWASher is an MLM-based EWAS method which tests the association of a probe with a trait conditioning on the random effects of a set of selected probes. The main analysis pipeline implemented in FaST-LMM-EWASher can be summarised as following:

1. Read the DNAm and phenotype data.
2. Remove probes with small and large mean beta values.
3. Compute the principal components for the individuals using the DNAm data.
4. Run MLM-based association analyses iteratively with increasing number of PCs (from 0 to 10) fitted as covariates. In each iteration, a feature selection process is conducted to select probes to be fitted as the random effects in the MLM (not including the probes in <50Kb distance from the probe in question). The iteration proceeds until the genome inflation factor is smaller than 1 or the number of PCs is 10.

The algorithm to select the random-effect probes can be described as

1) Split the sample into 10 sub-samples of the same size
2) For each sub-sample
    a. Use the target sub-sample as the test set and a pool of all the other sub-samples as the training set.
    b. Run linear regression analysis in the training set.
    c. Define a vector with fixed numbers, i.e., $\boldsymbol{v}$ = [1, 101, 201, 301, 401, 501, 601, 701, 801, 901, 2000, 3000, 4000, 5000, 6000, 7000, 8000, 9000]
    d. For each number in the vector $\boldsymbol{v}$ above, take the top $v_i$ probes ranked by the linear regression p-values in the training set to predict the phenotype in the test set.
    e. Calcualte SSE and the log likelihood from regressing the phenotype against the predictor in the test set.
3) Calculate the mean values of SSE and log likelihood across the 10 sub-samples for $v_i$.
4) Select a number in vector $\boldsymbol{v}$ (denoted by $v_{max}$) that shows the largest mean log likelihood or the smallest mean SSE.
5) Select the top $v_{max}$ probes from linear regression analysis in the whole sample.

**Supplementary Note 3: Quality control of the DNA methylation data in LBC**

A total of 3,191 whole blood samples from 3 waves were assayed with the Illumina HumanMethylation450 Bead Chips at 485,512 CpG sites across the genome. We removed 146 duplicated samples or samples for which >5% of the probes with intensity detection (against negative probes on the array) p-value >0.01. We also removed probes with intensity detection p-value >0.01 in >5% of the samples and probes on the sex chromosomes. We further excluded probes encompassing SNPs annotated in dbSNP and probes identified as potentially cross-hybridizing. A total of 3,018 samples and 307,360 probes were retained for analysis (**Supplementary Table 12**).

**Supplementary Note 4: Key features of OSCA**

We have mentioned in the main text the two primary applications of OSCA (OSCA-MOMENT and OSCA-OREML). Here, we listed a number of additional key features of OSCA (**Supplementary Figure 39**).

- Genetic association analysis based on linear regression for molecular traits, also known as omic-data-based quantitative trait locus (xQTL) analysis. The xQTL analysis is computationally intensive because of the large number of traits (e.g., tens of thousands of traits for gene expression and hundreds of thousands of traits for DNA methylation). To speed up the analysis, OSCA parallelises the analysis to array jobs and uses multi-threading to utilise multiple CPUs in one job. For analyses with a number of covariates, OSCA has an option to accelerate the computation by pre-adjusting the phenotype and omic measures by the covariates. The xQTL output (i.e., the summary statistics) are stored in the SMR BESD format [1], greatly alleviating the storage burden and also facilitating the follow-up analyses using SMR.
- Omic-data-based phenotype prediction. We have implemented in OSCA the omic-data-based best linear unbiased predictor (OBLUP) [2] to compute the joint effects of all measures of an omic profile, which can subsequently be used to predict the phenotypes of individuals in an independent sample. Note that the OBLUP analysis needs to be performed in conjunction with OREML. We will implement Lasso, Elastic net and multi-component OBLUP approaches in OSCA for phenotype prediction in the near future.
- Simulating phenotype based on omic data (see the **Supplementary Note 1** for the simulation method).
- Meta-analysis of xQTL summary statistics from multiple cohorts with or without sample overlap. OSCA estimates sample overlap between pairwise cohorts using summary statistics of the non-significant SNPs and performs a meta-analysis account for sampling covariance among the estimates from multiple cohorts by a generalised least squares approach [3]. If all the cohorts are independent, the method is equivalent to the conventional inverse-variance-weighted meta-analysis approach [4].
- Data storage. For the efficiency of data management, we store the omic data in binary format (BOD file format) in OSCA, which is more efficient to read and write compared to text format, and save the xQTL output in SMR BESD format as mentioned above. We also provide functions that support the conversion between a BOD file and a plain text file.

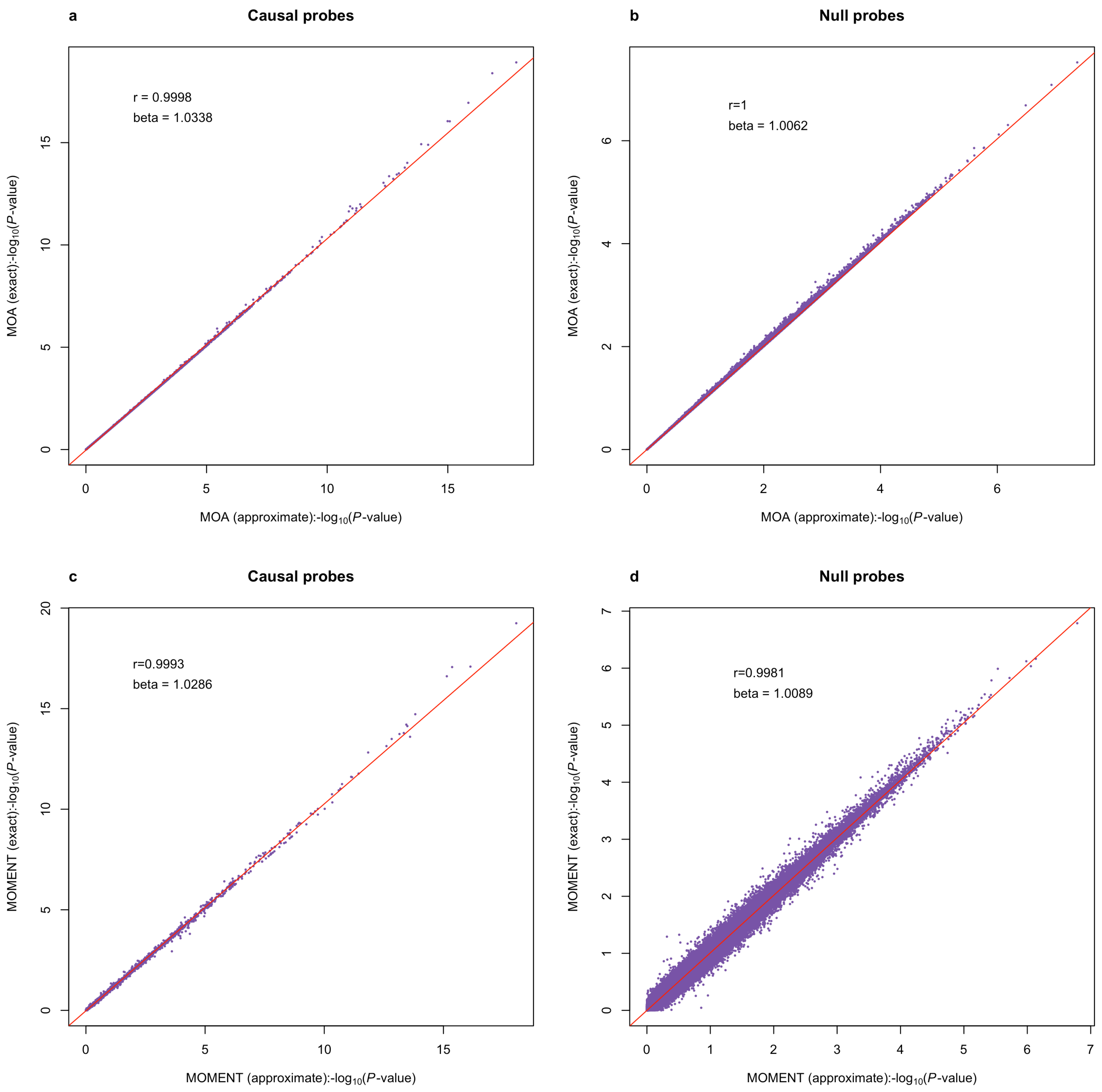


**Supplementary Figure 1** Comparison between the approximate and exact MOA/MOMENT approaches in simulation scenario 1. Shown are the pooled results from 30 simulation replicates. (a) approximate vs. exact MOA for the causal probes; (b) approximate vs. exact MOA for the null probes; (c) approximate vs. exact MOMENT for the causal probes; (b) approximate vs. exact MOMENT for the null probes; r = Pearson correlation; beta = regression coefficient.

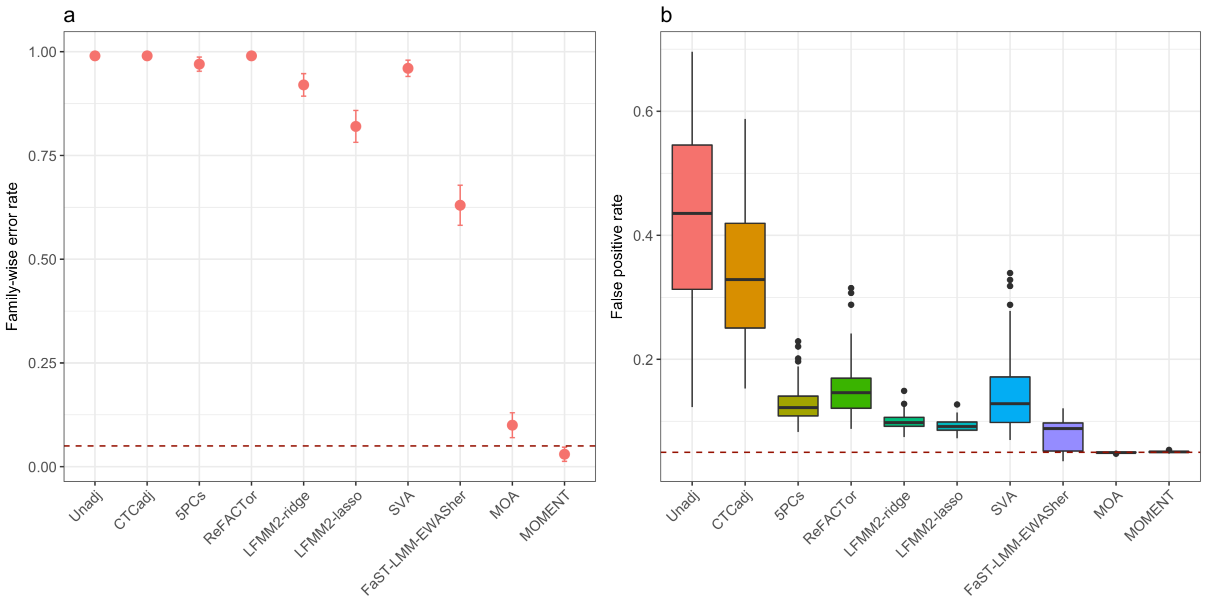


**Supplementary Figure 2** FWER and FPR for the MWAS methods in simulation scenario 1. The phenotypes were simulated based on the effects from 100 causal probes but no direct effects from the CTCs. (a) Each dot represents the FWER of a method (with an error bar representing +/- the SE). The dashed line at 0.05 is the expected FWER under the null. (b) False positive rate of a method from 100 simulation replicates, calculated as the proportion of null probes with p-values < 0.05 in each simulation replicate.

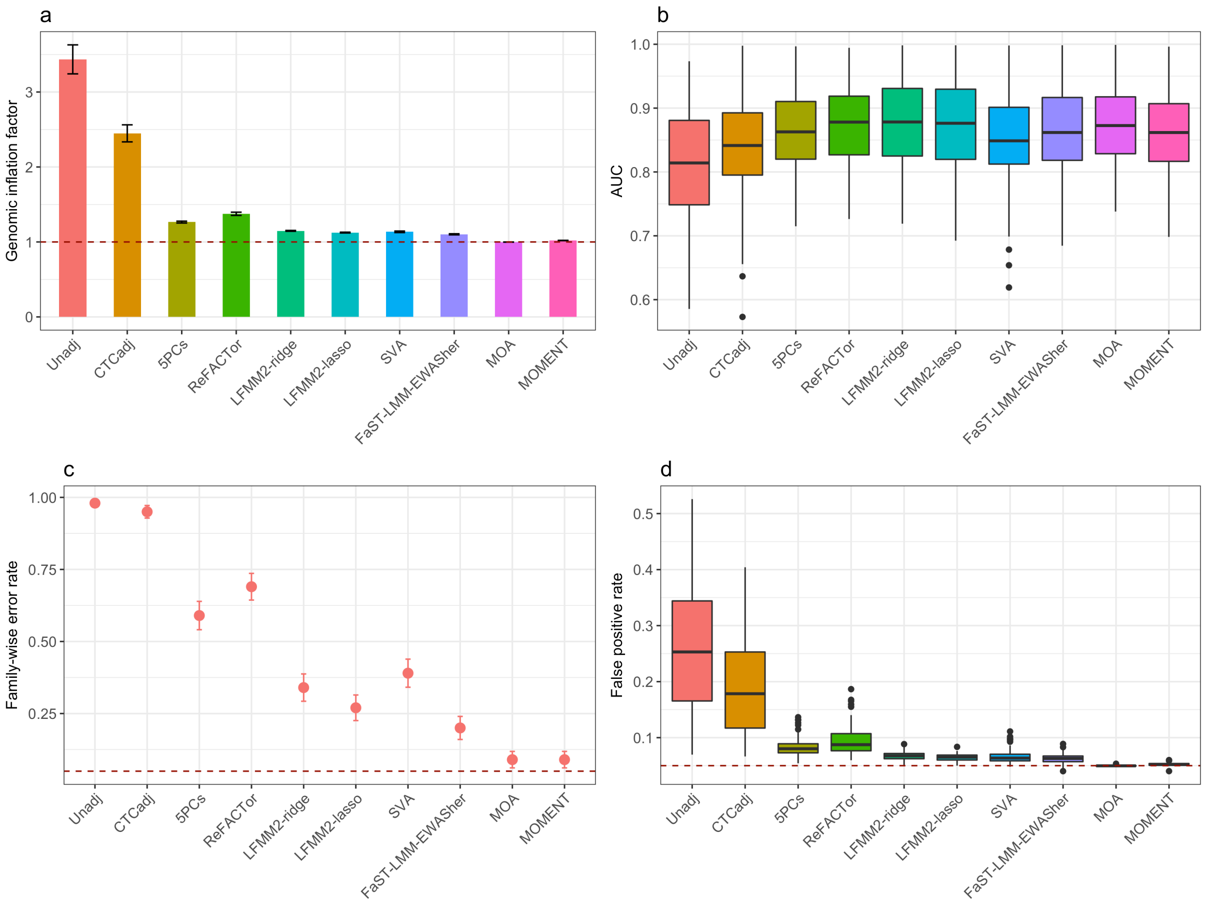


**Supplementary Figure 3** Power and false positive rate for the MWAS methods in simulation scenario 1 (phenotypes simulated based on effects from 10 causal probes). (a) Each column represents the mean of genomic inflation factor from a method across 100 simulation replicates with an error bar representing +/- SE of the mean. (b) Each column is a boxplot of the AUCs for a method from 100 simulation replicates. (c) Each dot represents the FWER of a method (with an error bar representing +/- the SE), where FWER is defined as the proportion of simulation replicates with at least one null probe at MWAS p-value < 0.05 / $m$ with $m$ being the number of probes. (d) Each column is a boxplot of FPRs of a method from 100 simulation replicates, where FPR is defined as the proportion of null probes with p-values < 0.05 in each simulation replicate.

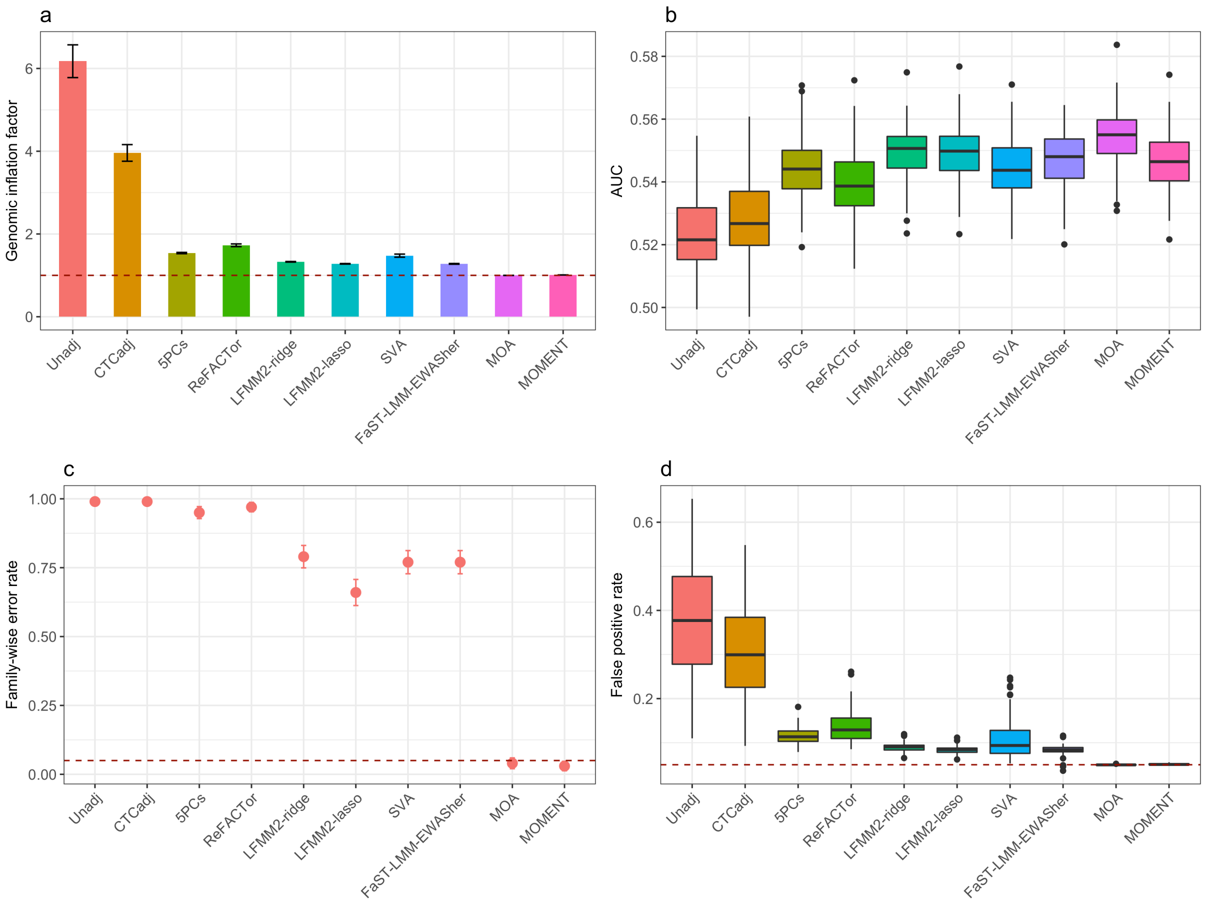


**Supplementary Figure 4** Power and false positive rate for the MWAS methods in simulation scenario 1 (phenotypes simulated based on effects from 1000 causal probes). See **Supplementary Figure 3** for details of the labels.

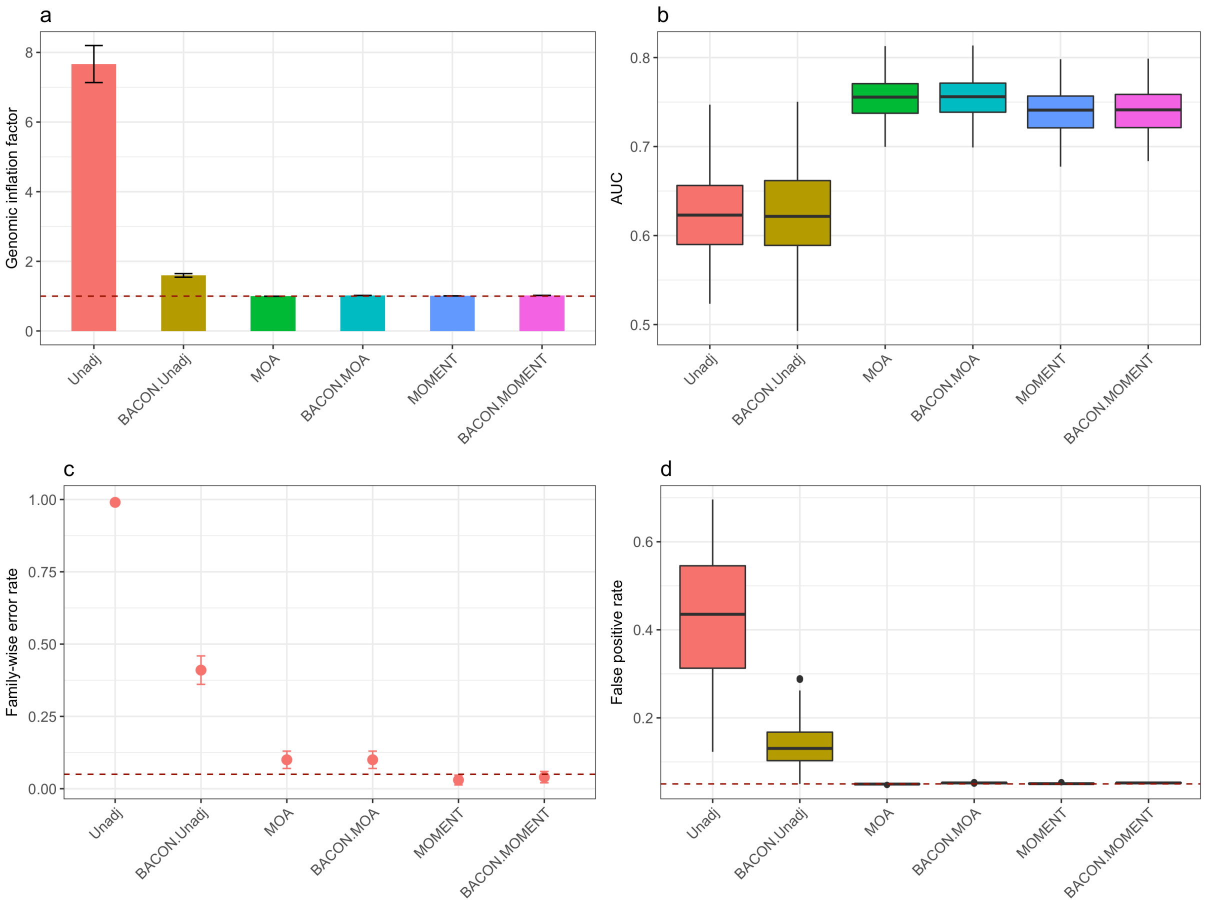


**Supplementary Figure 5** Genome inflation factor and false positive rate for Unadj, MOA and MOMENT before and after the BACON adjustment in simulation scenario 1 (phenotypes simulated based on effects from 100 causal probes). See **Supplementary Figure 3** for details of the labels.

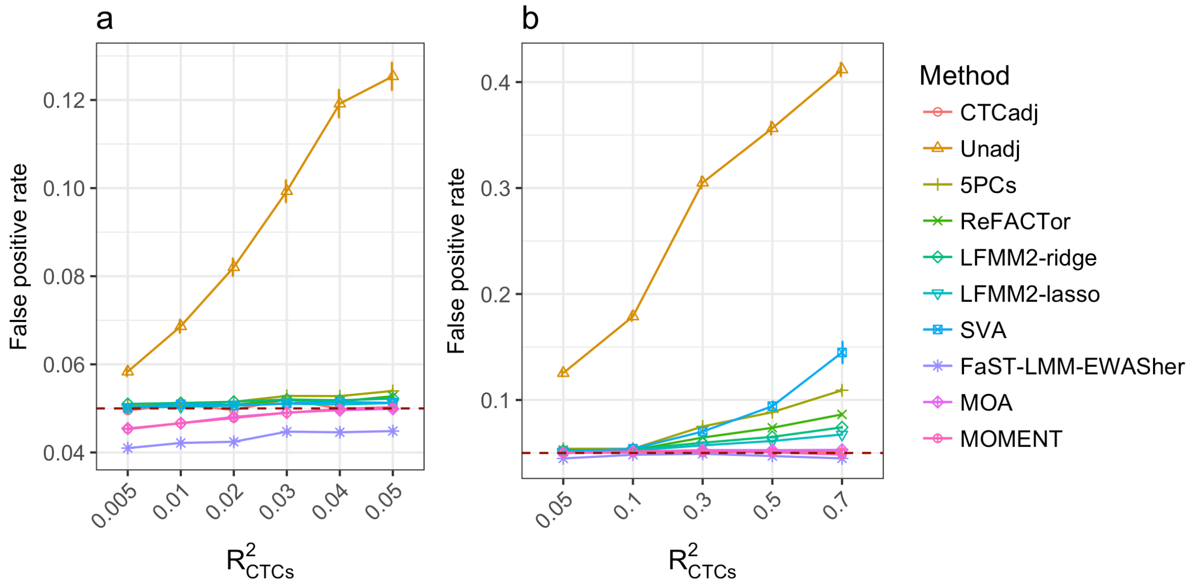


**Supplementary Figure 6** False positive rates of the MWAS methods in simulation scenario 2 (effects from CTCs but no causal probes). Shown on the horizontal axis are the $R^2_{\mathrm{CTCs}}$ values used to simulate the phenotype. Each dot represents the mean FPR (the proportion of null probes with p-values < 0.05 in each simulation replicate) from 1000 simulation replicates.

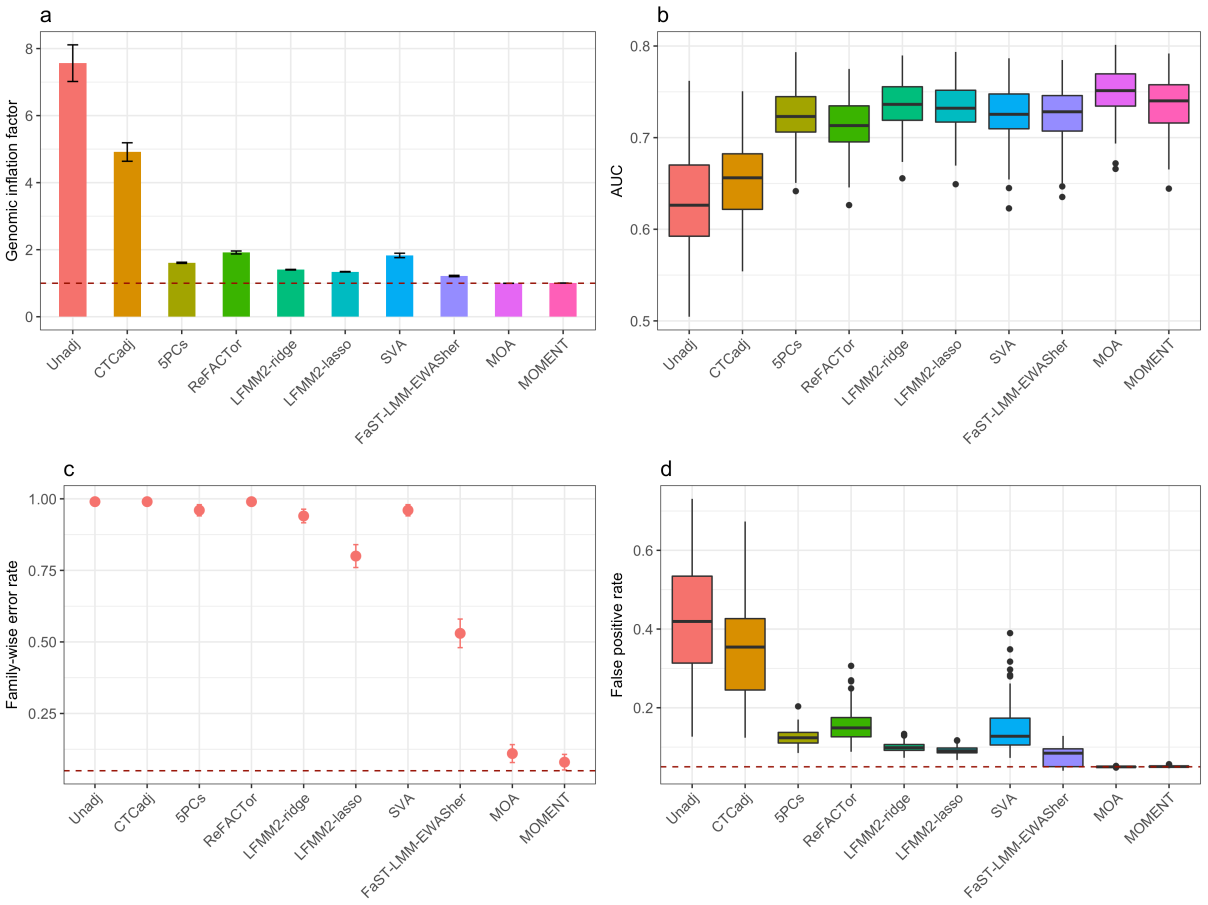


**Supplementary Figure 7** Power and false positive rate for the MWAS methods in simulation scenario 3 (phenotypes simulated based on effects from 100 causal probes and 5 CTCs). See **Supplementary Figure 3** for details of the labels.

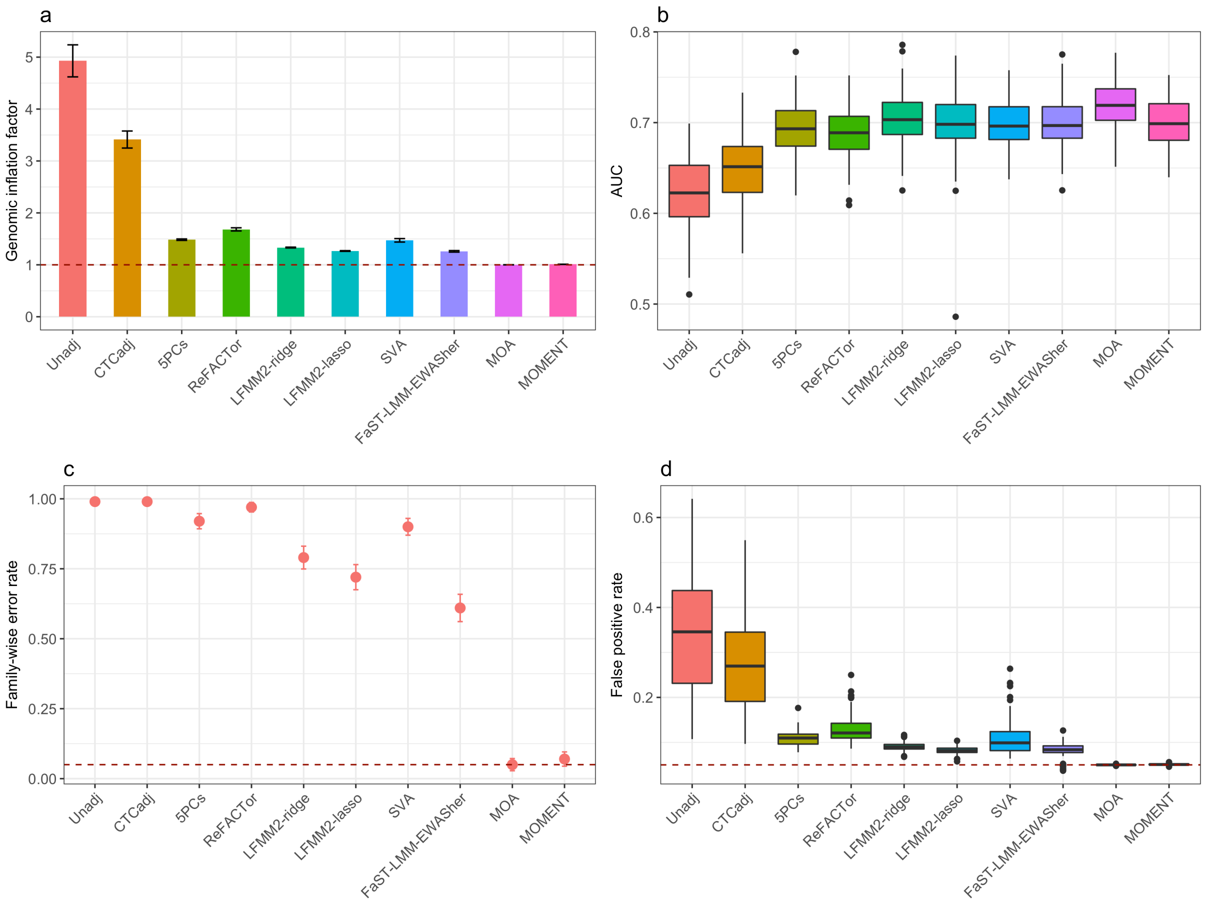


**Supplementary Figure 8** Power and false positive rate for the MWAS methods in simulation scenario 3 (phenotypes simulated based on effects from 100 causal probes and 5 CTCs) with LBC1936. See **Supplementary Figure 3** for details of the labels.

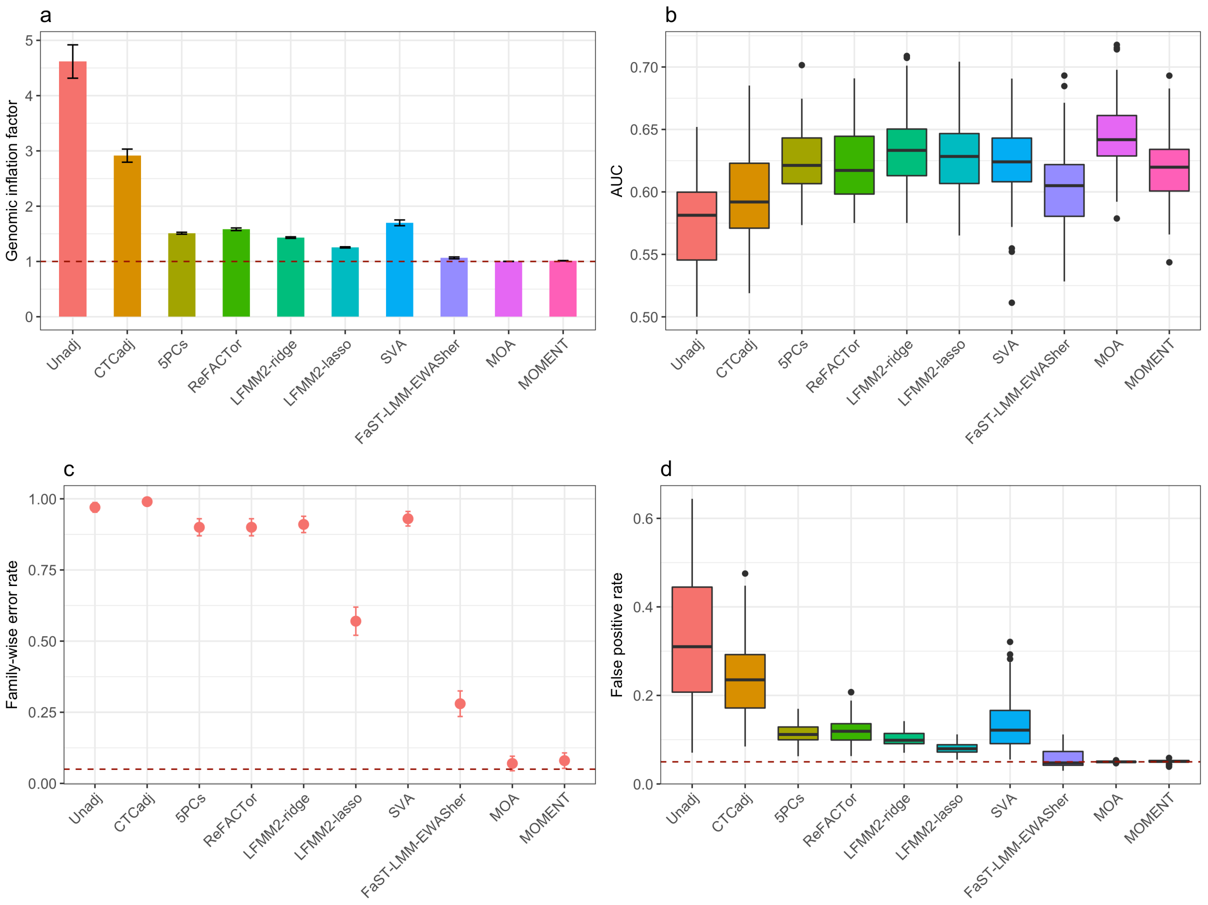


**Supplementary Figure 9** Power and false positive rate for the MWAS methods in simulation scenario 3 (phenotypes simulated based on effects from 100 causal probes and 5 CTCs) with LBC1921. See **Supplementary Figure 3** for details of the labels.

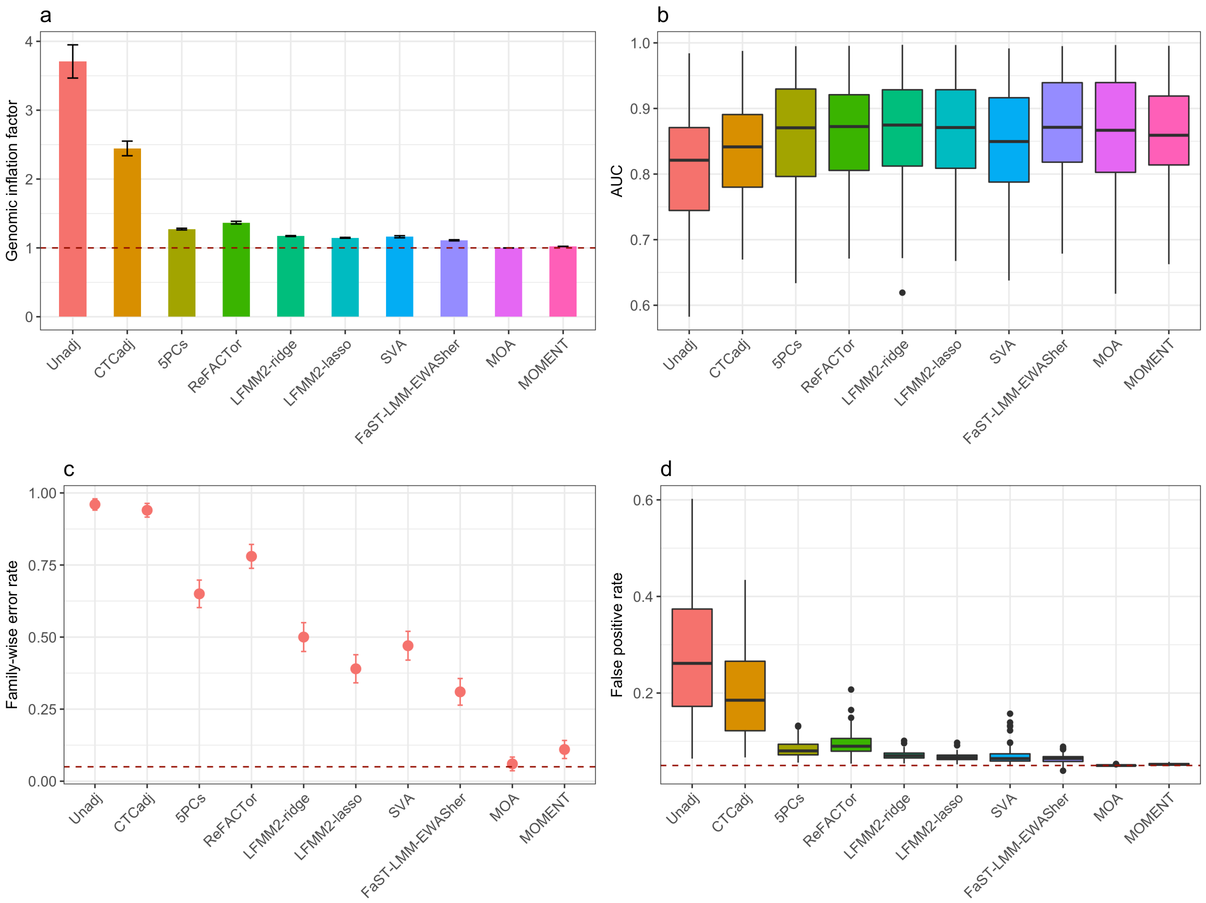


**Supplementary Figure 10** Power and false positive rate for the MWAS methods in simulation scenario 3 (phenotypes simulated based on effects from 10 causal probes and 5 CTCs). See **Supplementary Figure 3** for details of the labels.

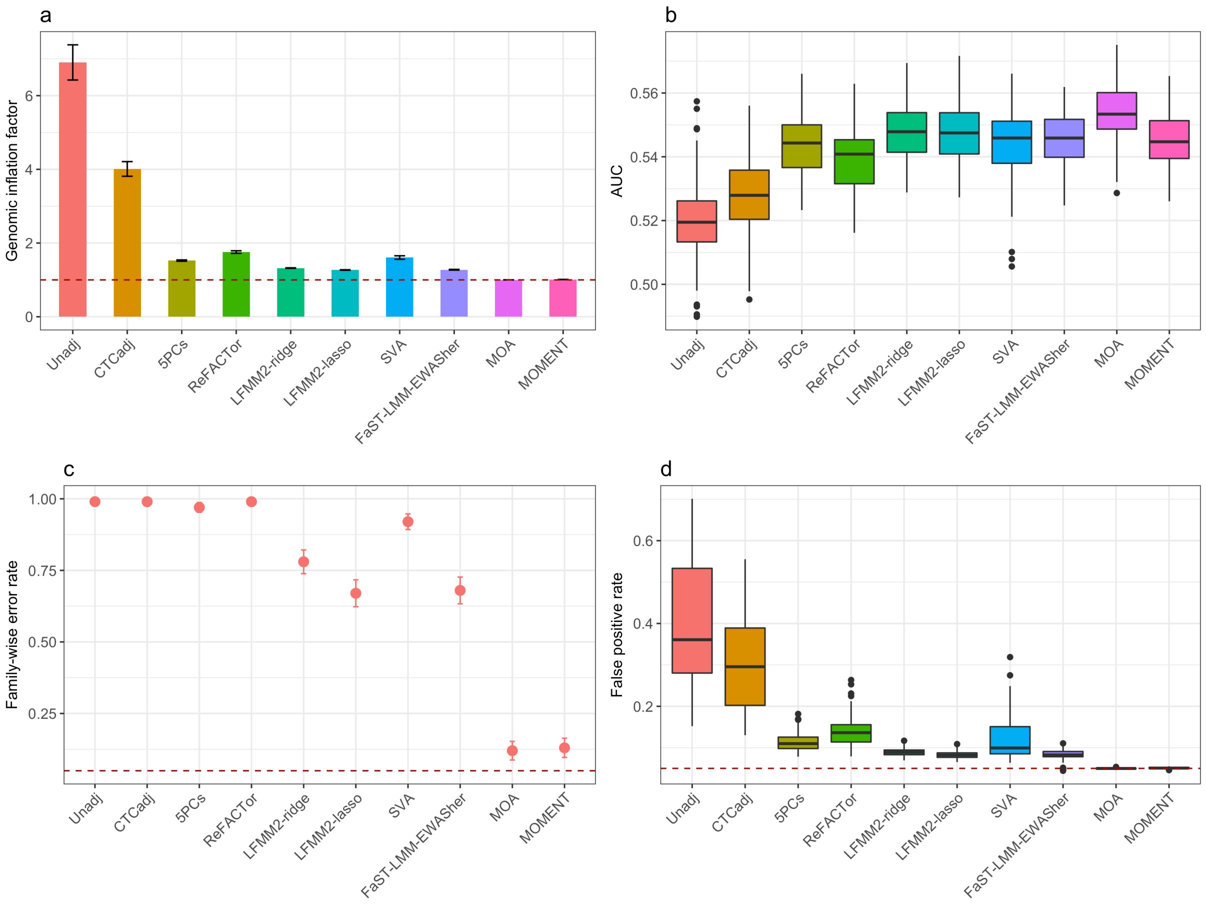


**Supplementary Figure 11** Power and false positive rate for the MWAS methods in simulation scenario 3 (phenotypes simulated based on effects from 1000 causal probes and 5 CTCs). See **Supplementary Figure 3** for details of the labels.

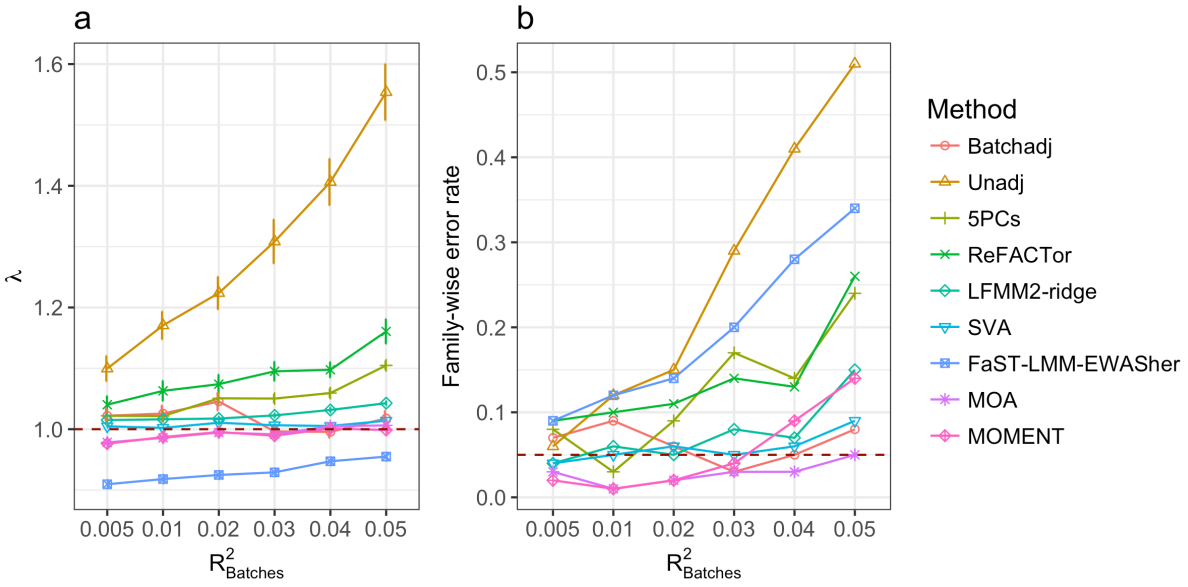


**Supplementary Figure 12** Genomic inflation factor and family-wise error rate for the MWAS methods in simulation scenario 2 (phenotypes simulated based on the effects of 22 batches without causal probes). Shown on the horizontal axis is the proportion of variance in the simulated phenotype explained by the batch effects ($R^2_{\text{batch}}$). (a) Each dot represents the mean $\lambda$ value from 100 simulation replicates given a specified $R^2_{\text{batch}}$ value for a method with an error bar representing +/- the SE of the mean. (b) Each dot represents the family-wise error rate.

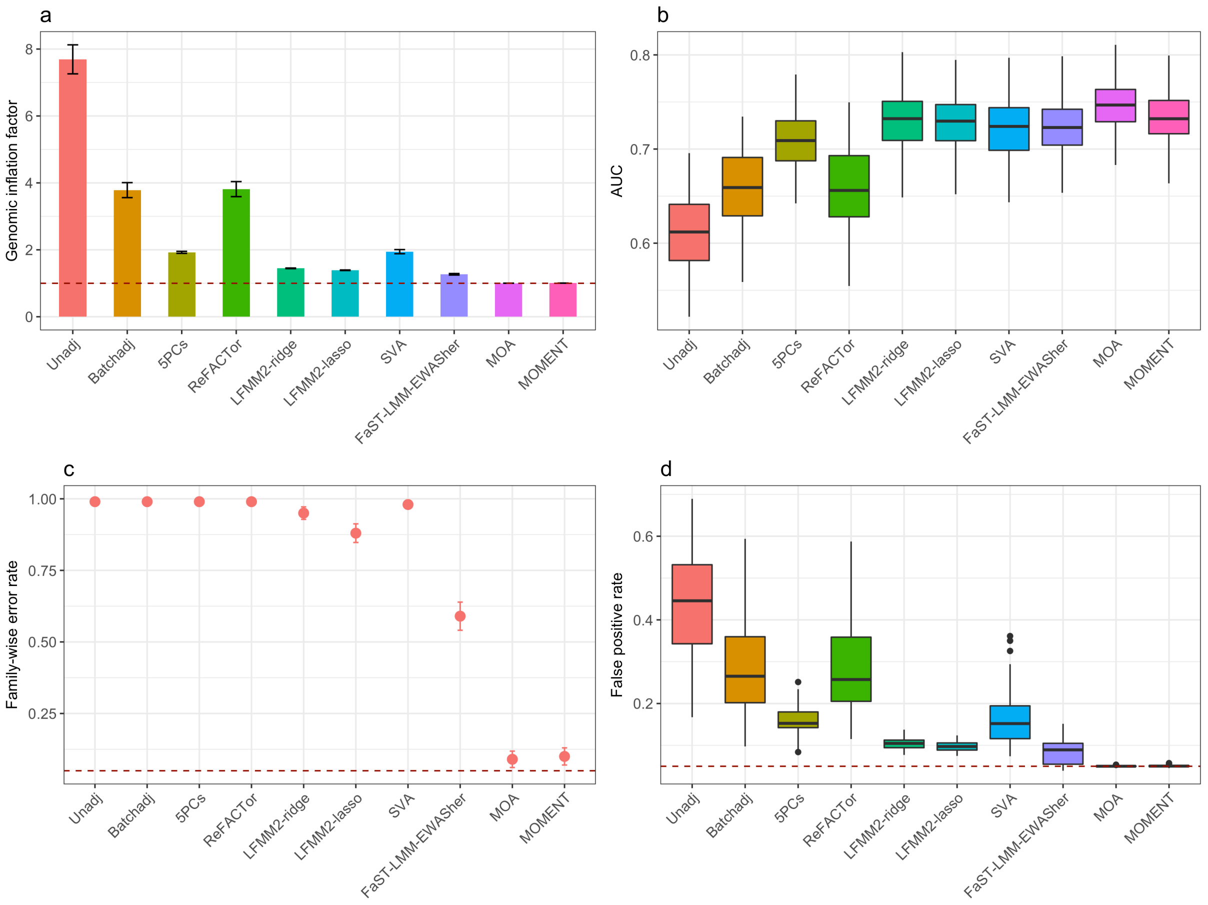


**Supplementary Figure 13** Power and false positive rate for the MWAS methods in simulation scenario 3 (phenotypes simulated based on effects from 100 causal probes and 22 batches). See **Supplementary Figure 3** for details of the labels.

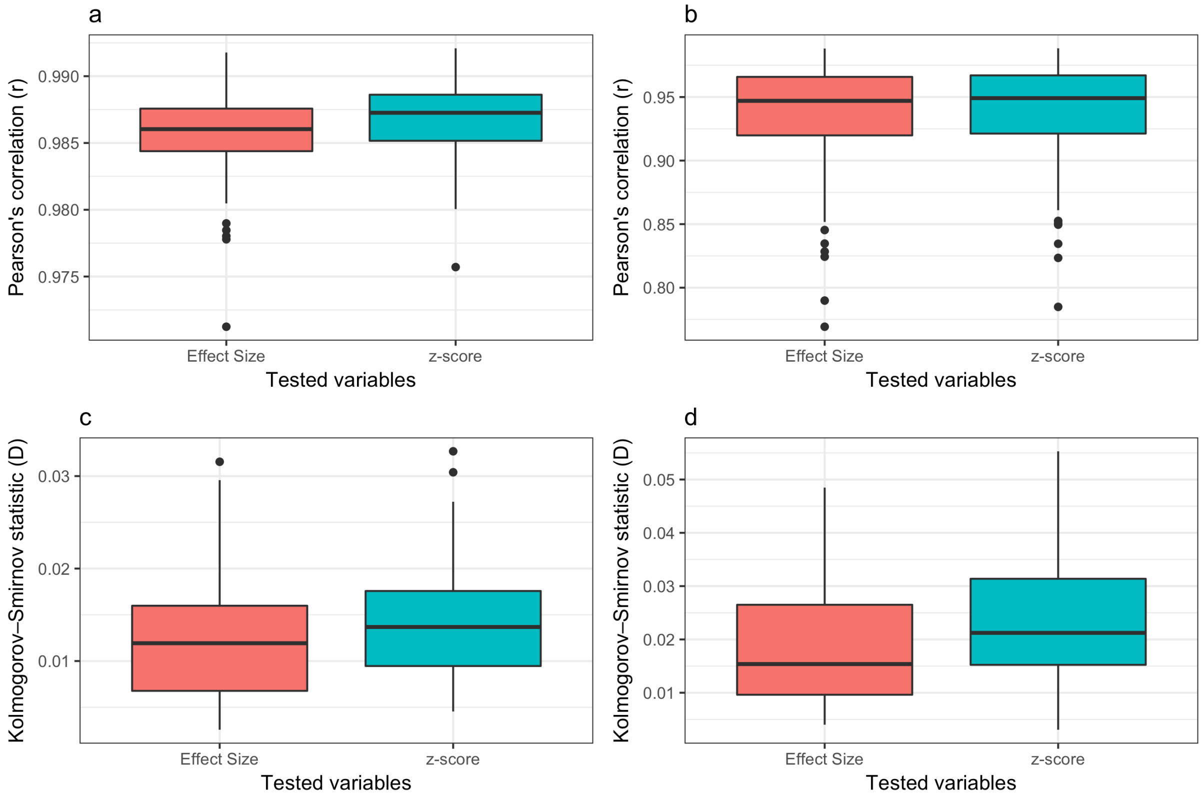


**Supplementary Figure 14** Comparison of the result from MOA/MOMENT analysis of the whole sample with that from a meta-analysis of summary statistics from MOA/MOMENT analyses of LBC1936 and LBC1921 across 100 simulation replicates. (a) Pearson correlation between the result from MOA of the whole sample and the meta-analysis of summary statistics from MOA. (b) Pearson correlation between the result from MOMENT of the whole sample and the meta-analysis of summary statistics from MOMENT. (c) Kolmogorov–Smirnov distance between the result from MOA of the whole sample and the meta-analysis of summary statistics from MOA. (d) Kolmogorov–Smirnov distance between the result from MOMENT of the whole sample and the meta-analysis of summary statistics from MOMENT.

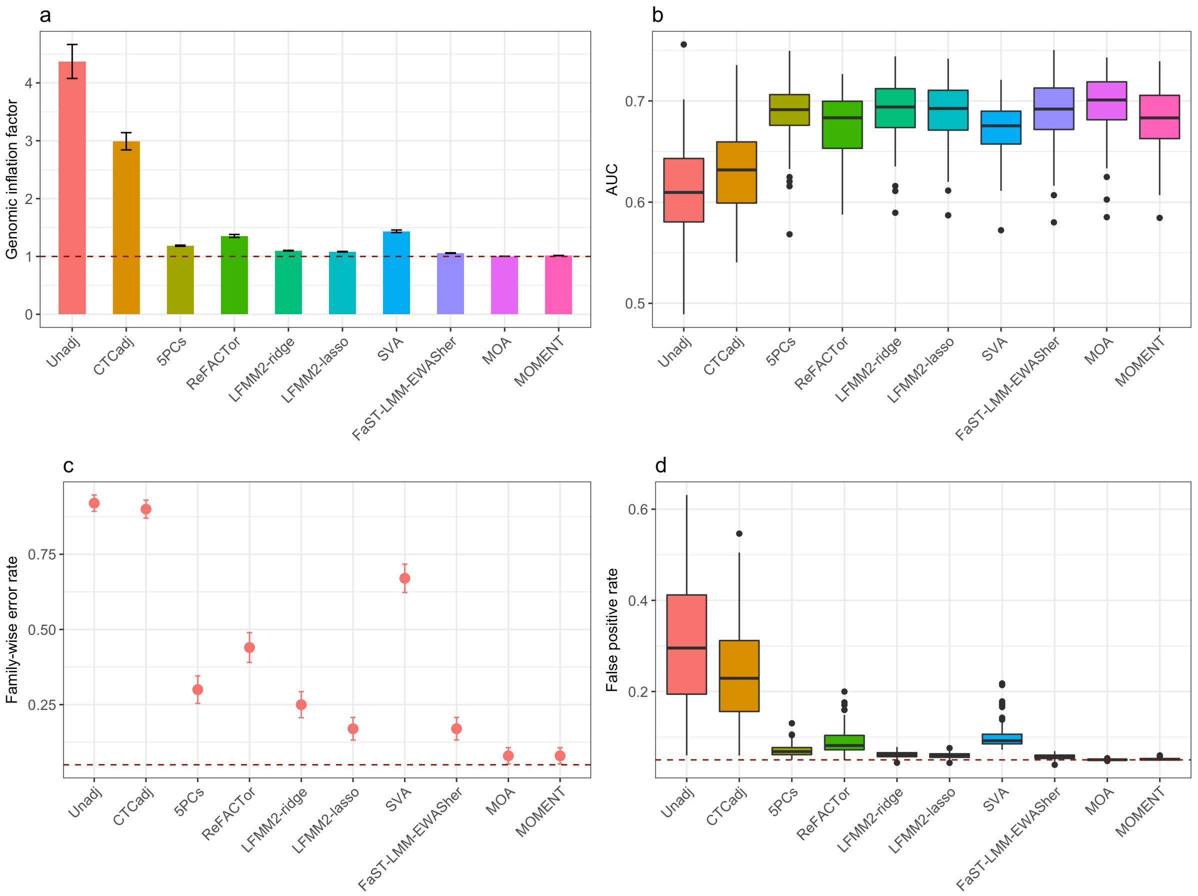


**Supplementary Figure 15** Power and false positive rate for the MWAS methods in application to case-control phenotypes. We simulated a quantitative trait based on the same setting as in simulation scenario 3 (**Supplementary Note 1**). In each simulation replicate, we ranked the individuals from the largest to the smallest by their simulated phenotypes, and dichotomised the trait by selecting the top 30% of the individuals as cases and the other individuals as controls.

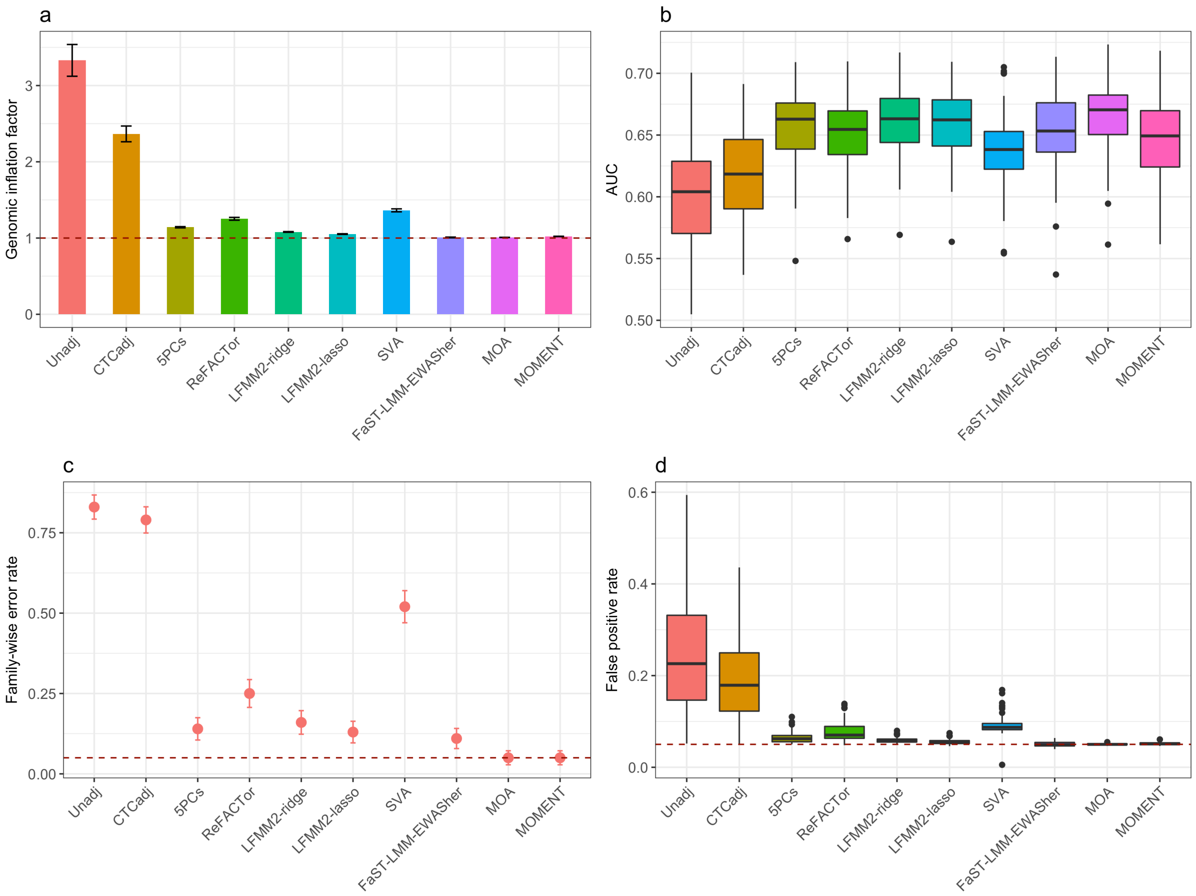


**Supplementary Figure 16** Power and false positive rate for the MWAS methods in application to case-control phenotypes with oversampled cases. We simulated a quantitative trait using the same setting as in simulation scenario 3 (**Supplementary Note 1**) and ranked the individuals from the largest to the smallest by their simulated phenotypes. We selected the top 30% of the individuals as cases and randomly selected the same number of individuals from the rest of the sample as controls.

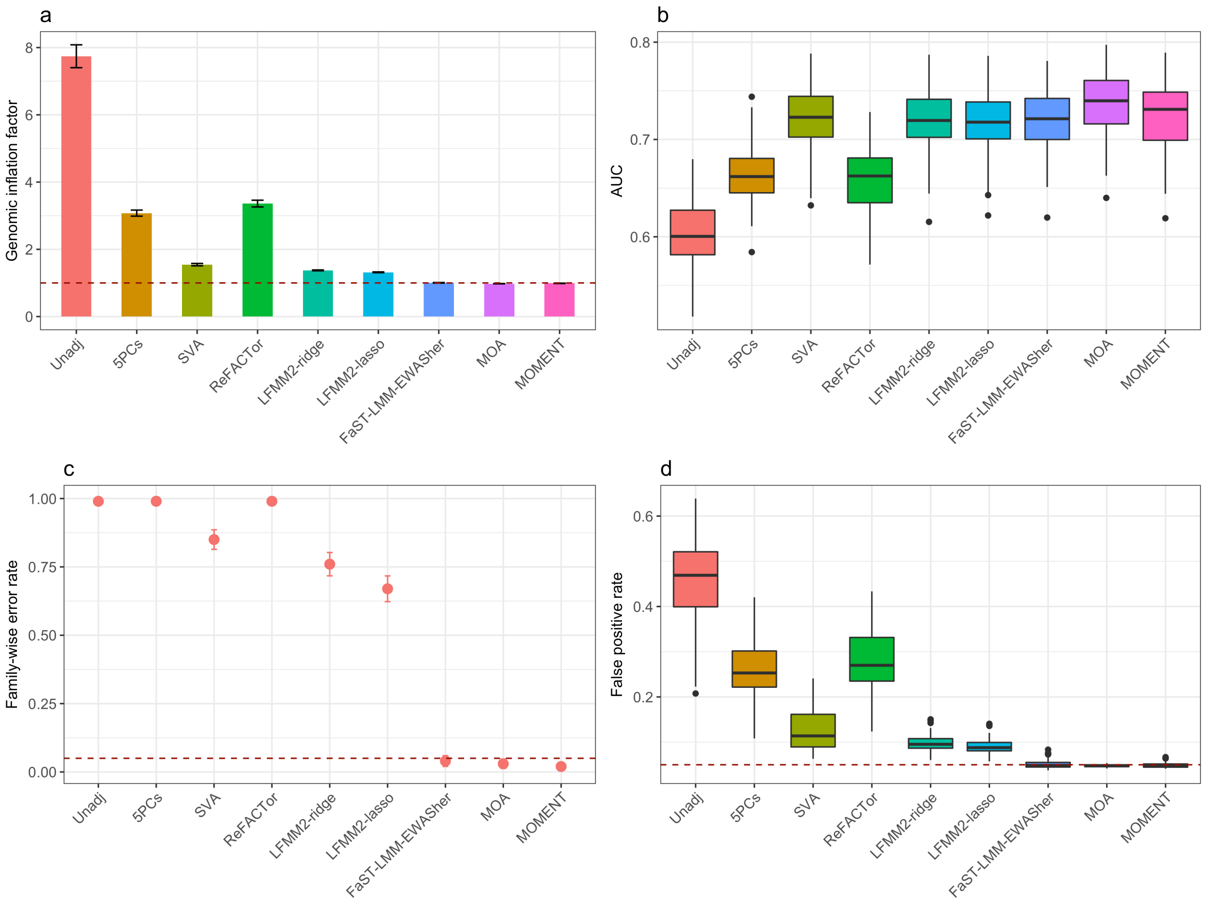


**Supplementary Figure 17** Power and false positive rate for the TWAS methods in application to gene expression data. We simulated a transcriptome-wide association study (TWAS) based on the SAFHS gene expression data, 19,648 probes on 1,219 individuals (**Methods**). The gene expression measures were adjusted for age and sex, leaving smoking status as one of the unmodeled confounding factors. We simulated a quantitative trait by randomly sampling 100 gene expression probes on the odd chromosomes as causal probes ($\rho^2 = 0.5$) using the same strategy as used in simulation scenario 1 and repeated the simulation 100 times.

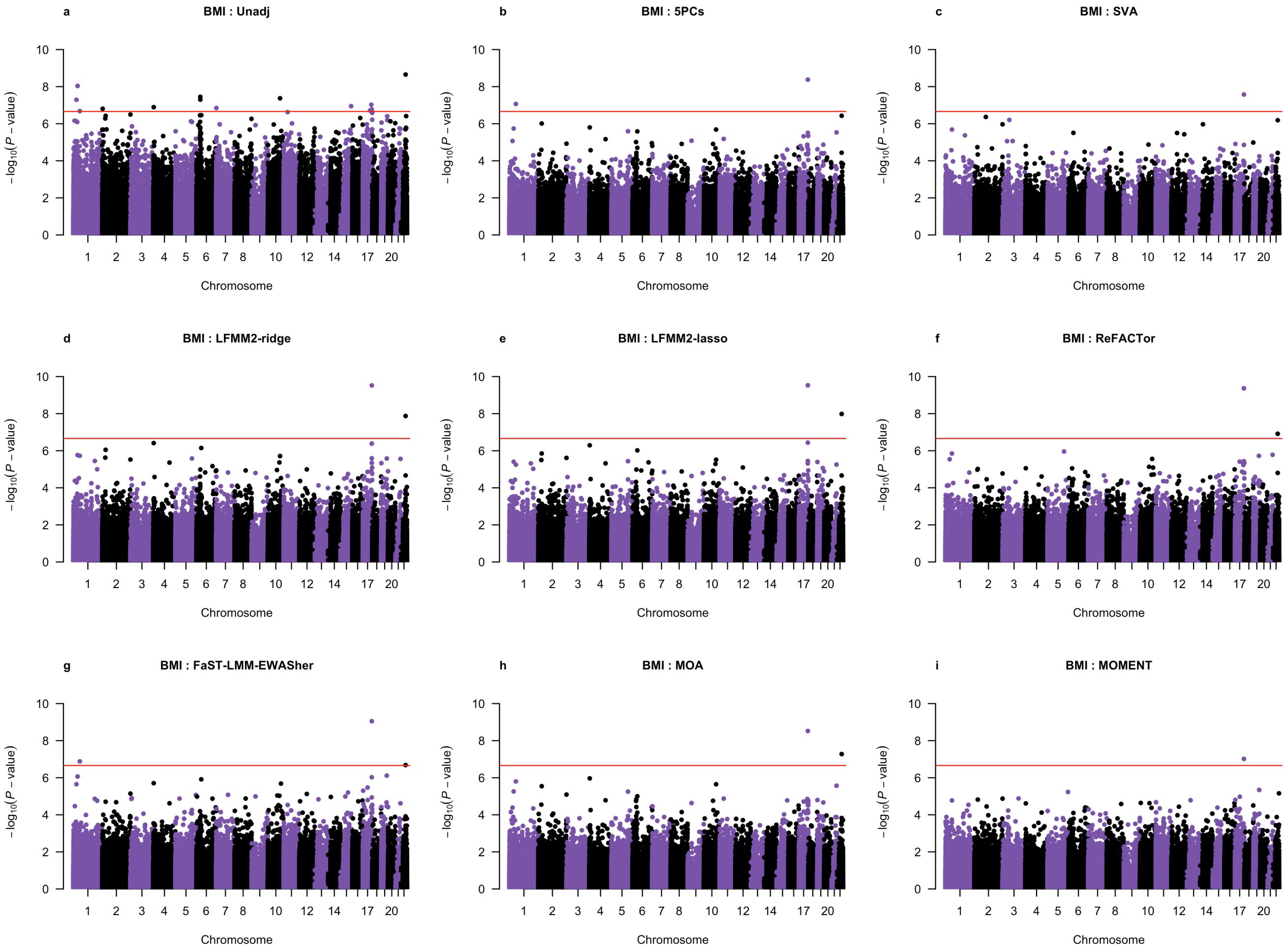


**Supplementary Figure 18** Manhattan plots of association p-values from 9 MWAS methods for BMI in the LBC. The red horizontal line represents the genome-wide significance level.

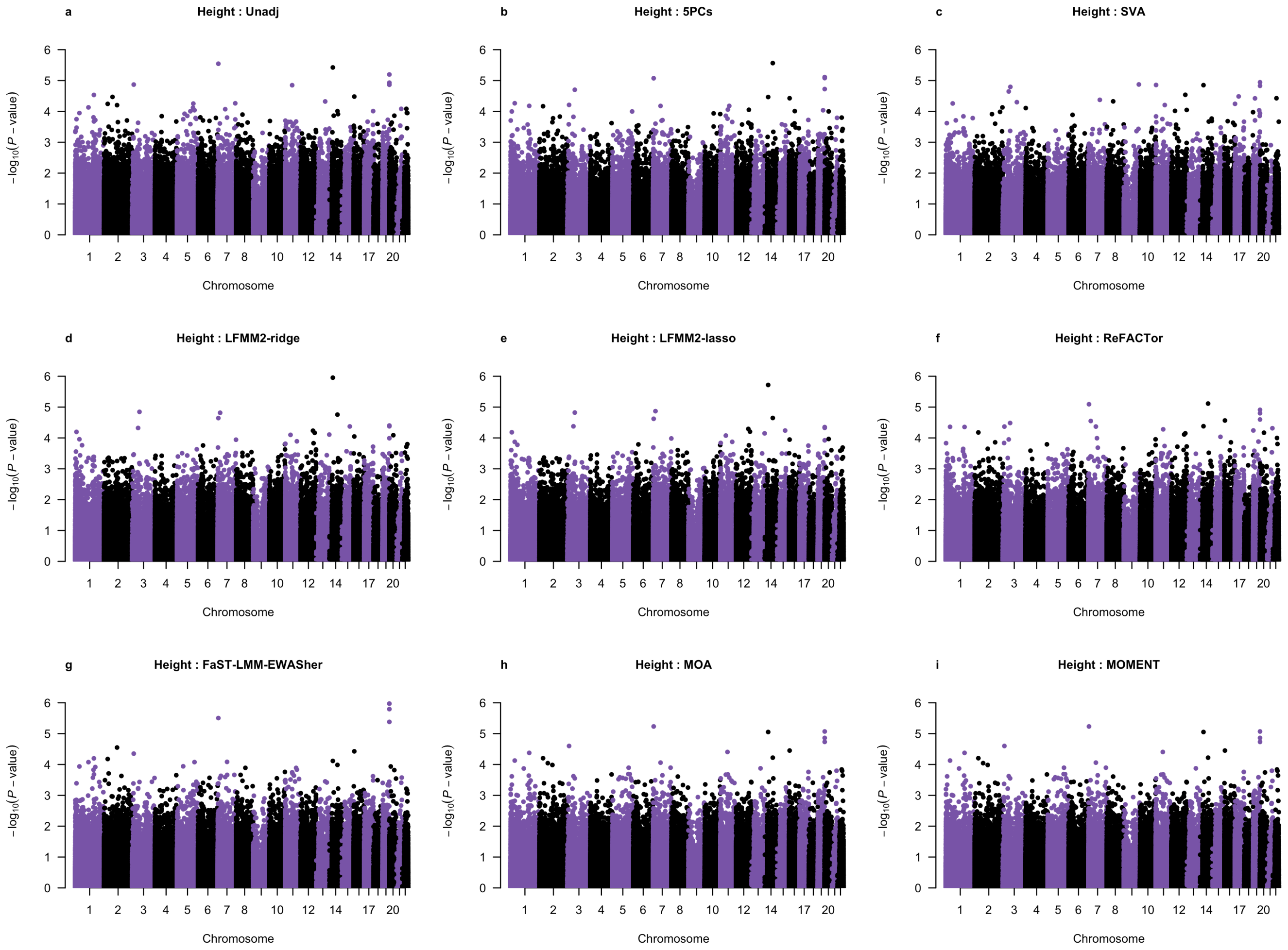


**Supplementary Figure 19** Manhattan plots of association p-values from 9 MWAS methods for height in the LBC.

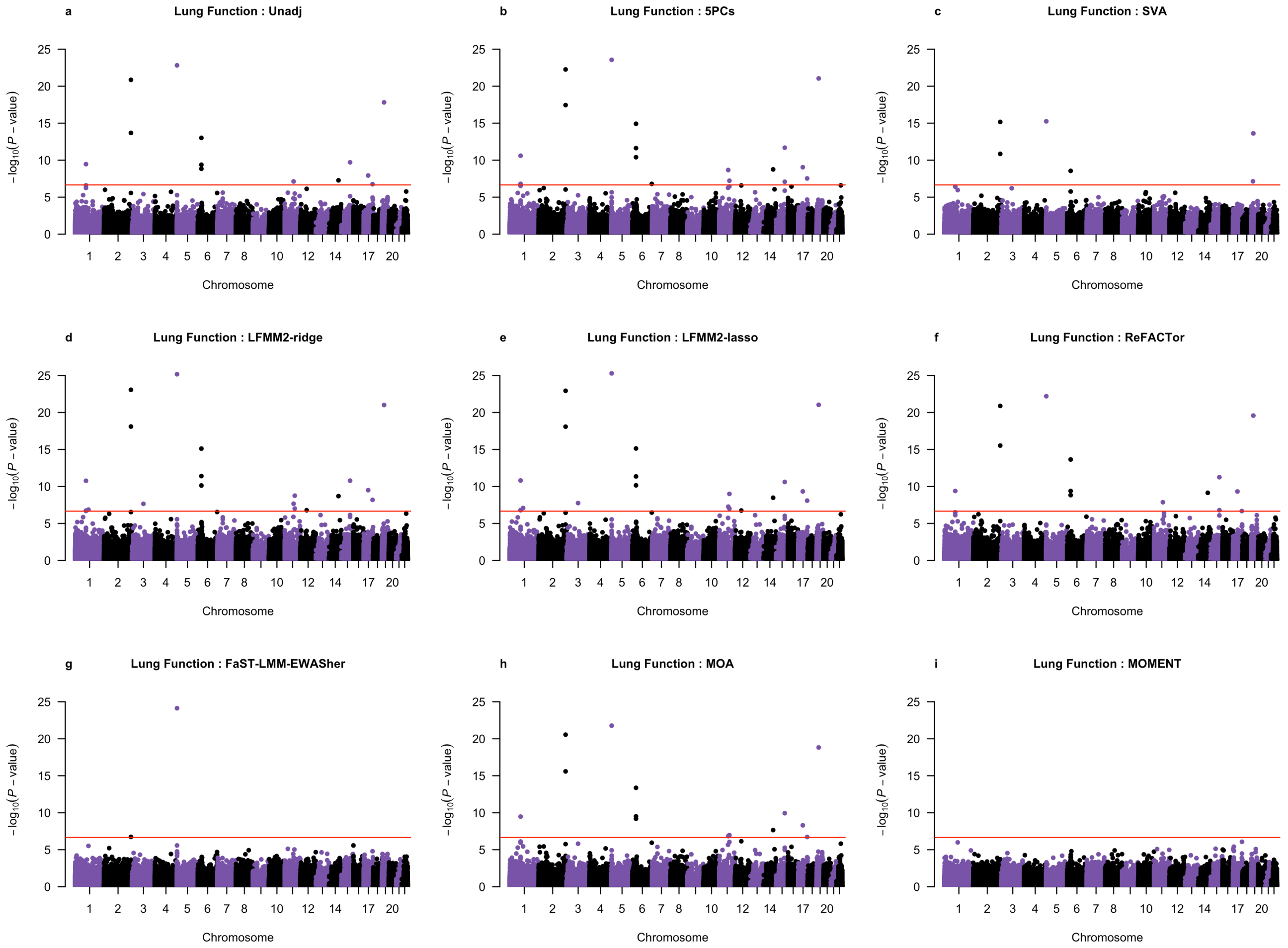


**Supplementary Figure 20** Manhattan plots of association p-values from 9 MWAS methods for lung function in the LBC.

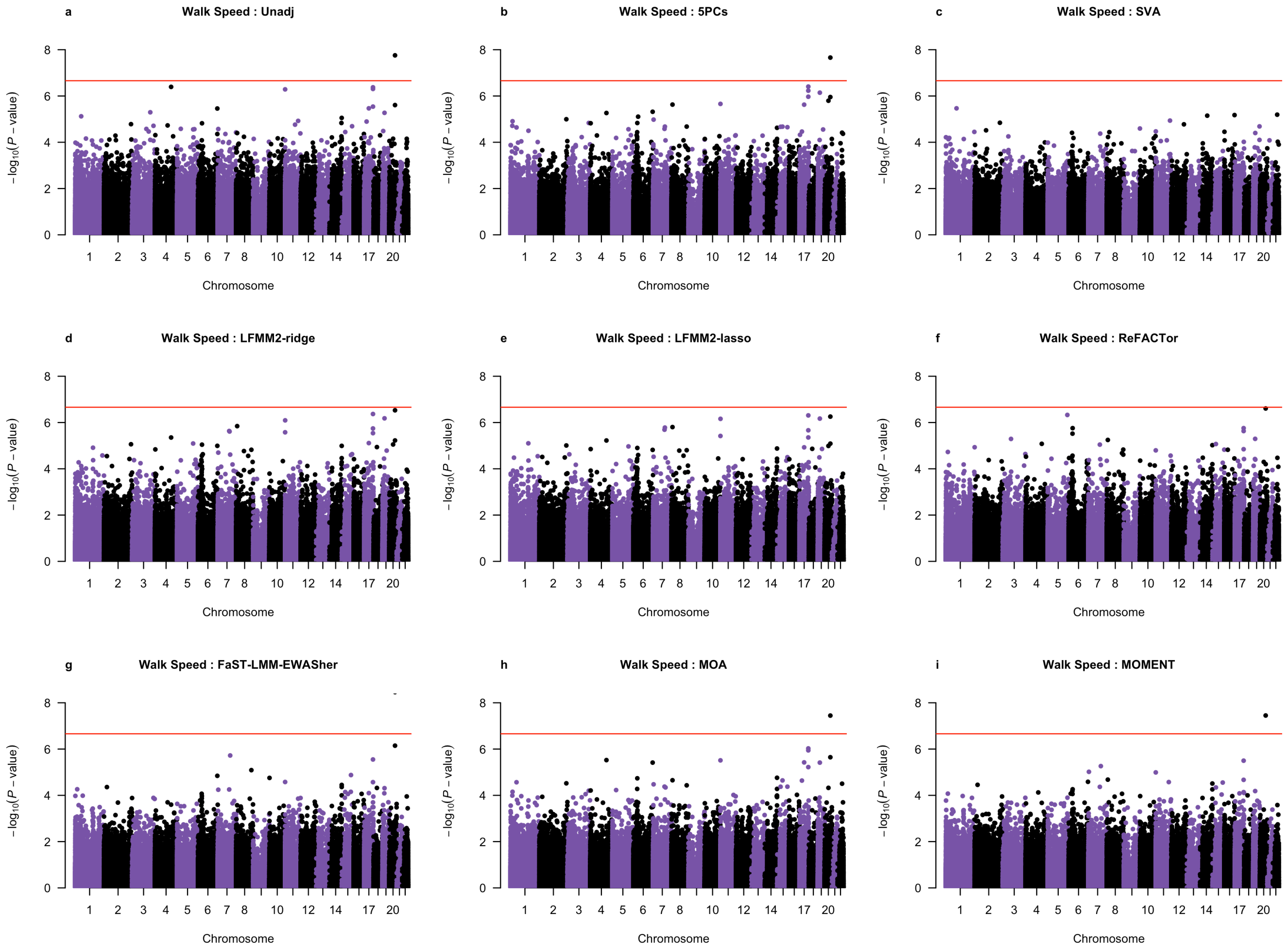


**Supplementary Figure 21** Manhattan plots of association p-values from 9 MWAS methods for walking speed in the LBC.

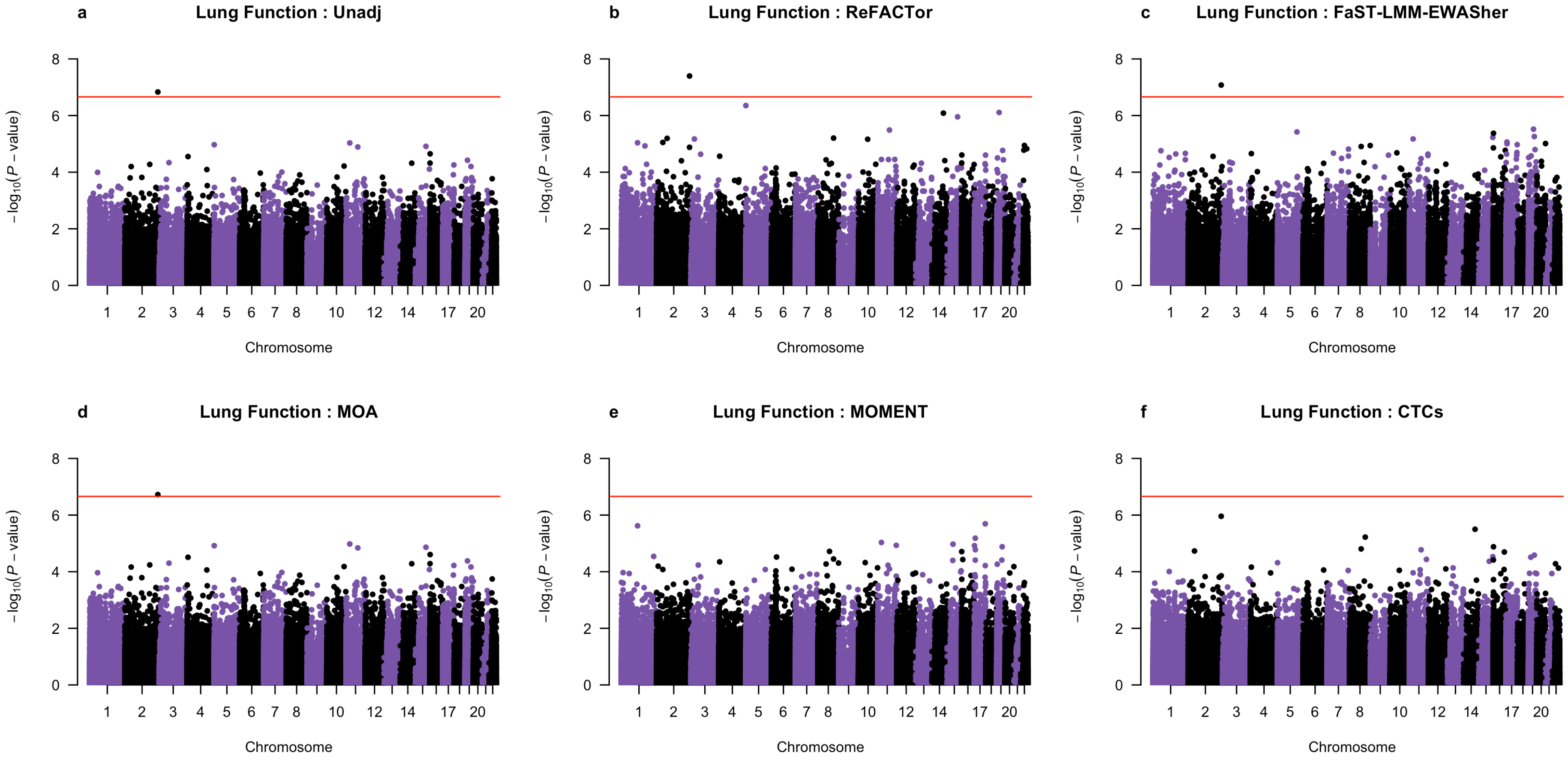


**Supplementary Figure 22** Manhattan plots of association p-values from 6 MWAS methods for lung function with smoking status fitted as a fixed covariate in the LBC.

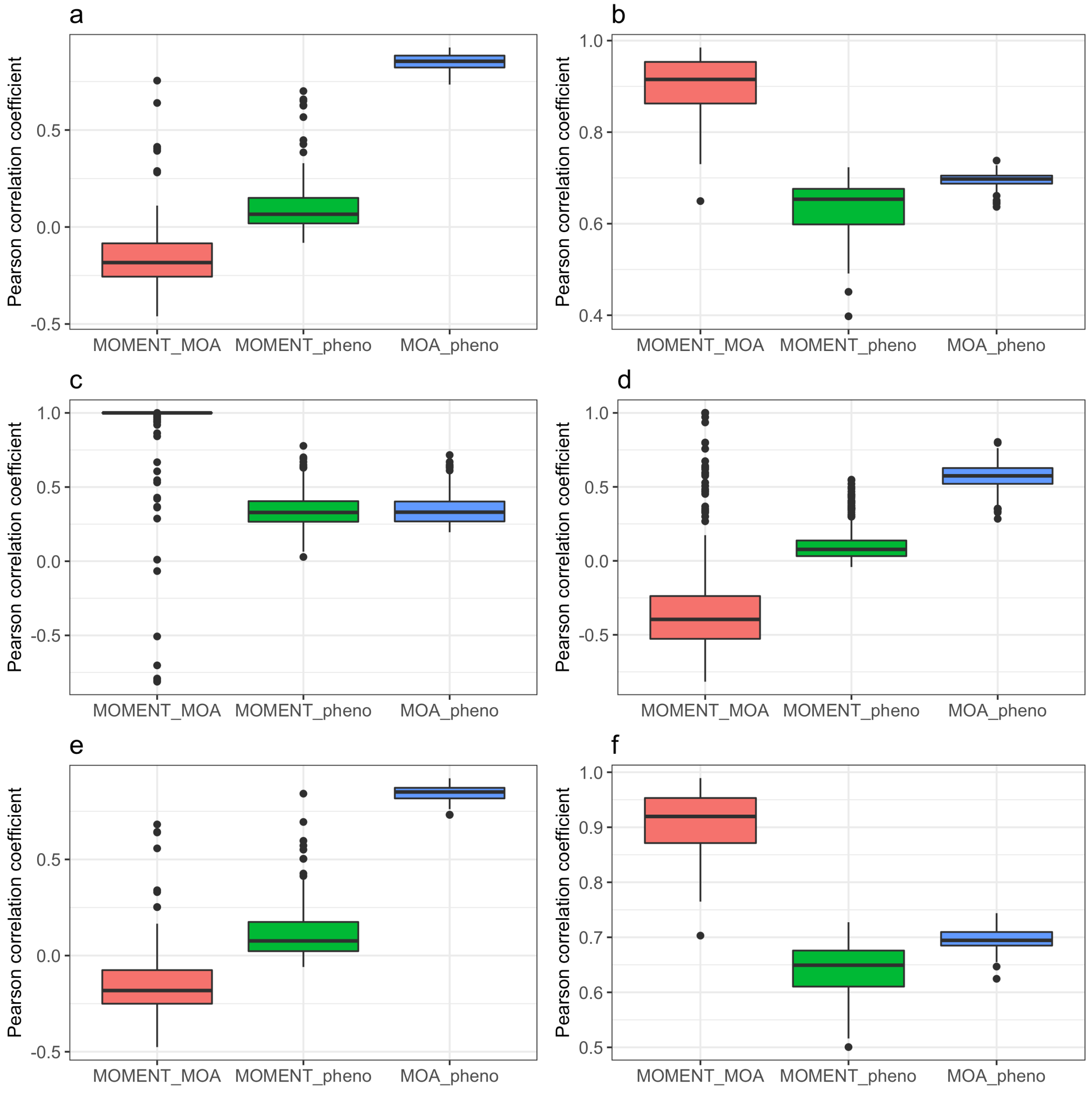


**Supplementary Figure 23** MWAS scores computed from null or causal probes for the simulated phenotypes in the LBC. MWAS score = $\sum_i w_i \hat{b}_i$ where $w_i$ is the standardised DNAm measure of the $i$-th probe with its effect $b_i$ estimated from MOMENT or MOA. Shown are the correlation between MOMENT and MOA MWAS scores, and the correlations of MOMENT and MOA scores with the phenotype. For the null probes, the MOMENT scores were much less correlated with the phenotype compared to the corresponding MOA scores, suggesting that MOMENT is more efficient at correcting for the correlation between the null and causal probes than MOA, which could lead to a negative correlation between MOMENT and MOA scores. (a) null probes from simulation scenario 1 (100 causal probes with $\rho^2 = 0.5$), (b) causal probes from simulation scenario 1, (c) null probes from simulation scenario 2 ($R^2_{\text{CTCs}} = 0.005$), (d) null probes from simulation scenario 2 ($R^2_{\text{CTCs}} = 0.3$), (e) null probes from simulation scenario 3 ($R^2_{\text{CTCs}} = 0.05$; 100 causal probes with $\rho^2 = 0.5$), (f) causal probes from simulation scenario 3.

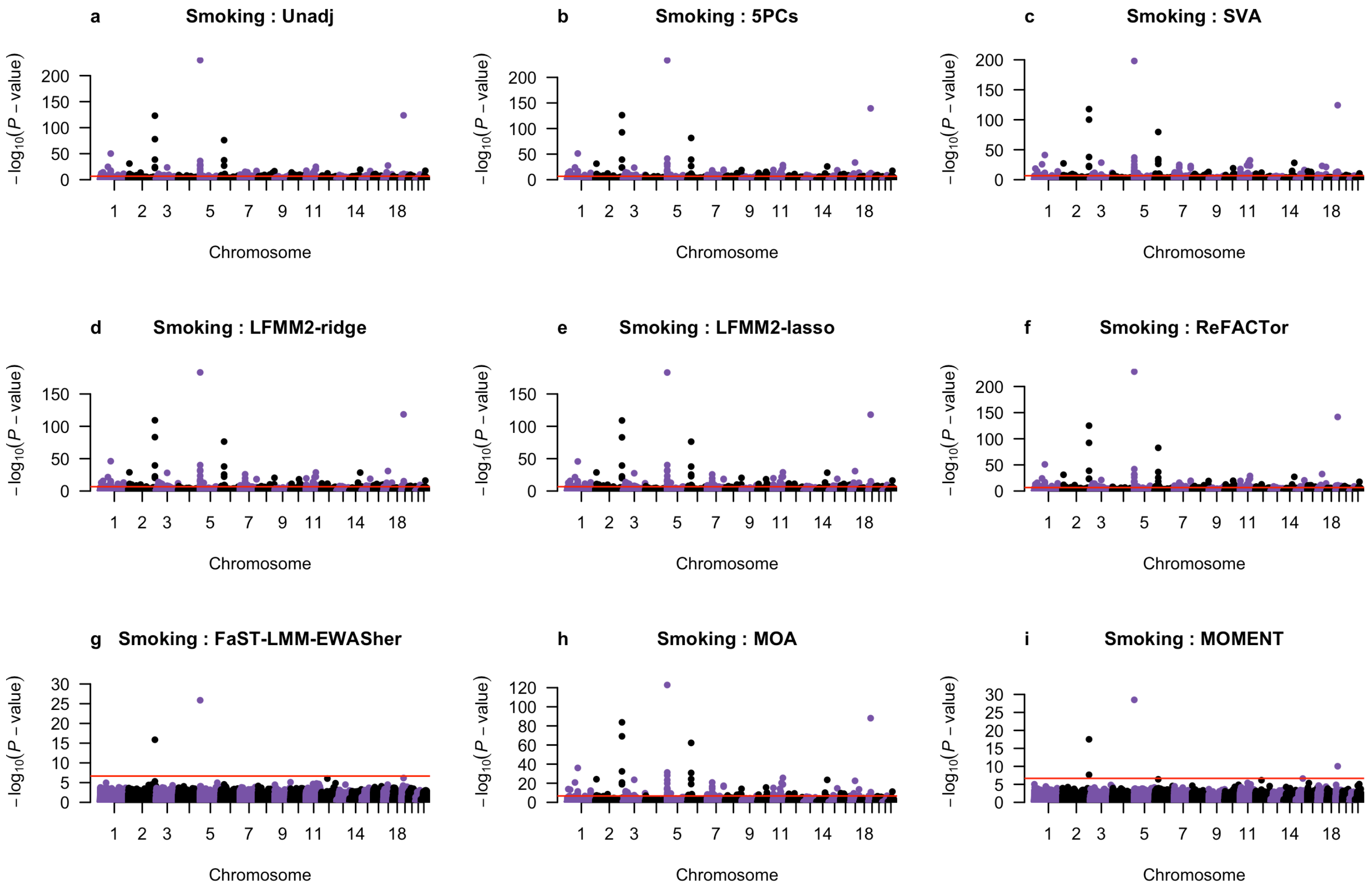


**Supplementary Figure 24** Manhattan plots of association p-values from 9 MWAS methods for a categorical phenotype of smoking status (coded as 0, 1 and 2 for non-smoker, former smoker and current smoker) in the LBC.

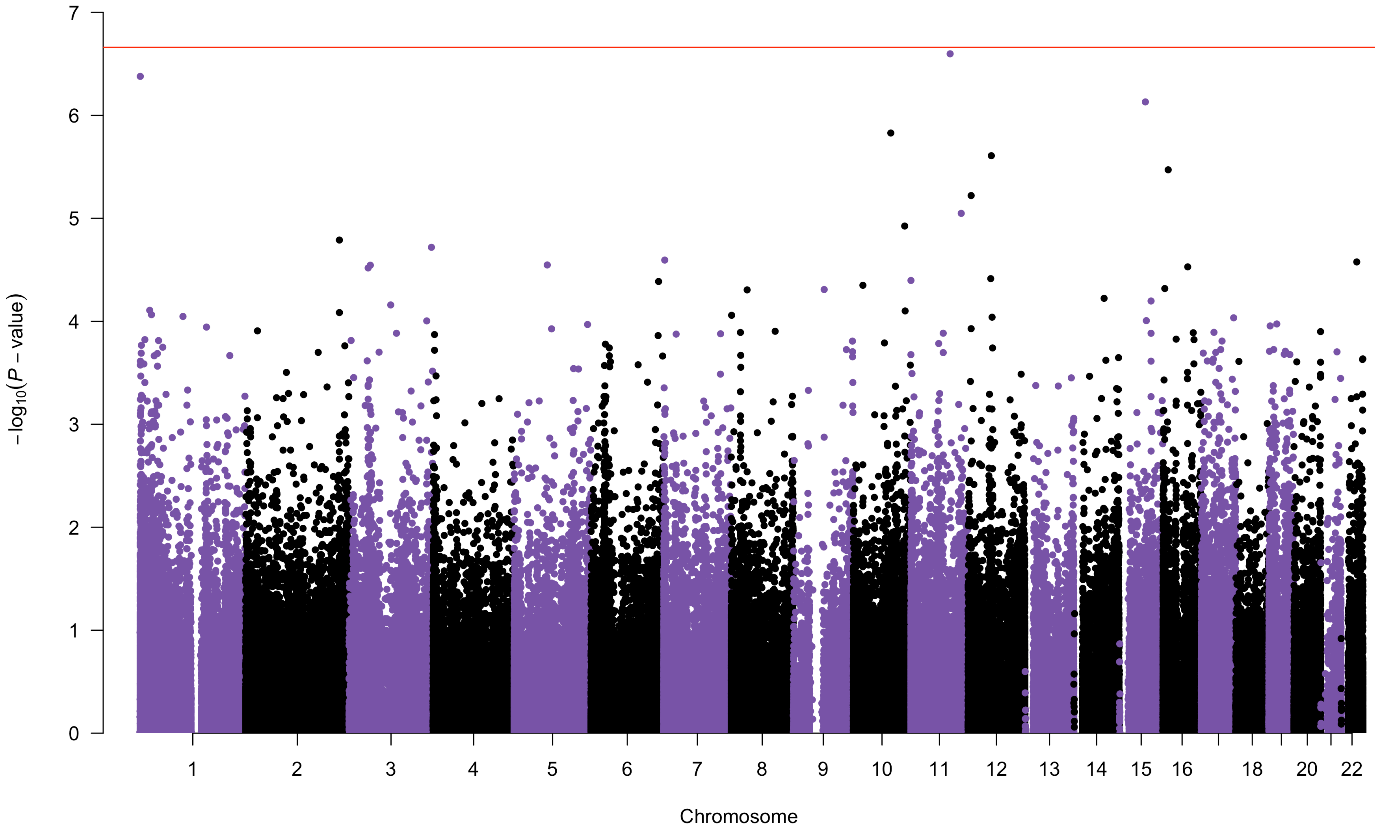


**Supplementary Figure 25** Manhattan plot of association p-values from MOA for a categorical phenotype (coded as 0, 1 or 2) of smoking status with the top 4 MOMENT signals fitted as fixed covariates in the LBC.

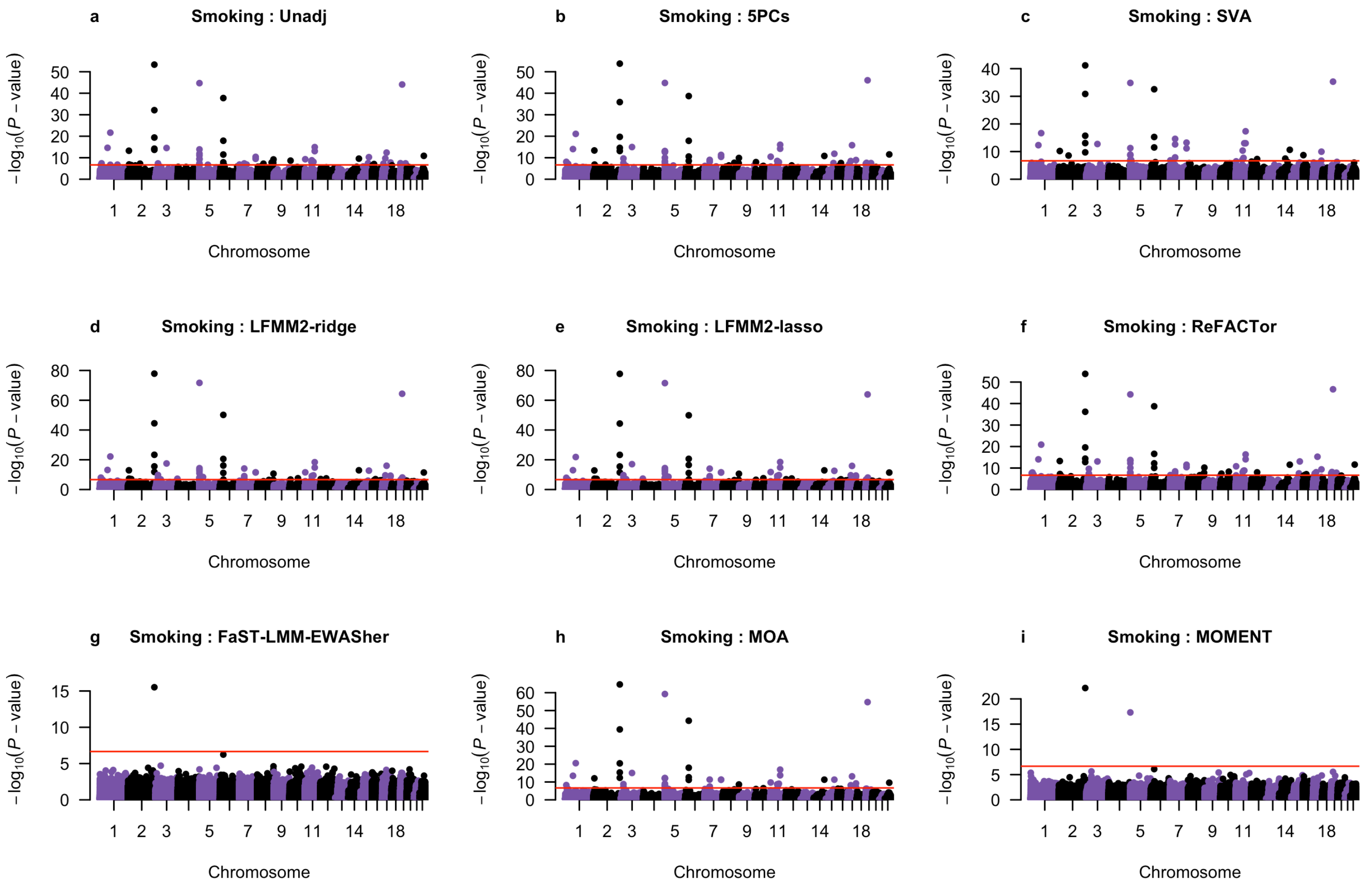


**Supplementary Figure 26** Manhattan plots of association p-values from 9 MWAS methods for a binary phenotype of smoking status (coded as 0 and 1 for non-smoker and former smoker or current smoker) in the LBC.

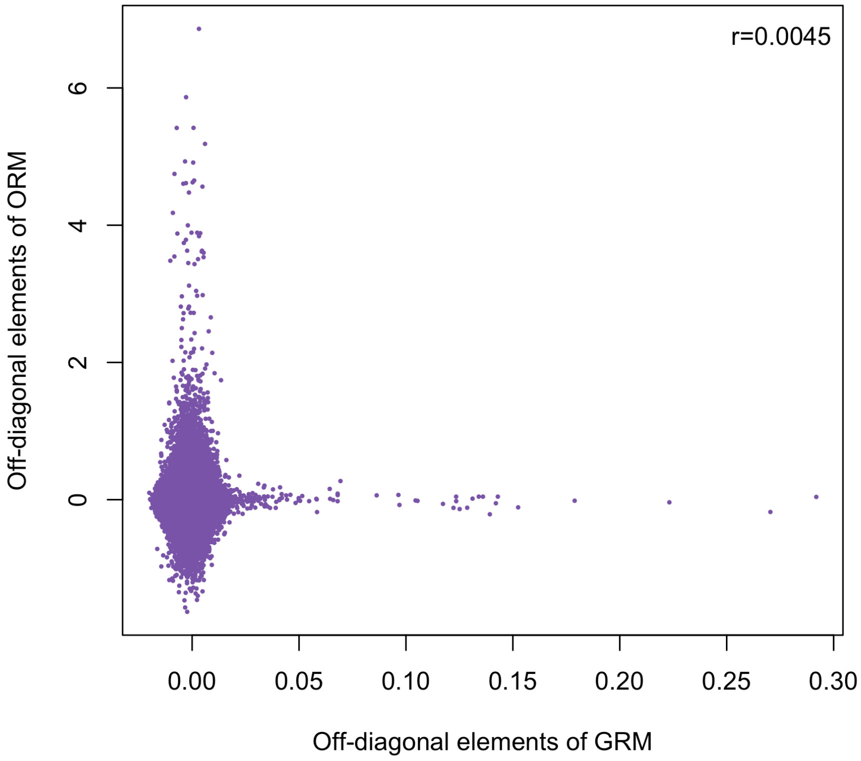


**Supplementary Figure 27** Correlation of the off-diagonal elements between GRM and ORM computed from genetic and DNAm data in the LBC.

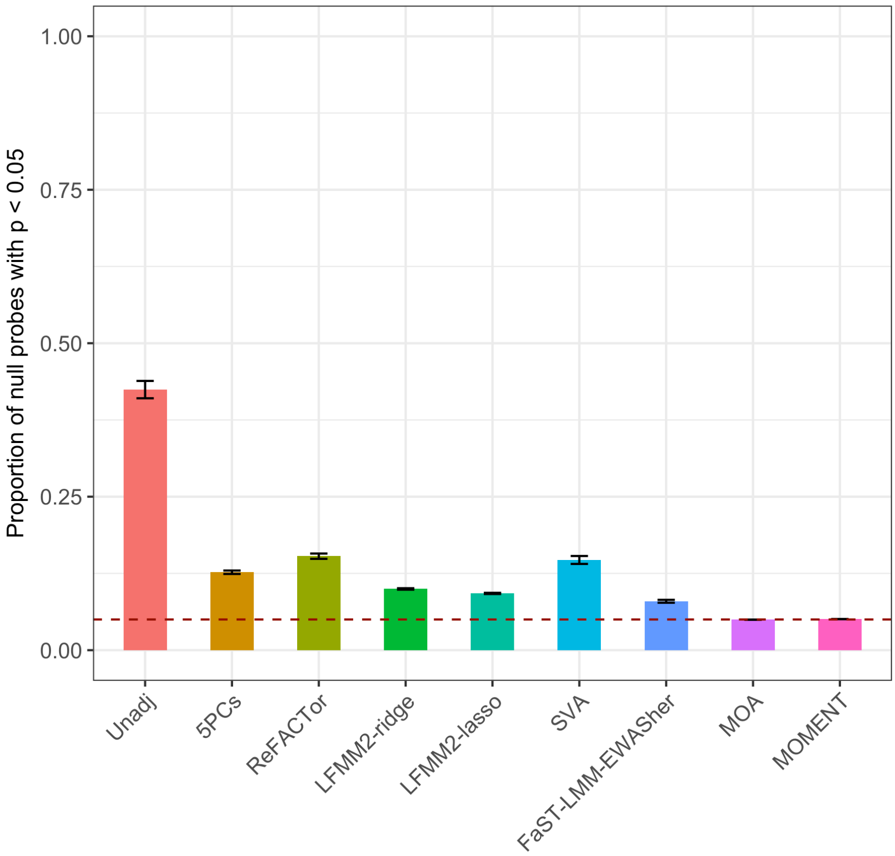


**Supplementary Figure 28** Proportion of null probes with $P_{\mathrm{MWAS}} < 0.05$ in simulation scenario 1. Each column represents the mean of proportion of null probes for a method from 100 simulation replicates with an error bar representing +/- SE of the mean.

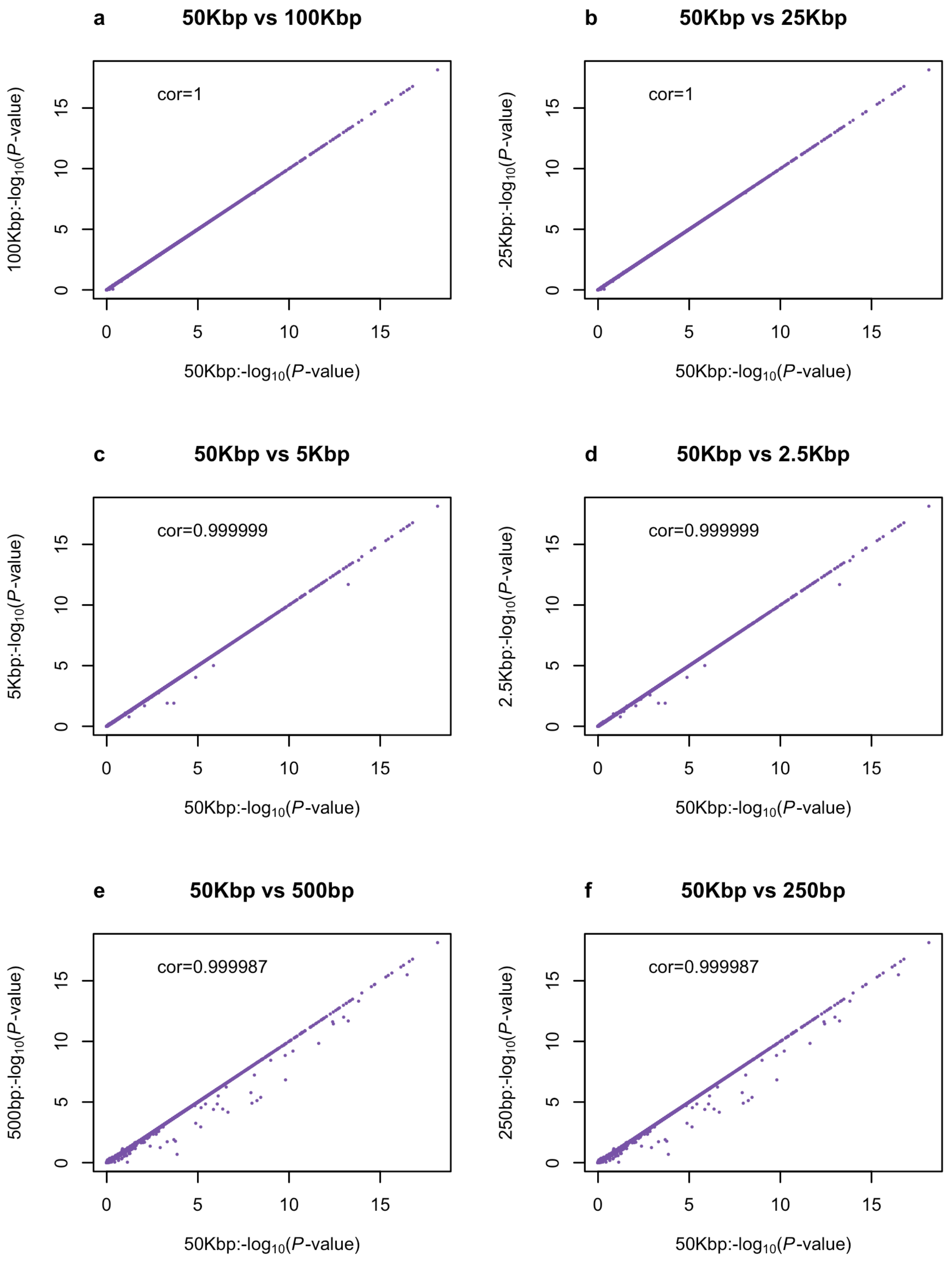


**Supplementary Figure 29** Testing the robustness of MOMENT to window size for DNAm data. We used a window size of 50Kbp (a total region of ~100Kbp) as the benchmark for comparison. (a) 100Kbp (b) 25Kbp (c) 5Kbp (d) 2500bp (e) 500bp and (f) 250bp.

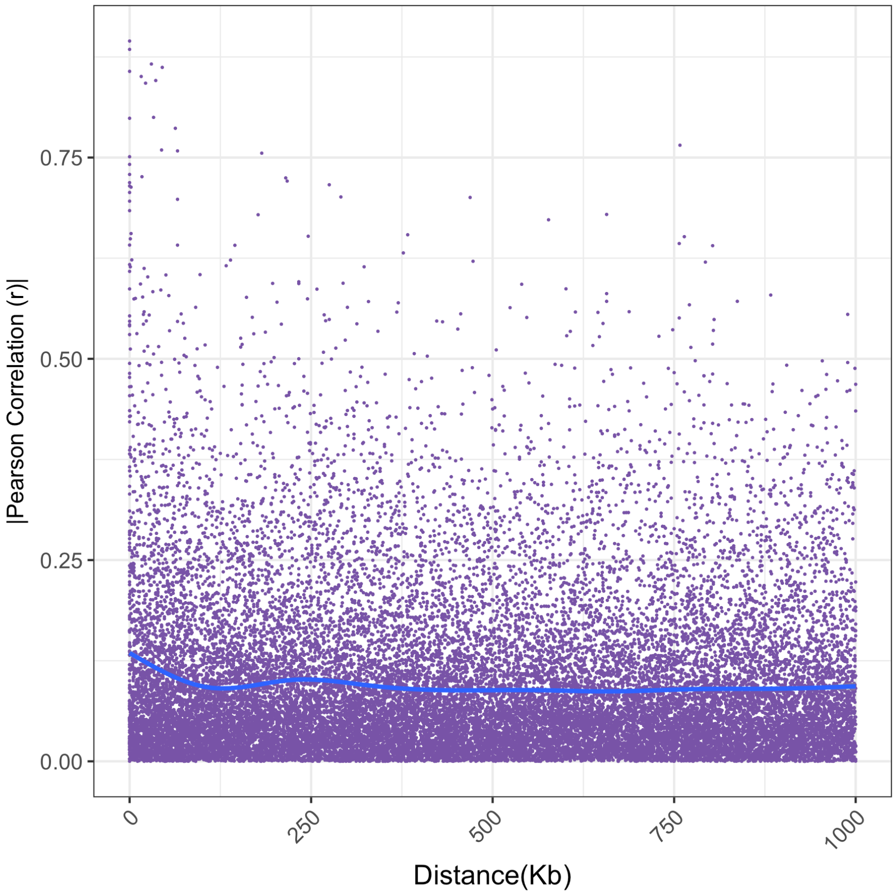


**Supplementary Figure 30** Decay of Pearson correlation between gene expression probes as a function of their physical distance.

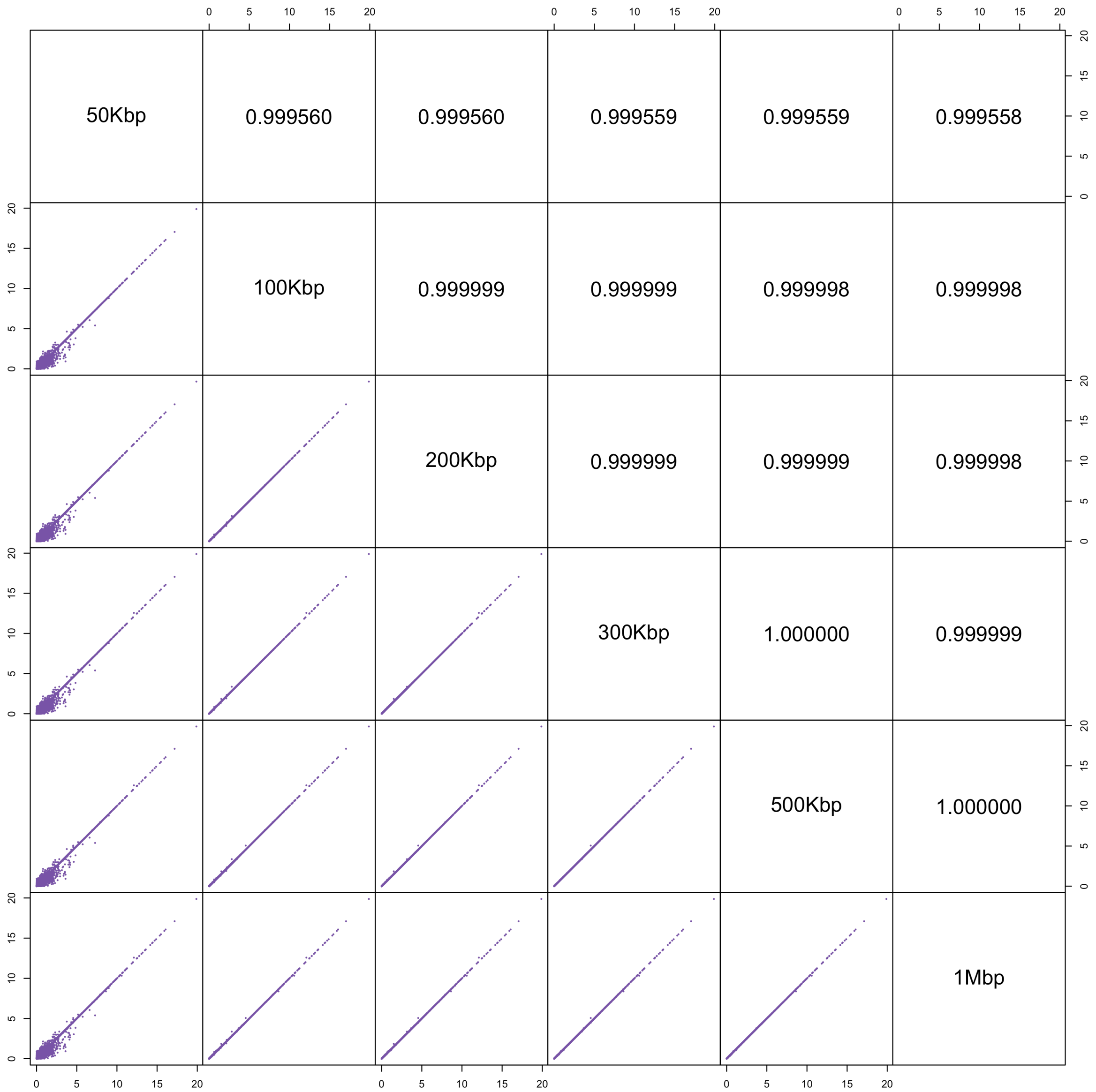


**Supplementary Figure 31** Testing the robustness of MOMENT to window size for gene expression data. We varied the window size from 50Kbp to 100Kbp, 200Kbp, 300Kbp, 500Kbp and 1Mbp. The upper triangular values are Pearson correlation coefficients and the lower triangular figures are the scatter plots of –log10(p-value).

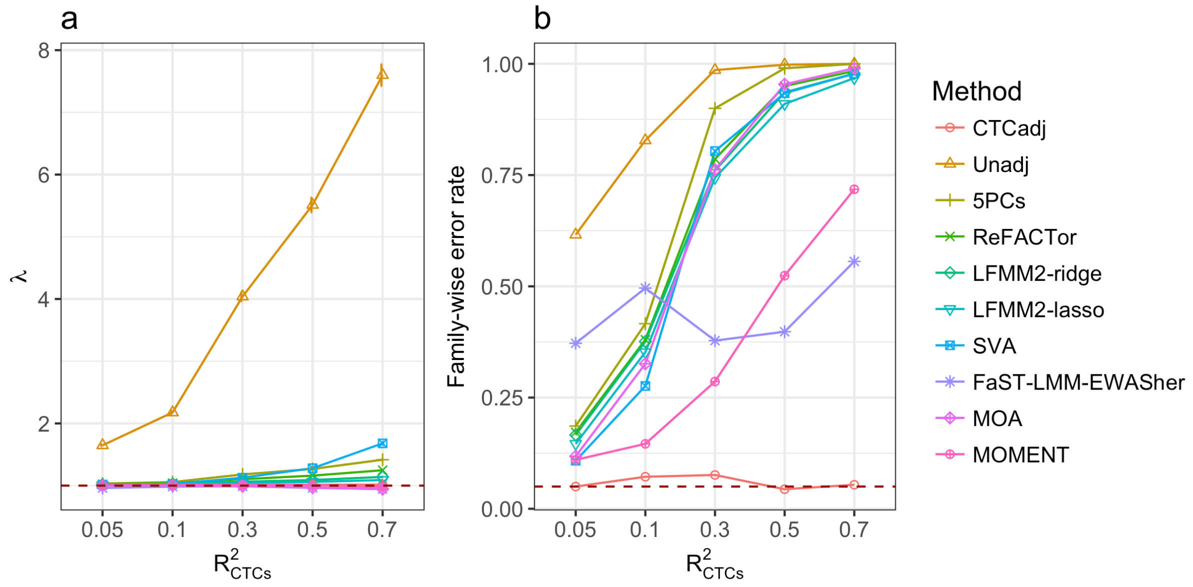


**Supplementary Figure 32** Genomic inflation factor and family-wise error rate for the MWAS methods in simulation scenario 2 with large $R^2_{\mathrm{CTCs}}$ (from 0.05 to 0.7). Shown on the horizontal axis are the $R^2_{\mathrm{CTCs}}$ values used to simulate the phenotype. (a) Each dot represents the mean $\lambda$ value from 1000 simulation replicates given a specific $R^2_{\mathrm{CTCs}}$ value for a method with an error bar representing +/- the SE of the mean. (b) Each dot represents the family-wise error rate.

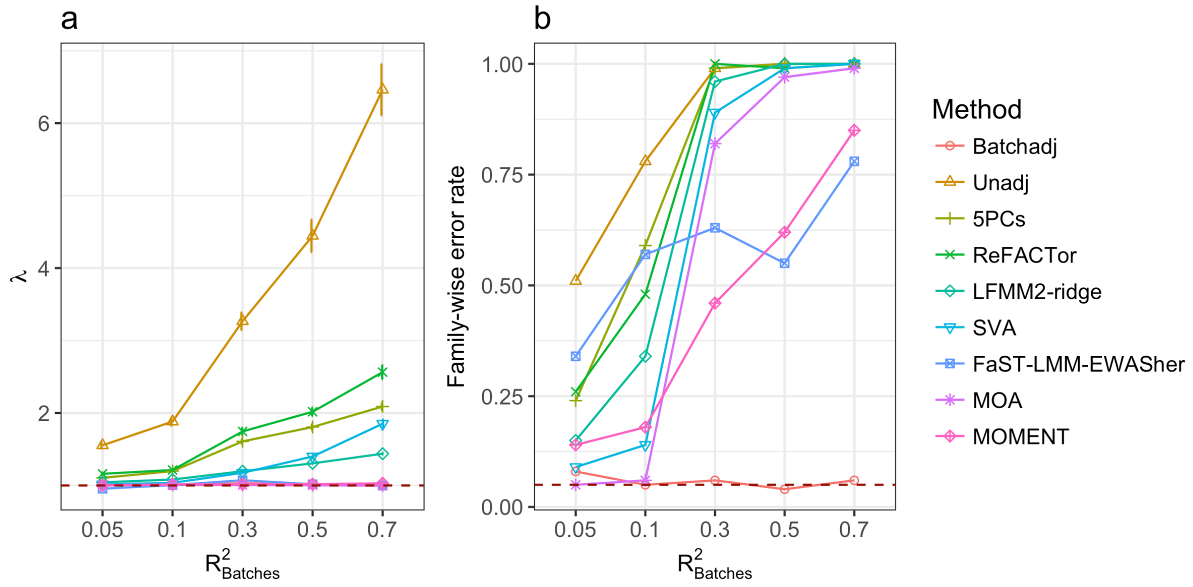


**Supplementary Figure 33** Genomic inflation factor and family-wise error rate for the MWAS methods in simulation scenario 2 with large $R^2_{\text{batch}}$ (from 0.05 to 0.7). Shown on the horizontal axis are the $R^2_{\text{batch}}$ values used to simulate the phenotype. (a) Each dot represents the mean $\lambda$ value from 100 simulation replicates given a specific $R^2_{\text{batch}}$ value for a method with an error bar representing +/- the SE of the mean. (b) Each dot represents the family-wise error rate.

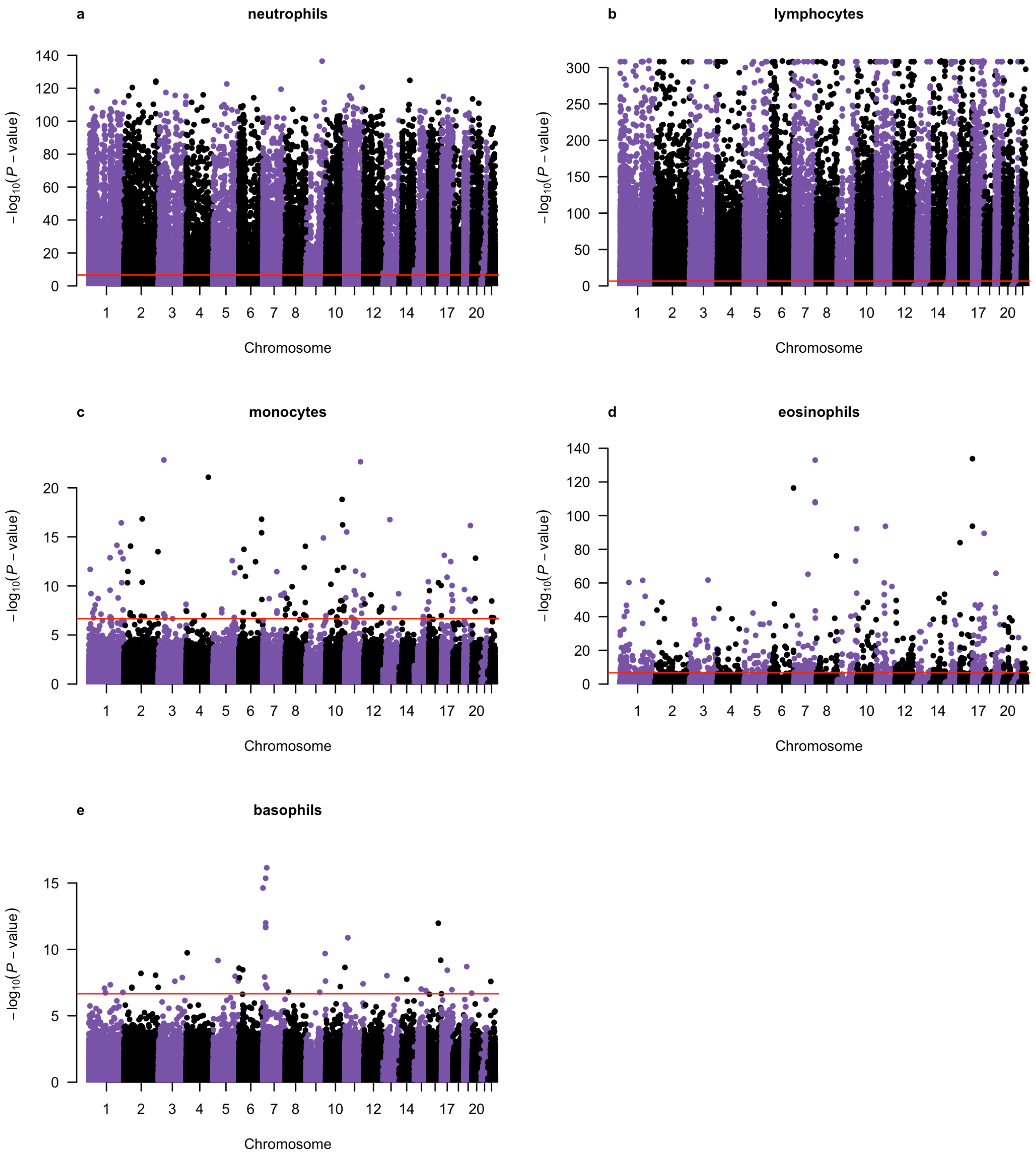


**Supplementary Figure 34** Manhattan plots of association p-values from linear regression for 5 CTCs. DNAm probe measures were adjusted for age, sex, experimental batches, and smoking status.

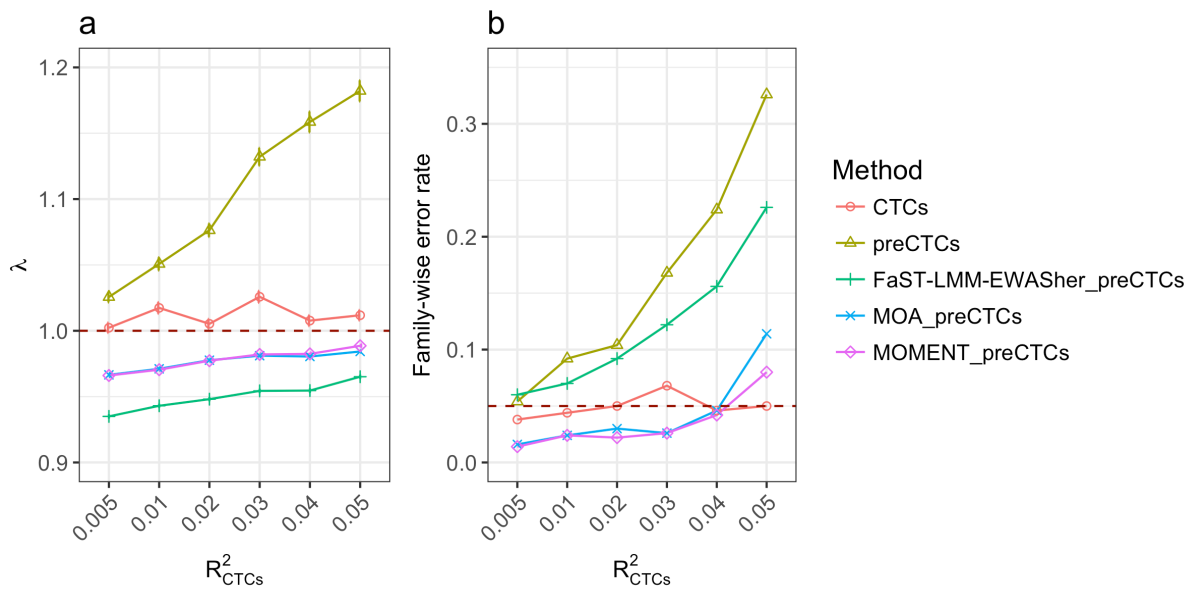


**Supplementary Figure 35** Genomic inflation factor and family-wise error rate for the MWAS methods with the measured and predicted CTCs fitted as covariates in simulation scenario 2. (a) Mean $\lambda$ value from 1000 simulation replicates given a specific $R^2_{\text{CTCs}}$ value for a method with an error bar representing +/- the SE of the mean. (b) False positive rate. CTCs: linear regression with the measured CTCs fitted as covariates. "preCTCs" or "_preCTCs": analysis with the predicted CTCs fitted as covariates.

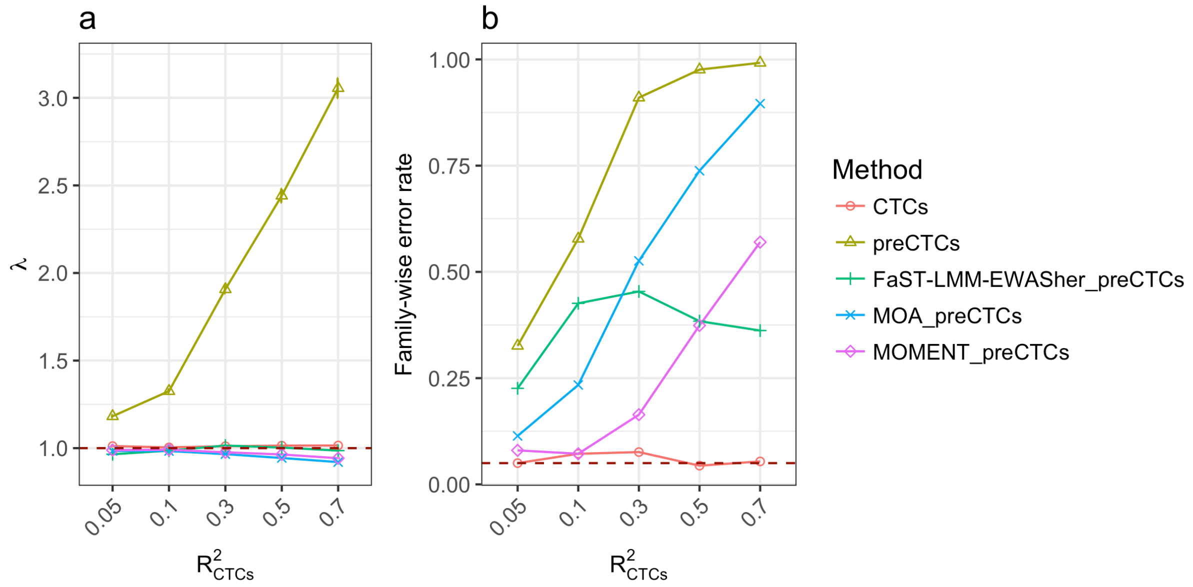


**Supplementary Figure 36** Genomic inflation factor and family-wise error rate for the MWAS methods with the predicted CTCs fitted as covariates in simulation scenario 2 with large $R^2_{\text{CTCs}}$ (ranging from 0.05 to 0.7). (a) Mean $\lambda$ value from 1000 simulation replicates with an error bar representing +/- the SE of the mean. (b) False positive rate. CTCs: linear regression with the measured CTCs fitted as covariates. “preCTCs” or “_preCTCs”: analysis with the predicted CTCs fitted as covariates.

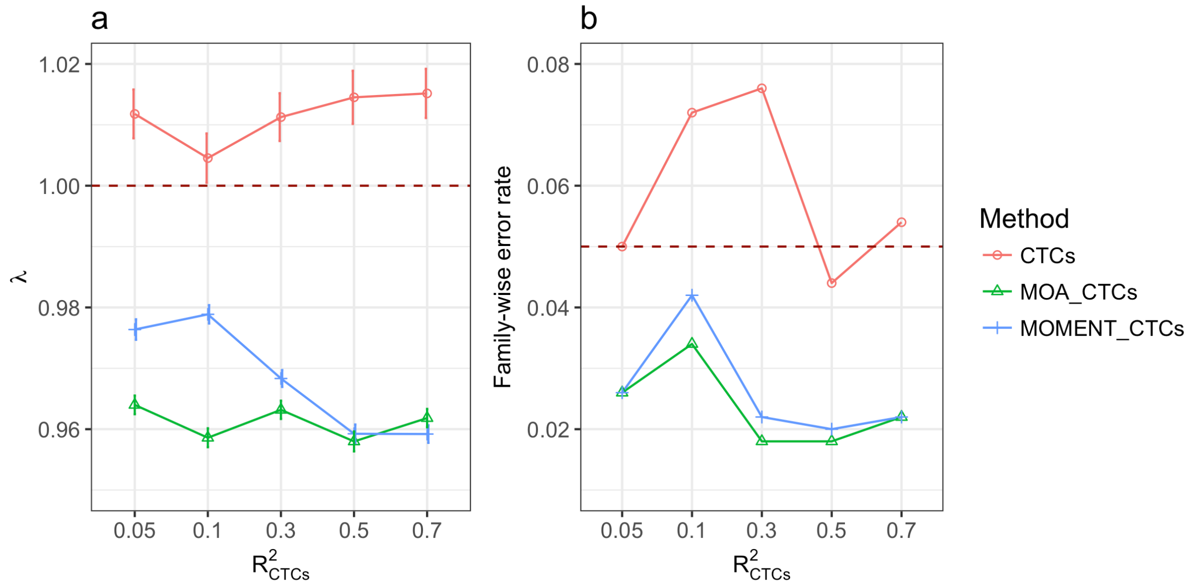


**Supplementary Figure 37** Genomic inflation factor and family-wise error rate for MOA and MOMENT with the measured CTCs fitted as covariates in simulation scenario 2 ($R^2_{\mathrm{CTCs}}$ ranging from 0.05 to 0.7). (a) Mean $\lambda$ value from 1000 simulation replicates with an error bar representing +/- SE of the mean. (b) False positive rate. CTCs: linear regression with the measured CTCs fitted as covariates. MOA_CTCs (or MOMENT_CTCs): MOA (or MOMENT) analysis with the measured CTCs fitted as covariates.

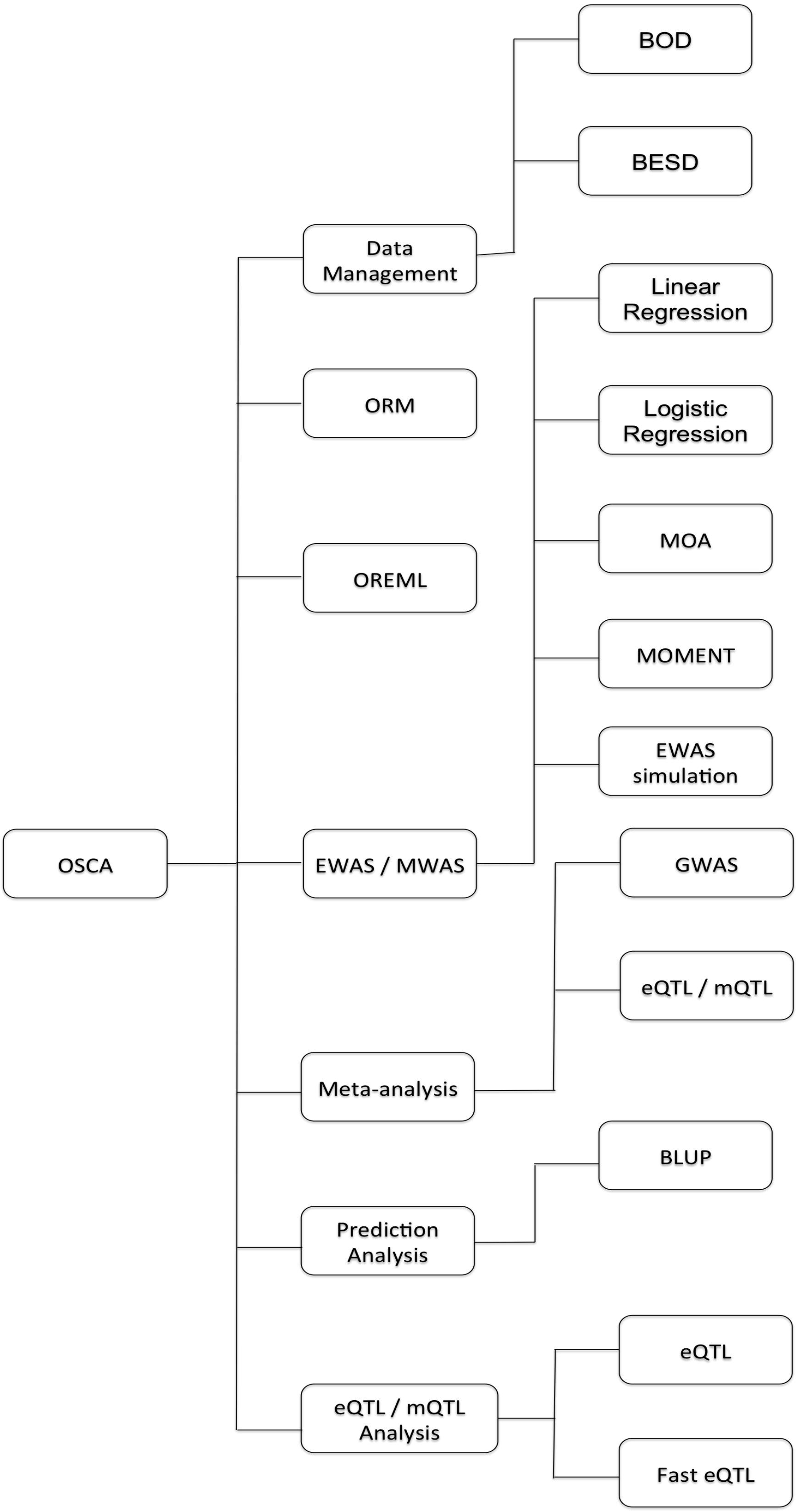


**Supplementary Figure 38** Schematic diagram of OSCA.

**Supplementary Table 1** Summary of the results from all the MWAS methods in simulation scenario 1.

| | Even chromosomes (null) | | | | $\overline{D}$ | s.e.m. | FWER | AUC | s.em. | FPR | s.e.m |
|---|---|---|---|---|---|---|---|---|---|---|---|
| | $\overline{x^2}$ | s.e.m. | $\bar{\lambda}$ | s.e.m. | | | | | | | |
| Unadjusted | 8.58 | 0.66 | 7.67 | 0.532 | 0.405 | 0.013 | 1.00 | 0.63 | 0.0049 | 0.424 | 0.014 |
| CTC | 4.95 | 0.26 | 4.64 | 0.24 | 0.328 | 0.01 | 1.00 | 0.65 | 0.0041 | 0.338 | 0.011 |
| 5PCs | 1.67 | 0.028 | 1.63 | 0.023 | 0.118 | 0.003 | 0.97 | 0.72 | 0.0025 | 0.127 | 0.003 |
| ReFACTor | 1.91 | 0.044 | 1.90 | 0.045 | 0.149 | 0.005 | 0.99 | 0.71 | 0.0030 | 0.153 | 0.004 |
| LFMM2-ridge | 1.42 | 0.010 | 1.41 | 0.009 | 0.083 | 0.002 | 0.92 | 0.74 | 0.0025 | 0.100 | 0.001 |
| LFMM2-lasso | 1.36 | 0.008 | 1.35 | 0.008 | 0.073 | 0.001 | 0.82 | 0.74 | 0.0025 | 0.092 | 0.001 |
| SVA | 1.97 | 0.081 | 1.81 | 0.061 | 0.134 | 0.007 | 0.96 | 0.72 | 0.0031 | 0.147 | 0.006 |
| FaST-LMM-EWASher | 1.25 | 0.020 | 1.24 | 0.019 | 0.054 | 0.003 | 0.63 | 0.73 | 0.0030 | 0.080 | 0.002 |
| MOA | 1.00 | 0.001 | 1.00 | 0.001 | 0.002 | 4e-4 | 0.10 | 0.75 | 0.0024 | 0.050 | 7.9e-5 |
| MOMENT | 1.01 | 0.001 | 1.01 | 0.001 | 0.004 | 5e-4 | 0.03 | 0.74 | 0.0028 | 0.051 | 1.3e-4 |

$\overline{x^2}$: mean chi-squared value.

s.e.m.: standard error of the mean.

$\bar{\lambda}$: mean genome inflation factor.

$\overline{D}$: Kolmogorov-Smirnov statistic to measure deviation of the p-value distribution from the expected distribution under the null.

**Supplementary Table 2** Summary of the results from all the MWAS methods in simulation scenario 3.

| | Even chromosomes (null) | | | | $\bar{\mathcal{D}}$ | s.e.m. | FWER | AUC | s.em. | FPR | s.e.m |
|---|---|---|---|---|---|---|---|---|---|---|---|
| | $\overline{x^2}$ | s.e.m. | $\bar{\lambda}$ | s.e.m. | | | | | | | |
| Unadjusted | 8.80 | 0.664 | 7.56 | 0.547 | 0.40 | 0.013 | 1.00 | 0.63 | 0.005 | 0.420 | 0.014 |
| CTC | 5.28 | 0.298 | 4.91 | 0.276 | 0.34 | 0.011 | 1.00 | 0.65 | 0.004 | 0.347 | 0.012 |
| 5PCs | 1.64 | 0.020 | 1.61 | 0.017 | 0.12 | 0.003 | 0.96 | 0.72 | 0.003 | 0.124 | 0.002 |
| ReFACTor | 1.92 | 0.041 | 1.92 | 0.044 | 0.15 | 0.005 | 0.99 | 0.71 | 0.003 | 0.155 | 0.004 |
| LFMM2-ridge | 1.42 | 0.010 | 1.40 | 0.009 | 0.08 | 0.002 | 0.94 | 0.73 | 0.003 | 0.100 | 0.001 |
| LFMM2-lasso | 1.35 | 0.009 | 1.34 | 0.008 | 0.07 | 0.001 | 0.80 | 0.73 | 0.003 | 0.092 | 0.001 |
| SVA | 2.03 | 0.092 | 1.83 | 0.066 | 0.14 | 0.007 | 0.96 | 0.73 | 0.003 | 0.150 | 0.007 |
| FaST-LMM-EWASher | 1.22 | 0.020 | 1.21 | 0.020 | 0.05 | 0.004 | 0.53 | 0.73 | 0.003 | 0.076 | 0.002 |
| MOA | 1.00 | 0.001 | 1.00 | 0.001 | 0.002 | 9.1e-5 | 0.11 | 0.75 | 0.003 | 0.050 | 7.2e-5 |
| MOMENT | 1.01 | 0.001 | 1.01 | 0.001 | 0.004 | 2.1e-4 | 0.08 | 0.74 | 0.003 | 0.051 | 1.6e-4 |

$\overline{x^2}$: mean chi-squared value.

s.e.m.: standard error of the mean.

$\bar{\lambda}$: mean genome inflation factor.

$\bar{\mathcal{D}}$: Kolmogorov-Smirnov statistic to measure deviation of the p-value distribution from the expected distribution under the null.

**Supplementary Table 3** Computational complexity, runtime and memory usage of the MWAS methods.

| MWAS Method | Computational complexity | Simulated phenotype | | | | Smoking status |
|---|---|---|---|---|---|---|
| | | Mean runtime (min) | s.e.m. | Mean memory usage (GB) | s.e.m. | Runtime (min) |
| SVA | $O(bmn^2)$ | 343.7 | 0.86 | 34.22 | 0.02 | 555.2 |
| LFMM2-ridge | $O(mn^2 + mnln(K))$ | 4.0 | 0.02 | 9.32 | 0.03 | 3.2 |
| LFMM2-lasso | $O(t(mn^2 + mnln(K)))$ | 144.0 | 1.96 | 9.52 | 0.12 | 413.1 |
| ReFACTor | $O(mn^2 + mnk + tn^2)$ | 7.8 | 0.01 | 17.3 | 0.01 | 9.5 |
| FaST-LMM-EWASher | $O(mn^2 + k(ln^2 + n^3 + mn))$ | 249.9 | 7.67 | 15.8 | 0.01 | 162.9 |
| MOA (exact) | $O(mn^2 + mn^3)$ | 12352.8 | 393.94 | 4.7 | 0.015 | 15485.3 |
| MOA (approximate) | $O(mn^2 + n^3 + mn)$ | 5.7 | 0.15 | 4.7 | 7e-5 | 4.0 |
| MOMENT (exact) | $O(m(mn^2 + n^3))$ | 20840.84 | 860.88 | 4.7 | 0.009 | 18975.8 |
| MOMENT (approximate) | $O(k(mn^2 + n^3) + n^3 + mn)$ | 16.6 | 0.15 | 4.7 | 5e-3 | 16.5 |

Notations: $m$ is the number of probes; $n$ is the sample size; SVA: $b$ denotes the number of repeats used to obtain null statistics; LFMM2-ridge: $K$ denotes the number of latent factors; LFMM2-lasso: $t$ denotes the iteration times; ReFACTor: $k$ denotes the number of cell types and $t$ denotes the number of differential methylated sites; FaST-LMM-EWASher: $k$ denotes the number of fitted PCs ranging from 0 to 10 and $l$ denotes the number of probes selected by the feature selection; MOMENT (approximate): $k$ denotes the number of probes included in the first component.

Simulated phenotype: 228,694 DNAm probes on 1,341 from the LBC cohorts. Smoking status (coded as 0, or 1 for non-smoker or former smoker and current smoker): 228,694 DNAm probes on 1341 from the LBC cohorts. All the analyses were performed on the same computer platform (Intel(R) Xeon(R) CPU E5-2670 v3 @ 2.30GHz) using only one CPU thread. The mean runtime and memory usage were computed from 30 replicates for the exact approach and 100 replicates for the other approaches in simulation scenario 1.

The runtime of SVA for Smoking status includes surrogate variables computing using SVA package and logistic regression by fitting the surrogate variables as covariates using OSCA. The runtime of ReFACTor for Smoking status includes sparse PCs estimation using ReFACTor and logistic regression by fitting the Pcs as covariates using OSCA.

**Supplementary Table 4** Genome-wide significant DNAm probes identified by MOA for three traits in the LBC.

| Trait | CHR | Probe | bp | Gene | *β* | SE | *P* | #Methods |
|---|---|---|---|---|---|---|---|---|
| BMI | 17 | cg11202345 | 76976057 | LGALS3BP | 4.50 | 0.74 | 1.07E-09 | 9 |
| BMI | 22 | cg20496314 | 39759864 | SYNGR1 | 3.78 | 0.69 | 5.29E-08 | 5 |
| Lung Function | 5 | cg05575921 | 373378 | AHRR | 2.76 | 0.28 | 1.69E-22 | 8 |
| Lung Function | 2 | cg05951221 | 233284402 | ALPPL2 | 4.19 | 0.44 | 2.83E-21 | 8 |
| Lung Function | 19 | cg03636183 | 17000585 | F2RL3 | 4.00 | 0.44 | 1.50E-19 | 7 |
| Lung Function | 2 | cg01940273 | 233284934 | ALPPL2 | 4.19 | 0.51 | 2.57E-16 | 7 |
| Lung Function | 6 | cg06126421 | 30720080 | IER3 | 2.67 | 0.35 | 4.27E-14 | 7 |
| Lung Function | 15 | cg00310412 | 74724918 | SEMA7A | 4.86 | 0.75 | 1.16E-10 | 6 |
| Lung Function | 6 | cg15342087 | 30720209 | IER3 | 5.51 | 0.88 | 3.08E-10 | 6 |
| Lung Function | 1 | cg09935388 | 92947588 | GFI1 | 2.18 | 0.35 | 3.34E-10 | 6 |
| Lung Function | 6 | cg24859433 | 30720203 | IER3 | 4.74 | 0.77 | 6.46E-10 | 6 |
| Lung Function | 17 | cg19572487 | 38476024 | RARA | 2.74 | 0.47 | 5.05E-09 | 6 |
| Lung Function | 14 | cg13525276 | 81426012 | TSHR | -2.40 | 0.43 | 2.22E-08 | 6 |
| Lung Function | 11 | cg11660018 | 86510915 | PRSS23 | 3.24 | 0.61 | 1.04E-07 | 4 |
| Lung Function | 11 | cg20886049 | 76493545 | TSKU | 2.93 | 0.56 | 1.46E-07 | 4 |
| Lung Function | 17 | cg18181703 | 76354621 | SOCS3 | 3.15 | 0.60 | 1.91E-07 | 6 |
| Walk Speed | 20 | cg05232694 | 48809539 | CEBPB-AS1 | -1.66 | 0.30 | 3.58E-08 | 5 |

#Methods: number of methods (out of a total of 9) by which the probe was identified at a genome-wide significance level.

**Supplementary Table 5** Number of genome-wide significant DNAm probes identified by different MWAS methods with smoking status fitted as a covariate for lung function in the LBC.

| | Lung Function | |
|---|---|---|
| | Lambda | Number of significant associations |
| Unadj | 0.891 | 1 |
| 5PCs | 1.064 | 1 |
| ReFACTor | 1.049 | 3 |
| FaST-LMM-EWASher | 0.996 | 1 |
| MOA | 0.893 | 1 |
| CTCs | 0.945 | 0 |

Unadj: linear regression with smoking status fitted as a covariate. 5PCs: linear regression with the top 5 PCs and smoking status fitted as covariates. ReFACTor: linear regression with 5 sparse PCs from ReFACTor and smoking status fitted as covariates. FaST-LMM-EWASher: FaST-LMM-EWASher with smoking status fitted as a covariate. MOA: MOA with smoking status fitted as a covariate. CTCs: linear regression with smoking status and 5 cell types fitted as covariates.

**Supplementary Table 6** Accuracy of predicting smoking status by DNAm probes in the LBC cohorts.

| | 3 probes identified by MOMENT | 89 probes identified by MOA | 149 probes identified by linear regression |
|---|---|---|---|
| AUC | 0.895 | 0.843 | 0.801 |

Note: 1) The LBC1936 cohort (*n* = 906) was used as the training set and the LBC1921 cohort (*n* = 436) as used as the test set. 2) The numbers of probes detected by the three methods in the training set were smaller than those in the whole set because of the smaller sample size of the training set than the whole set. 3) The area under the ROC curve (AUC) for the categorical phenotype (coded as 0, 1 or 2) was computed using the R package pROC [5].

**Supplementary Table 7** Estimated proportion of variance in a phenotype captured by all DNAm probes by OSCA-OREML from 500 simulations based on the LBC data.

| | NULL | | uni (0.1) | | multi50 (0.2) | | multi500 (0.5) | |
|---|---|---|---|---|---|---|---|---|
| | Mean $\hat{\rho}^2$ | s.e.m. | Mean $\hat{\rho}^2$ | s.e.m. | Mean $\hat{\rho}^2$ | s.e.m. | Mean $\hat{\rho}^2$ | s.e.m. |
| Probes standardised | 0.0051 | 0.0008 | 0.098 | 0.0023 | 0.198 | 0.0022 | 0.501 | 0.0024 |
| Probes centralised | 0.0048 | 0.0009 | 0.106 | 0.0024 | 0.210 | 0.0023 | 0.511 | 0.0024 |

Probes standardised: each probe is standardised across all individuals to compute the ORM.

Probes centralised: each probe is centralised across all individuals to compute the ORM.

NULL: the phenotype was simulated from a standard normal distribution with no probe effects.

uni (0.1): randomly sample 1 probe as the causal probe to simulate the phenotype with $\rho^2 = 0.1$.

multi50 (0.2): randomly sample 50 probes as the causal probes to simulate the phenotype with $\rho^2 = 0.2$.

multi500 (0.5): randomly sample 500 probes as the causal probes to simulate the phenotype with $\rho^2 = 0.5$.

s.e.m.: standard error of the mean.

**Supplementary Table 8** Proportion of phenotypic variance explained by all the DNA methylation probes estimated by OSCA-OREML for height and BMI in the LBC.

| | Height | | | | BMI | | | |
|---|---|---|---|---|---|---|---|---|
| | $\hat{h}^2_{\mathrm{SNP}}$ | $\hat{\rho}^2_{me}$ | Joint $\hat{h}^2_{\mathrm{SNP}}$ | Joint $\hat{\rho}^2_{me}$ | $\hat{h}^2_{\mathrm{SNP}}$ | $\hat{\rho}^2_{me}$ | Joint $\hat{h}^2_{\mathrm{SNP}}$ | Joint $\hat{\rho}^2_{me}$ |
| Probes standardised | 0.39 (0.25) | -0.0050 (0.0086) | 0.39 (0.25) | -0.0056 (0.0083) | 0.42 (0.25) | 0.065 (0.038) | 0.40 (0.24) | 0.065 (0.038) |
| Probes centralised | | 0.0068 (0.02) | 0.39 (0.25) | 0.0063 (0.02) | | 0.075 (0.044) | 0.39 (0.24) | 0.074 (0.044) |

$\hat{h}^2_{\mathrm{SNP}}$: estimate of the proportion of variance explained by all the SNPs (i.e. SNP-based heritability).

$\hat{\rho}^2_{me}$: estimate of the proportion of variance captured by all the DNAm probes.

Joint $\hat{h}^2_{\mathrm{SNP}}$: proportion of variance explained by all the SNPs estimated when the GRM and MRM are fitted jointly in OSCA-OREML.

Joint $\hat{\rho}^2_{me}$: proportion of variance captured by all the DNAm probes estimated when the GRM and MRM are fitted jointly in OSCA-OREML.

**Supplementary Table 9** Summary descriptive of phenotypes for the unrelated individuals in the LBC.

| | **LBC1936** | **LBC1921** |
|---|---|---|
| Sample size | 906 | 436 |
| Age (years); mean (SD) | 69.60 (0.58) | 79.14 (0.83) |
| Percentage of Female | 49.45% | 60.32% |
| BMI (kg/m2); mean (SD) | 27.75 (4.37) | 26.18 (4.05) |
| Height (cm); mean (SD) | 166.37 (8.96) | 163.00 (9.25) |
| Lung Function (L); mean (SD) | 2.36 (0.69) | 1.87 (0.63) |
| Walk Speed (seconds); mean (SD) | 3.86 (1.22) | 4.75 (1.98) |

**Supplementary Table 10** Summary of the sample sizes used in different analyses in the LBC.

| | **LBC1936 (*n* = 920)** | **LBC1921 (*n* =446)** |
|---|---|---|
| Unrelated individuals | 1,342 | |
| Sample size for Height | 1,338 | |
| Sample size for BMI | 1,337 | |
| Sample size for lung function | 1,336 | |
| Sample size for walk speed | 1,332 | |
| Number of individuals with CTC data | 1,319 | |

**Supplementary Table 11** Associations of the 5 cell types ($x$) with the 4 traits ($y$) in the LBC.

| | BMI | | | | Height | | | | Lung Function | | | | Walk Speed | | | |
|---|---|---|---|---|---|---|---|---|---|---|---|---|---|---|---|---|
| | $\beta$ | SE | Adjusted $R^2$ | $n$ | $\beta$ | SE | Adjusted $R^2$ | $n$ | $\beta$ | SE | Adjusted $R^2$ | $n$ | $\beta$ | SE | Adjusted $R^2$ | $n$ |
| Neutrophils | -0.032 | 0.02 | 0.016 | 1315 | 0.009 | 0.02 | 0.0011 | 1316 | -0.059 | 0.02 | 0.047 | 1314 | 0.034 | 0.02 | 0.0139 | 1310 |
| Lymphocytes | 0.027 | 0.02 | | | -0.030 | 0.02 | | | -0.032 | 0.02 | | | -0.016 | 0.02 | | |
| Monocytes | 0.403 | 0.16 | | | -0.224 | 0.16 | | | -0.657 | 0.16 | | | 0.46 | 0.16 | | |
| Eosinophils | 0.276 | 0.19 | | | -0.206 | 0.19 | | | -0.489 | 0.19 | | | 0.27 | 0.19 | | |
| Basophils | 2.31 | 0.72 | | | 0.38 | 0.72 | | | -0.939 | 0.70 | | | 0.58 | 0.72 | | |

**Supplementary Table 12** Descriptive summary of each wave of the LBC data.

| | Total sample size | Female sample size | Male sample size | Mean age | Median age |
|---|---|---|---|---|---|
| LBC21_WAVE1 | 436 | 263 | 173 | 79.1 | 79.2 |
| LBC21_WAVE3 | 174 | 94 | 80 | 86.7 | 86.6 |
| LBC21_WAVE4 | 82 | 44 | 38 | 90.1 | 90.1 |
| LBC36_WAVE1 | 906 | 448 | 458 | 69.6 | 69.6 |
| LBC36_WAVE2 | 801 | 381 | 420 | 72.5 | 72.6 |
| LBC36_WAVE3 | 619 | 296 | 323 | 76.3 | 76.3 |
